# Genetic Code Expansion for Site-Specific Encoding of a Switchable, Intrinsic Fluorophore-Quencher Pair to Monitor Protein Dynamics

**DOI:** 10.64898/2026.06.26.734876

**Authors:** Priyanda Giri, Yarra Venkatesh, Moriah H. Mathis, Christina M. Hurley, Chloe M. Jones, Olivier Ndogo Eteme, Zachary M. Hostetler, Richard B. Cooley, Rahul M. Kohli, Ryan A. Mehl, E. James Petersson

**Author notes:** These authors contributed equally to this work (Priyanda Giri, Yarra Venkatesh). E-mail: E.J.P.

## Abstract

Precisely modifying proteins at multiple sites in their native, folded structures offers unique opportunities to answer molecular and cellular-level biological questions. Here, we present a genetic code expansion strategy for site-specific integration of a fluorophore-quencher pair comprising two non-canonical amino acids—acridonylalanine (Acd) and methyltetrazinyl phenylalanine (Tet) — into a protein expressed in *E. coli*. The Acd and Tet pair requires no post-translational labeling, and quenching can be switched off by biorthogonal or photochemical reactions of Tet for convenient internal control experiments. Mechanistic studies based on Stern–Volmer quenching, fluorescence lifetime measurements, and “proline ruler” peptides established the distance dependence of quenching. As proof-of-concept, we applied this strategy to study: 1) calmodulin, a calcium-sensing protein, 2) RecA, a DNA damage sensor in bacteria, and 3) LexA, a transcriptional repressor whose activation by RecA governs acquired antibiotic resistance in bacteria. Using these proteins, we demonstrate that dual Acd/Tet labeling provides molecular-level insights into protein dynamics, enables high-throughput drug screening, and advances tools for studying protein structure–function relationships.

## Introduction

Small fluorophore-quencher pairs offer a powerful and versatile approach for studying protein folding and dynamics in real time with minimal perturbation.^[1–2]^ Recently, genetic code expansion (GCE) technology has facilitated the incorporation of multiple distinct non-canonical amino acids (ncAAs) into a single protein of interest using mutant aminoacyl tRNA synthetase (RS) and tRNA pairs.^[3]^ Dual encoding of two biorthogonal reactive handles using GCE has been reported for proteins expressed in *E. coli* and in mammalian cells.^[4–7]^ Recently, doubly bio-orthogonal handles containing ncAAs for one-pot preparation of protein conjugates have been developed.^[8]^ Many of these efforts have focused on the incorporation of two reactive ncAAs, followed by labeling through the mutually orthogonal SPAAC (Strain-Promoted Azide-Alkyne Cycloaddition) and IEDDA (Inverse Electron Demand Diels-Alder) reactions. However, these hybrid systems are only partially genetically encoded and often suffer from limitations including heterogeneous labeling, large acceptor size, and spectral bleed-through. Despite these advancements, there remains a significant need for dual encoding GCE systems that can directly encode small fluorescent probes into the protein backbone. Building a dual encoding system would eliminate the need for external labeling, ensure precise stoichiometry, and minimize structural perturbation for monitoring protein conformational dynamics.

We have previously studied protein folding and dynamics using GCE encoding of the intrinsically fluorescent amino acids acridon-2-yl-alanine (Acd, δ, Figure 1), paired with various quenching or Förster resonance energy transfer (FRET) partners through protein ligation or cysteine modification.^[9–13]^ However, these methods are either labor-intensive or require that the protein have no other Cys residues. Here, we use dual encoding to pair Acd with the tetrazine ncAA Tet3.0Me (Tet, τ, R = Me, Figure 1).^[14]^ Tet offers a biorthogonal handle for rapid (10⁶ M⁻¹s⁻¹) IEDDA reactions with strained *trans*-cyclooctenes (sTCOs) that complete in minutes at submicromolar concentrations. IEDDA with Tet could be used to attach a FRET partner for Acd. However, in this case, we will not focus on Tet as a biorthogonal handle, but instead on its ability to act as an intrinsic quencher of Acd fluorescence. While through-bond quenching of fluorophores by tetrazines has been observed in conjugates used in IEDDA reactions for protein labeling and in photo-caging of fluorophores,^[15–17]^ to our knowledge, tetrazines have not been previously used as through-space quenchers. Here, we demonstrate that Acd and Tet can be encoded as a FRET pair and employed for measuring a variety of dynamic processes in proteins.

**Figure 1.**
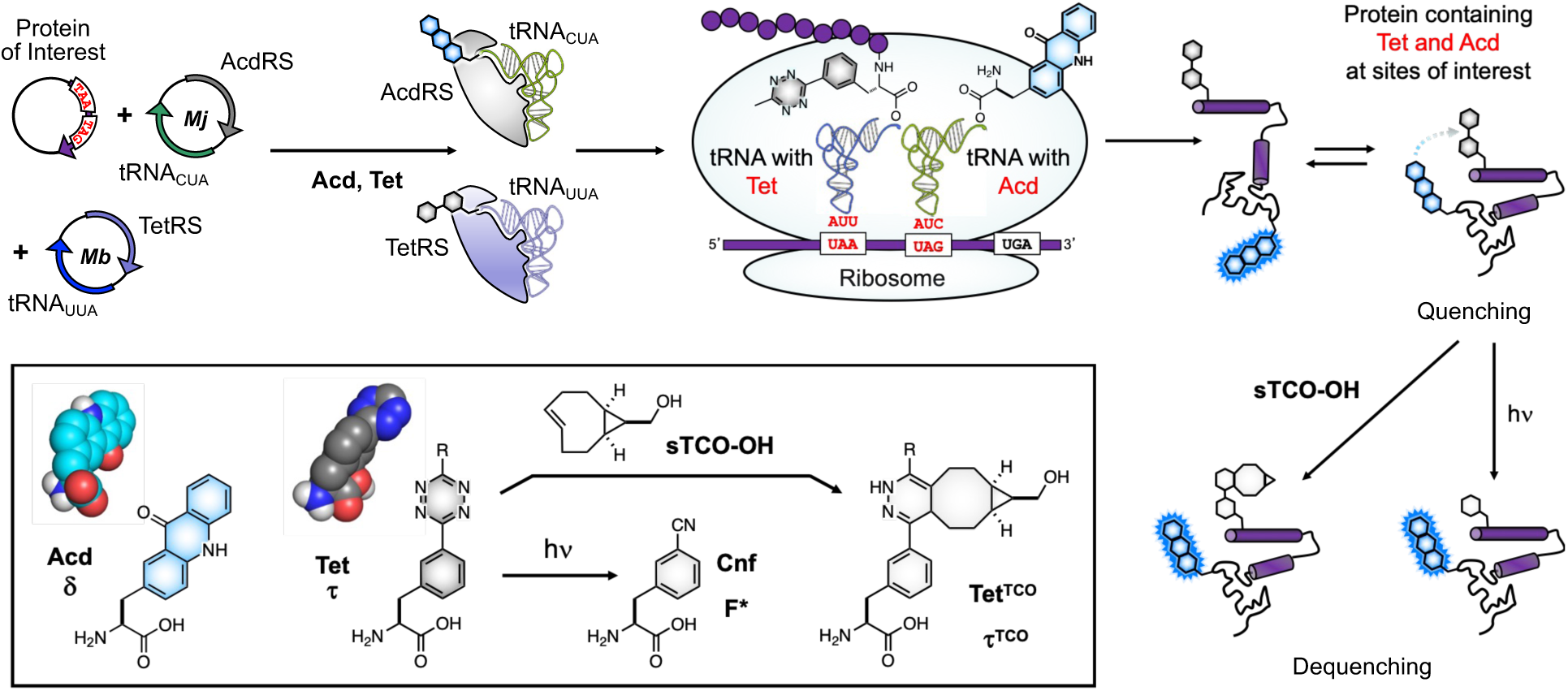
Dual Encoding of Acd and Tet for Protein Dynamics Measurements. Cells are transformed with plasmids encoding TetRS with tRNA_UUA_ and AcdRS with tRNA_CUA_ for incorporation of Tet at a UAA codon and Acd at a UAG codon, and a plasmid encoding the protein of interest with TAA and TAG codons for ncAAs and a TGA codon for translation termination. Ribosomal translation produces a protein labeled with Acd and Tet that can be used to track conformational changes through FRET. De-quenching sTCO-OH or 254 nm irradiation can be used to determine FRET efficiency. Inset: Acd (δ) and Tet (τ, R = Me) structures, space-filling models and de-quenching reactions. Tet IEDDA reaction with sTCO-OH or photolysis to give adduct Tet^TCO^ (τ^TCO^) or Cnf (F*), respectively.

Tet offers valuable properties as a FRET partner for Acd. One challenge of working with FRET pairs is that changes in fluorescence may result from environmental effects on the chromophores, as well as energy transfer between them. Interpreting FRET data often requires generating donor-only or acceptor-only constructs to correct for these effects. Here, we investigate whether Tet quenching can instead be switched off to generate an Acd donor-only control from the Acd/Tet double-labeled protein. We envision that Tet dequenching can occur by disrupting conjugation of the tetrazine π-system through reaction with sTCO (using sTCO-OH as a simple example, Figure 1 inset). We also investigate another simple, biorthogonal way to relieve quenching: the photolysis of the tetrazine ring. Upon UV irradiation, tetrazines are known to undergo photolysis, a reaction that has been used in cleaving peptide linkers and restoring the fluorescence of quenched fluorophores.^[15]^ We test the hypothesis that Tet photolysis to form *m*-cyanophenylalanine (Cnf, F*, Figure 1 inset), N_2_, and CH_3_CN can be used to restore Acd fluorescence with minimal perturbation. Either IEDDA reaction with sTCO-OH or photolysis would provide a convenient way to switch off quenching, obviating the need for separate donor-only control constructs. Thus, Acd/Tet dual encoding provides a fully genetically encoded fluorescence reporting strategy in which quenching can be chemically or photochemically switched off within the same protein sample, enabling straightforward measurements of conformational changes.

## Results and Discussion

### Mechanistic Basis of Tet-mediated Acd Quenching

We began by performing Stern-Volmer titrations with tetrazine phenylacetic acid (Tet*), showing that it quenches Acd 16-fold at 20 mM (Figure 2A, SI Figure S1), which is much more potent quenching than what we have previously observed for the natural amino acids Trp or Tyr (3-fold and 2-fold, respectively).^[11]^ This shows that Acd will be sensitive to proximity-based quenching by Tet even in proteins with Trp or Tyr residues. Tet quenching of Acd has the potential to occur both through an electron transfer mechanism^[17]^ as well as through a FRET mechanism, based on spectral overlap of Acd emission with the weak Tet absorbance centered at 528 nm (Figure 2D). We also performed time-resolved fluorescence lifetime measurements using time-correlated single photon counting (TCSPC), which showed a decrease in Acd lifetime from 14.56 ns to 10.08 ns on the addition of Tet (0–5 mM), suggesting a dynamic quenching mechanism (Figure 2B, SI Table S1).

**Figure 2.**
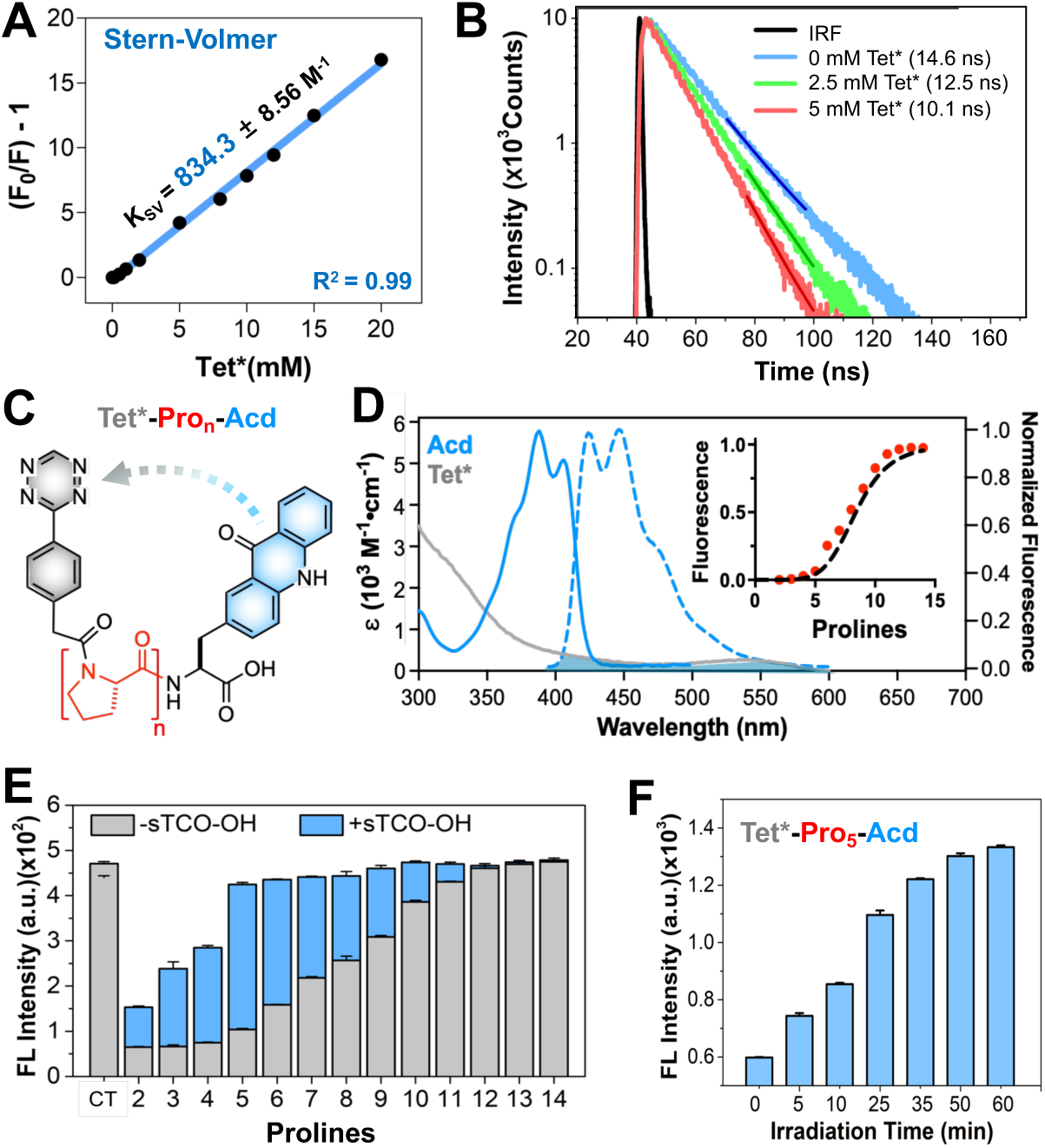
(A) Stern-Volmer quenching of Acd with an increasing concentration of Tet*. (B) Fluorescence lifetime of Acd with an increasing concentration of Tet. (C) Design of the proline ruler (n = 2–14) containing the FRET donor Acd and acceptor Tet* at C- and N-terminus, respectively. (D) Acd/Tet spectra showing Acd absorbance (solid blue line) and emission (dashed blue line), and Tet absorbance (grey solid line) with overlap shaded. Inset: The predicted distance dependence of quenching, using a metric of 3 Å per Pro, is shown as a dashed line in the inset. Circles show experimental fluorescence emission measurements for Tet*-Pro_n_-Acd, n = 2–14, peptides, normalized to the emission of a Leu-Pro_4_-Acd control peptide (CT). (E) Fluorescence emission for the Tet*-Pro_n_-Acd (n = 2-14) peptides and Leu-Pro_4_-Acd control peptide before and after IEDDA reaction with sTCO-OH. (F) Tet*-Pro_5_-Acd irradiation with 254 nm light, showing dequenching by photolysis of Tet*.

To investigate the distance-dependence of Acd quenching by Tet, we synthesized a group of peptides with Tet* at the N-terminus and Acd at the C-terminus, separated by increasing numbers of proline residues (Tet*-Pro_n_-Acd, n = 2 to 14), as well as a Leu-Pro_4_-Acd control peptide (Figure 2C). All peptides were purified by high-performance liquid chromatography (HPLC) and characterized by liquid chromatography mass spectrometry (LC-MS) or matrix-assisted laser desorption ionization mass spectrometry (MALDI-MS); synthesis and characterization are described in the SI (Scheme S1, Tables S2-S4, Figure S2-S3). Measuring the fluorescence intensities of Tet*-Pro_n_-Acd peptides in phosphate buffer at pH 7.0 revealed a sigmoidal distance dependence with a half-maximal change in fluorescence at 24 Å (8 Pro), using the common approximation of 3 Å per Pro residue (Figure 2D inset and SI Figure S4). This result is in perfect agreement with the theoretical distance dependence for FRET based on a Förster radius (R_0_) of 24 Å, determined from spectral overlap calculations (Figure 2D inset and SI Figure S5). However, it is possible that other processes such as electron transfer and Dexter transfer could still contribute to quenching at short range. Nonetheless, Acd/Tet dual labeling of proteins should clearly be able to detect conformational changes over 10-30 Å distances.

### Bioorthogonal and Photochemical Strategies Enable Controlled Modulation of Acd Fluorescence

Next, we used the Tet*-Pro_n_-Acd peptides to test dequenching with the rapid and highly selective IEDDA reaction of Tet with an sTCO, using the simple alcohol, sTCO-OH (Figure 2E). We were gratified to see that a 15-minute reaction with sTCO-OH eliminated Acd quenching for Tet*-Pro_n_-Acd peptides with n ≥ 5 (completion of reactions was confirmed by MALDI-MS and HPLC, SI Figure S6-S7). For Tet*-Pro_n_-Acd peptides, n = 2–4, some residual quenching remained, which may be the result of contact-based quenching and/or residual FRET between Acd and the Tet-sTCO-OH adduct. Indeed, Stern-Volmer analysis indicates that following sTCO-OH reactions, quenching is much less efficient than what is observed for Tet* (SI Figure S1). Thus, the IEDDA reaction of Tet with a simple sTCO molecule should provide a valuable internal control to evaluate the extent of quenching in a FRET experiment. We next investigated whether photolysis could be used to restore Acd fluorescence with minimal perturbation. As a representative example, we photolyzed Tet*-Pro_5_-Acd, shown in Figure 2F. Tet* efficiently underwent photolysis at 254 nm; HPLC and MALDI-MS analysis further confirmed that photolysis resulted in the corresponding Cnf-Pro_n_-Acd peptide (SI Figure S8). Taken together, our peptide data showed that Acd FRET should be a useful tool for examining conformational changes in proteins, with straightforward access to internal controls to roughly quantify FRET efficiency.

### Development of a Dual-Encoding Strategy for an Intrinsic Acd/Tet FRET Pair

To make use of Acd/Tet in the study of protein dynamics, we developed a GCE system for dual encoding in *E. coli* using the mutually orthogonal *M. barkeri* TetRS/tRNA_UUA_ and *M. jannaschii* AcdRS /tRNA_CUA_ pairs for incorporation of Tet at a TAA codon and Acd at a TAG codon, respectively (Figure 1).^[14, 18]^ Our Acd/Tet system represents the first cell-based GCE incorporation of an intrinsic FRET pair. We demonstrate the versatility of our dual encoding system in the study of three different proteins of interest: 1) calmodulin (CaM), a calcium-sensing protein^[19]^; 2) Recombination protein A (RecA), a DNA repair protein;^[20]^ and 3) LexA, a key regulator of the SOS response that enables acquired antibiotic resistance in bacteria.^[21–22]^ These offer tractable targets for a dual encoding system, since we previously used GCE to incorporate single ncAAs, including Acd, into these three proteins, and CaM has been used by others to test dual ncAA GCE.^[5, 11, 13, 23]^

### Site-Specific Incorporation of Acd and Tet into Calmodulin

We began by optimizing the dual incorporation of Acd/Tet pairs in CaM. CaM is a ubiquitous calcium-binding protein that serves as a major calcium sensor in eukaryotic cells, and it is a protein whose dynamics we have previously investigated with a variety of non-perturbing chromophore pairs.^[9, 11, 24]^ The CaM plasmid was modified with TAA to encode Tet in place of Phe_93_ and TAG to encode Acd in place of Leu_113_. Examination of NMR data indicate that this positioning places Acd and Tet within a target range where quenching should be sensitive to Ca^2+^-induced conformational changes (Figure 3A).^[25–26]^ We used electroporation to co-transform the pUltral-TetRS/tRNA_UUA_, pEVOL-AcdRS/tRNA_CUA_, and pTXB1-CaM-TAA_93_TAG_113_ plasmids into *E. coli* BL21 DE3 cells to enable expression of CaMτ_93_δ_113_ (τ = Tet, δ = Acd) fused to a C-terminal GyrA intein containing a His_6_ tag (SI Figure S9). This fusion construct facilitates the elimination of truncated proteins from early termination, as the intein with its His_6_ tag is only produced for the full-length protein and can be cleaved tracelessly using 2-mercaptoethanol.^[27]^

**Figure 3.**
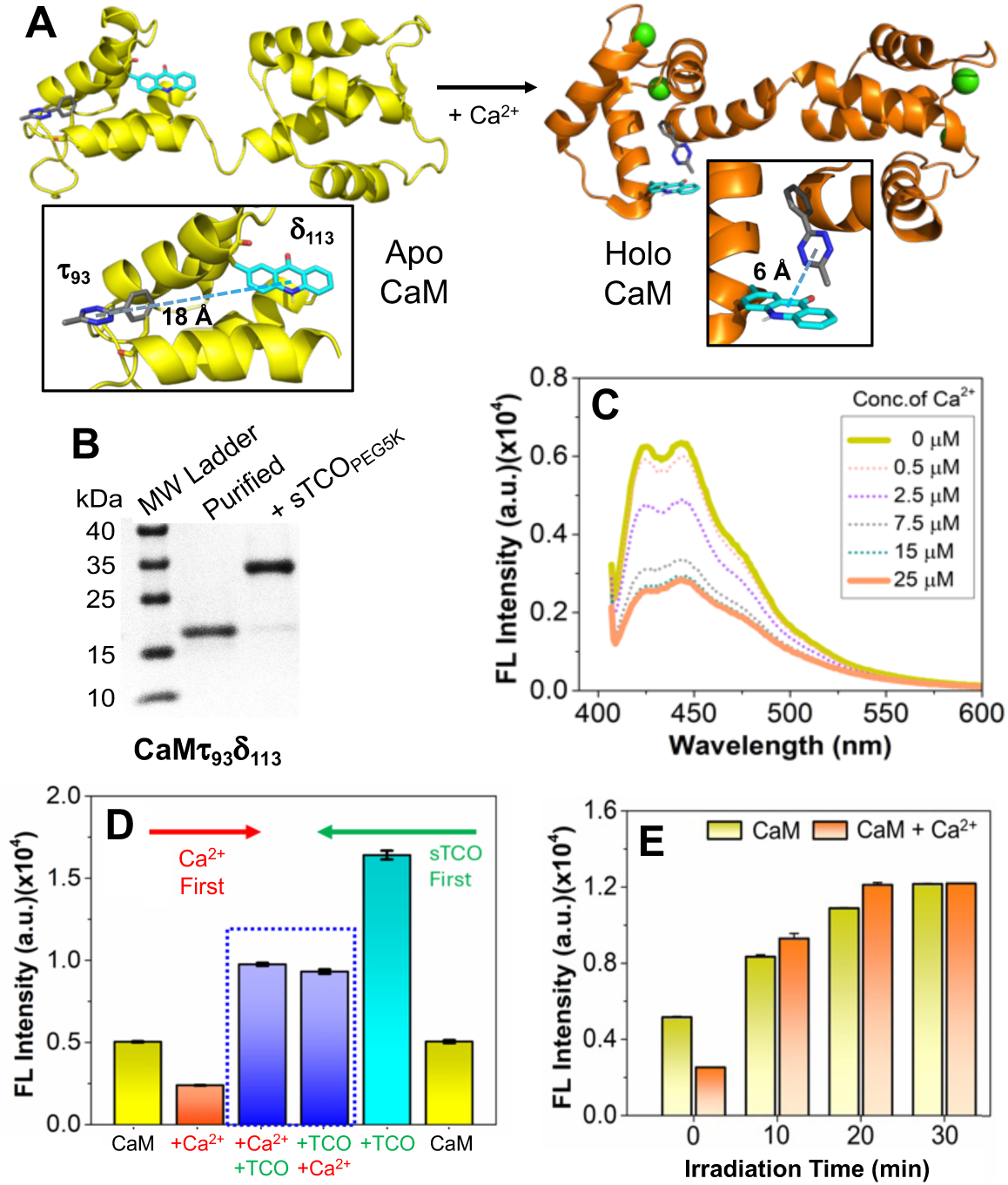
CaM Ca²⁺-Induced Conformational Change. A) CaM structure showing the distance between residues 93 and 113 in the apo (PDB ID 1cfd)^[25]^ and holo states (PDB ID 1x02). (B) Gel analysis of purified CaMτ_93_δ_113_ and confirmation of Tet incorporation via reaction with sTCO_PEG5K_. (C) CaMτ_93_δ_113_ fluorescence spectra during titration with Ca²⁺. (D) Validation of conformational change by varying the order of addition of sTCO-OH and Ca²⁺. (E) Recovery of Acd fluorescence after photolysis of Tet under 254 nm irradiation. Acd excitation/emission wavelengths were 385/450 nm.

Protein production efficiency was monitored *via* gel analysis after lysis, purification, and intein cleavage. At 18 °C, no significant protein production was observed. Increasing the temperature to 25 °C and 30 °C improved protein yields but resulted in significant premature intein cleavage at 30 °C, making 25 °C optimal (SI Figure S9). We purified the full-length protein using Ni affinity chromatography followed by anion exchange chromatography on fast protein liquid chromatography (FPLC), confirming Tet/Acd incorporation *via* a gel shift assay with sTCO_PEG5K_ (an sTCO with a 5 kDa polyethylene glycol chain), fluorescence gel images, and MALDI-MS analysis (Figure 3B and SI Figure S10). Notably, MALDI-MS analysis revealed three peaks corresponding to the full-length Tet/Acd-labeled protein (+Met), an N-terminal methionine-cleaved form (-Met, a known modification due to cleavage by methionine aminopeptidase),^[11]^ and a cysteine adduct formed between Tet and cellular cysteine during expression. These constructs were successfully separated by analytical HPLC and characterized by MALDI-MS (SI Figure S10).

After purifying CaMτ_93_δ_113_ (+Met/-Met), we demonstrated the utility of the Acd/Tet pair by monitoring CaM conformational changes upon Ca²⁺ binding. As shown in Figure 3C and SI Figure S26, the fluorescence intensity decreases by 2.1-fold at saturating Ca²⁺ concentrations. We next validated that the decrease in Acd fluorescence intensity is due to quenching by Tet, and not changes in the Acd environment. As demonstrated earlier, reacting Tet with sTCO-OH *via* an IEDDA reaction to form Tet^TCO^ should break tetrazine conjugation and relieve quenching. To test this, we varied the order of the reactions, either adding Ca²⁺ first and then performing the IEDDA reaction, or starting with the IEDDA reaction followed by the addition of Ca²⁺. Both orders of addition resulted in 1.9-fold increases in fluorescence relative to the starting state (Figure 3D).

Separately, we examined the other mode for Tet de-quenching. Following irradiation with 254 nm light, Tet efficiently underwent photolysis to form Cnf, restoring the Acd fluorescence of the CaM dual construct both in the presence and absence of Ca²⁺ (Figure 3E). Notably, this photolysis strategy achieved maximum fluorescence recovery of Acd (4.8-fold relative to the unirradiated, Ca^2+^-bound state), surpassing the sTCO-OH dequenching reaction (4.1-fold). We also attempted photolysis of Tet at 365 nm; however, no photolysis was observed even after 100 minutes of irradiation (SI Figure S27).

Finally, we asked if observed quenching measurements aligned with estimations we might generate from our data for the Tet*-Pro_n_-Acd peptides. We therefore made crude models of Acd_113_ and Tet_93_ in the apo (PDB ID 1cfd)^[25]^ and holo (PDB ID 1x02)^[26]^ states by alignment with the Phe_93_ and Leu_113_ sidechains. The 2.4-fold quenching in the apo state and the ∼4.5-fold quenching in the holo state are consistent with the levels of quenching expected based on the separation of the centers of Acd_113_ and Tet_93_ (18 Å apo, 6 Å holo) in these models (Figure 3A). Collectively, these experiments on CaM – a structurally well-defined, dynamic protein – show that we can monitor intramolecular conformational change with experimental validation from Acd de-quenching, either through reaction of Tet with sTCO-OH to form Tet^TCO^ or photolysis to form Cnf.

### Site-Specific Incorporation of Acd and Tet into RecA

To move from studying intramolecular dynamics to complex oligomeric assembly of proteins, we applied the dual incorporation approach to RecA, a key bacterial protein involved in homologous recombination, SOS response activation, and DNA strand exchange.^[20, 28–29]^ RecA can serve as a DNA damage sensor by forming filamentous assemblies along single-stranded DNA (ssDNA), referred to as RecA*, in an ATP-dependent manner (Figure 4C). To design a RecA construct in which Tet quenching of Acd could be used to monitor RecA* formation, we focused on two previously-identified ncAA tolerant positions,^37^ Arg_33_ and Ile_102_, that are distant (∼40 Å) in the monomeric RecA subunit but are close to one another in adjacent subunits (∼6 Å) of the RecA* polymer (Figure 4A). We dual encoded Tet and Acd at positions Arg_33_ and Ile_102_, respectively, and the RecAτ_33_δ_102_ construct was expressed and purified by FPLC using a heparin column. Successfully dual-labeled RecA was characterized by gel and MALDI-MS analysis (Figure 4B and SI Figure S11 and S24). The sTCO_PEG5K_ reaction proceeded to approximately 70% completion, indicating either misincorporation or tetrazine reduction (see LexA section below for further discussion).^[30]^

**Figure 4.**
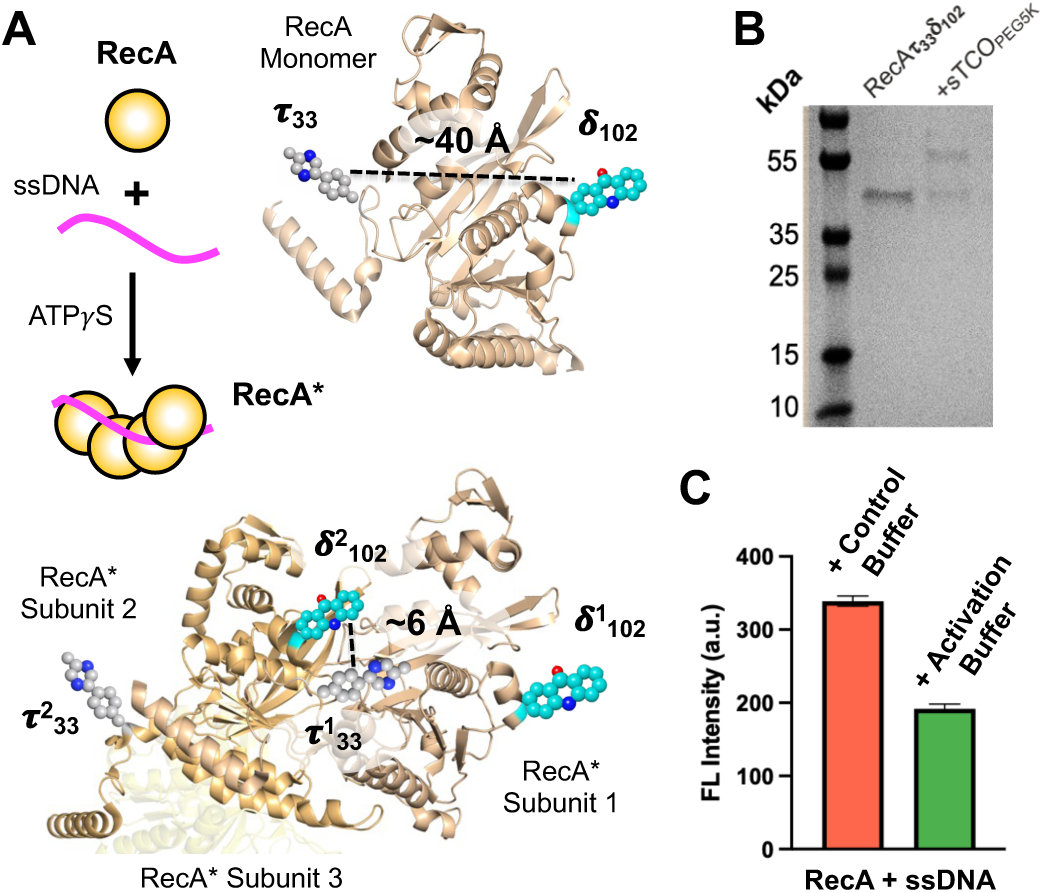
Acd/Tet Quenching Detects RecA Filamentation. (A) Scheme for RecA filamentation to form RecA* with RecA monomer and two RecA* subunit structures showing distances between Tet_33_ and Acd_102_ (PDB ID 2reb and 8trg, respectively). (B) Gel analysis of purified RecAτ_33_δ_102_, and confirmation of Tet incorporation via reaction with sTCO_PEG5K_. (C) Quenching of Acd fluorescence on RecA* formation in Activation Buffer with ssDNA and ATPγS. Control Buffer lacks ATPγS.

In spite of incomplete Tet labeling, we wished to determine whether our design could indeed detect RecA* formation, so we mixed RecAτ_33_δ_102_ with 18-mer (GGT)_6_ ssDNA in the presence of either a control buffer or an activation buffer. Consistent with the above-mentioned model, in the presence of activation buffer (containing non-hydrolyzable ATP analog ATPγS), the RecA filament mixture exhibited a significant reduction in fluorescence compared to the control mixture without ATPγS (Figure 4C).

### A Real-Time FRET Biosensor for Monitoring LexA Cleavage Kinetics

Moving from studying oligomerization of a single protein to multi-protein complexes, we next chose to investigate the interaction of LexA and RecA, which is central to the SOS response, the bacterial DNA damage response pathway underlying adaptation to environmental stress and antibiotic resistance.^[22]^ SOS genes are normally kept silent by the repressor LexA and encode genes, including error-prone DNA polymerases, for bypassing DNA lesions. When RecA* filaments form on ssDNA, they can induce autoproteolysis of LexA, activating the SOS response (Figure 5A). Little et al. studied RecA*’s role in LexA autoproteolysis, revealing that RecA* binding aligns the catalytic residues (Ser_119_ and Lys_156_) with the scissile bond (Ala_84_-Gly_85_), enabling self-cleavage of LexA (LexA**).^[31]^ However, traditional methods of monitoring LexA cleavage using gel analysis require additional steps, such as running a gel after the cleavage assay and quantifying the gel bands for kinetic measurements, which makes them labor-intensive and time-consuming compared to an Acd-Tet FRET-based biosensor that can potentially monitor the kinetics in real time.

**Figure 5.**
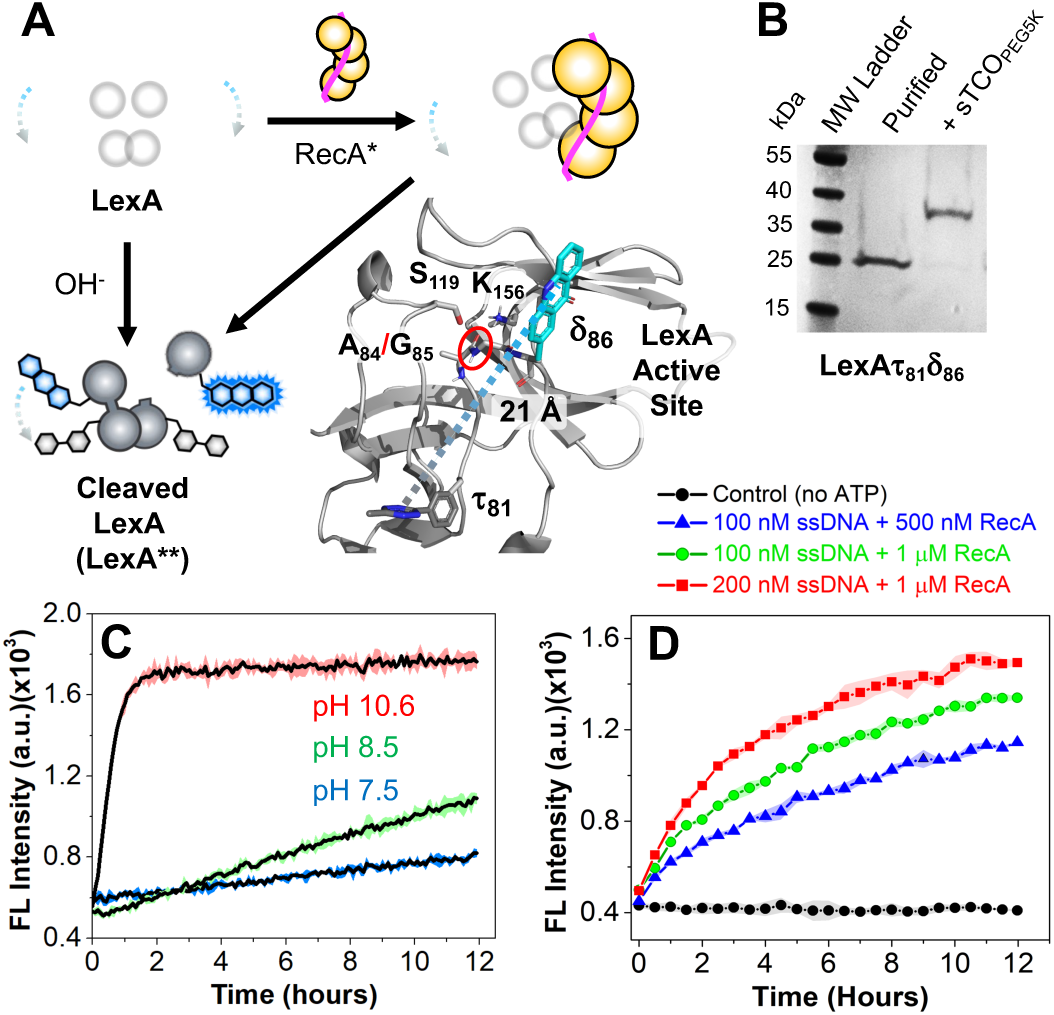
LexA Cleavage. (A) Scheme for pH- or RecA*-dependent induction of LexA cleavage to form LexA** and design of LexAτ_81_δ_86_ with Acd and Tet flanking the A_84_-G_85_ cleavage site (red circle), as shown in a model of the protease domain from one monomer of LexA (PDB ID 8trg). (B) Gel analysis of expressed LexAτ_81_δ_86_ and confirmation of Tet incorporation by reaction with sTCO_PEG5K_. (C) pH-mediated LexA cleavage monitored by fluorescence activation. (D) Monitoring RecA*-induced LexA cleavage by fluorescence activation. Acd excitation/emission wavelengths were 385/450 nm.

To that end, we used our dual encoding strategy to make fluorescence sensors by strategically introducing Acd and Tet modifications at different positions within LexA. We modified LexA at positions Arg_81_ and Glu_86_ for Tet and Acd, respectively, as these positions are tolerant to mutagenesis and/or Acd incorporation based on prior studies and flank the scissile bond (Ala_84_-Gly_85_), offering the promise that LexA autoproteolysis could result in relief of Acd quenching.^[32–33]^

The LexAτ_81_δ_86_ construct was successfully expressed, purified by size exclusion chromatography on FPLC, and characterized using gel and MALDI-MS analysis (Figure 5B and SI Figure S13 and S25). In this case, the sTCO_PEG5k_ reaction confirmed full Tet incorporation (Figure 5B). Some FPLC fractions contained partially reduced Tet (see SI Figure S22 for details on the effect of Tet oxidation/reduction states on labeling), which could be efficiently re-oxidized to the reactive form using horseradish peroxidase (HRP), as demonstrated by the sTCO_PEG5K_ gel shift assay.

We first monitored cleavage kinetics using the Acd/Tet pair under different pH conditions, as alkaline conditions are known to promote RecA*-independent LexA self-cleavage (Figure 5A). At pH 10.6, we observed high fluorescence activation compared to pH 7.5 or 8.5 (Figure 5C) indicating faster cleavage kinetics, consistent with previous reports and independent gel analysis (SI Figure S28).^[34]^

Having validated that our construct can act as a reporter of autoproteolysis, we next monitored RecA*-induced LexA cleavage using dual-labeled LexAτ_81_δ_86_ (Figure 5D). A mixture of 25 µM LexAτ_81_δ_86_ and activated RecA* (varying concentrations of WT-RecA and ssDNA) was analyzed over 12 h. Significant time-dependent fluorescence changes were observed, while no changes occurred in the absence of the RecA* co-factor ATPγS. Doubling either RecA (1 µM) or ssDNA (200 nM) concentrations resulted in faster fluorescence activation, demonstrating an expected concentration dependence for the apparent rate of RecA*-induced LexA cleavage. Quantitative gel analysis corroborated these fluorescence-based cleavage kinetics (SI Figure S28).

### Probing RecA-Induced Cleavage Loop Dynamics in LexA Using Acd/Tet FRET

Beyond acting as a reporter of the interaction between LexA and RecA, we posited that our dual encoded pair could reveal otherwise unobservable mechanistic steps during activation of the SOS response. Specifically, as identified by Luo *et al*., LexA conformational dynamics are thought to play a critical role in activation.^[35]^ LexA is a dimeric protein in which each monomer can exist in either a cleavable or a non-cleavable state. Using X-ray crystallography, the LexA “Quad” mutant (QM = L_89_P, Q_92_W, E_152_A, K_156_A) has been determined to be trapped in the cleavable conformation without undergoing the chemical steps of cleavage due to a protease-inactivating K_156_A mutation.^[35]^ In spite of this structural insight, the dynamics of this process have remained elusive. Conformational interconversion to the cleavable state (which we refer to as LexA*) would be required prior to LexA auto-proteolysis; however, the role of RecA* in promoting this conformational transition remains unclear, because previous methods could not probe into the mode of interaction to differentiate RecA* preferentially binding LexA* (conformational selection) or RecA* binding and then inducing a transition to LexA* (induced conformational change). The potential intermediates populated in these pathways are shown in Figure 6A.

**Figure 6.**
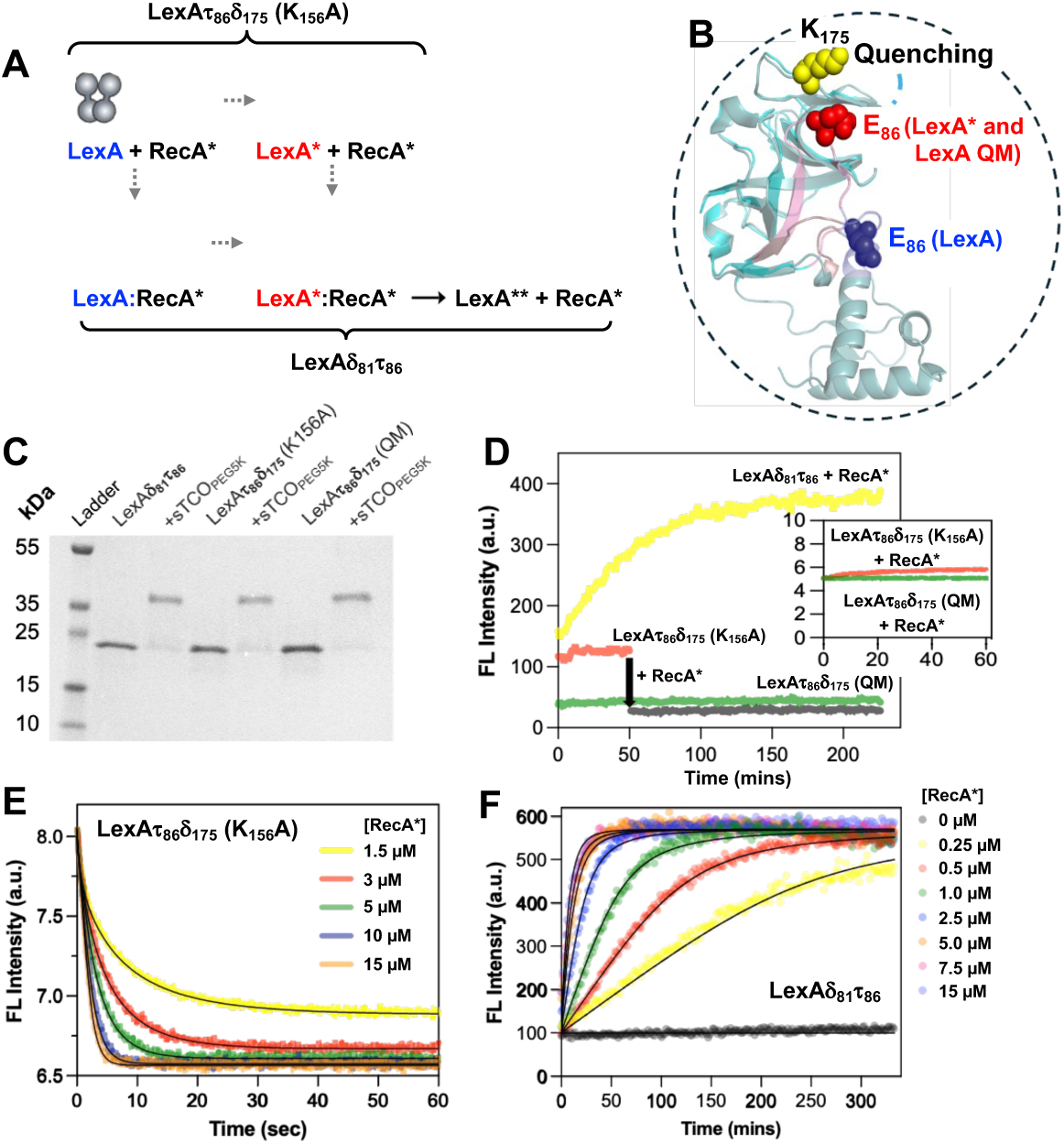
Monitoring kinetics of LexA loop rearrangement and cleavage. (A) Reaction model used for fitting experimental data. RecA*: activated RecA; LexA*: LexA in cleavable conformation; LexA**: Cleaved LexA. Possible intermediates, indicated with gray dashed arrows, include binding prior to conformational change (LexA:RecA*) and conformational change prior to binding (LexA* + RecA*). LexA*τ*_86_*δ*_175_ (K156A) monitors LexA-to-LexA* conversion. LexAδ_81_τ_86_ monitors entire reaction. (B) Crystal structure of LexA monomer in the non-cleavable state (PDB ID: 1jhh, chain A; blue) superimposed on the CTD LexA structure in the cleavable LexA* state (PDB ID: 1jhe, chain A; red), showing distance between residues 86 and 175 in the different states. (C) Gel analysis of expressed Acd/Tet labeled LexA constructs and confirmation of Tet incorporation by reaction with sTCO_PEG5K_. (D) Comparing changes in fluorescence intensity of LexA*τ*_86_*δ*_175_ (K156A) on addition of activated RecA filaments to that of LexA*τ*_86_*δ*_175_ (QM) and LexAδ_81_τ_86_ on a microplate reader. Inset: Fluorescence quenching effect on LexA*τ*_86_*δ*_175_ (K_156_A) and LexA*τ*_86_*δ*_175_ (QM) in the presence of RecA*. (E) Monitoring LexA cleavage loop rearrangement by fluorescence quenching, with the best-fit curves from three independent concentration series experiments shown as black lines for each concentration. (F) Monitoring kinetics of RecA*-induced LexA cleavage by fluorescence activation on a microplate reader, with the best-fit curves from three independent concentration series experiments shown as black lines for each concentration.

To first isolate the binding step of the LexA-RecA* interaction, we incorporated Acd at position K_175_ in LexA, again in a catalytically inactive construct (K_156_A), so that the autoproteolytic step does not complicate the analysis. Following the overexpression and purification of LexA, successful incorporation of Acd at K_175_ was confirmed through SDS-PAGE fluorescence (SI Figure S21). We rapidly mixed 500 nM LexA*δ*_175_ (K_156_A) with RecA* from concentrations 0.08 *μ*M to 15 *μ*M (nearly 200-fold range) in a stopped-flow instrument and monitored for 60 seconds (SI Figure S29). The apparent association rates for RecA* binding to LexA increased as a function of RecA* concentration, and the binding curves did not approach the same maximum anisotropy values for all the concentrations, consistent with a reversible binding model. Based on reaction simulation fitting for a one-step reversible binding model, we obtained association and dissociation rates of 0.0842 *μ*M^-1^s^-1^ and 0.079 s^-1^, respectively. The calculated apparent dissociation constant K_D_ was 0.94 *μ*M for the LexA:RecA interaction, which is similar to the K_D_ value of 0.79 *μ*M previously reported for LexAδ ^24^

Next, to resolve the mechanism of LexA loop dynamics, we expressed two inactive (K_156_A) Acd/Tet dual-labeled constructs – LexA*τ*_86_*δ*_175_ (K156A) and LexA*τ*_86_*δ*_175_ (QM) – and observed high levels of Tet incorporation (Figure 6C, SI Figure S17-S20). Respective Tet and Acd labeling at the E86 and K_175_ positions was predicted to permit Acd quenching given their close proximity (∼12 Å) in the cleavable state. For LexA*τ*_86_*δ*_175_ (K_156_A), we expected to see a decrease in fluorescence upon mixing with RecA* due to Acd quenching in the cleavable state. However, since LexA*τ*_86_*δ*_175_ (QM) is already trapped in the cleavable state, on RecA* binding, we did not expect to see any change in fluorescence. This is indeed what we observed on the plate reader (Figure 6D), so we turned to stopped flow experiments to resolve the kinetics of this conformational change step.

Unlike cleavable LexA, where dissociation of fragments drives the reaction to completion, the non-cleavable mutant reaches a steady-state conformational equilibrium and does not go to completion. On mixing 1.5 *μ*M of LexAτ_86_δ_175_ (K156A) with varying concentrations of activated RecA* from 1.5 *μ*M to 15 *μ*M in the stopped-flow apparatus, we observed an increase in the amplitude of the fluorescence exponential decay with increasing concentration of RecA* (Figure 6E). Importantly, LexA*τ*_86_*δ*_175_ (K156A) did not show any decrease in fluorescence when mixed with the RecA buffer lacking ATPγS (Figure 6D), and additional control experiments demonstrated that the RecA*-induced change in Acd fluorescence was almost exclusively due to Tet quenching and not a change in Acd environment (Figure S38). The observed LexA*τ*_86_*δ*_175_ (K_156_A) kinetic traces were characterized by a distinct biphasic exponential decay and fit well to a hybrid model including initial binding to one of the LexA conformers followed by loop stabilization (SI Figure S33), with a loop rearrangement rate of 1.36 s^-1^. As an important confirmation of this model, the LexA*τ*_86_*δ*_175_ (QM) had a similar level of fluorescence to RecA*-bound LexA*τ*_86_*δ*_175_ (K_156_A), consistent with pre-population of a cleavable LexA conformation (Figure 6D). Performing fluorescence anisotropy measurements of the LexA*δ*_175_ (QM) with activated RecA* filaments using a stopped-flow apparatus demonstrated nearly 10-fold weaker binding relative to LexA*δ*_175_(K_156_A), largely due to much faster dissociation (SI Figure S31). This is consistent with structural data showing that W_92_ in LexA (QM) occupies part of the RecA* binding site, weakening LexA/RecA* binding.^[36]^ Altogether, these observations support our hypothesis that activated RecA* selectively binds a LexA conformer and then further stabilizes the loop rearrangement to the cleavable state.

### Resolving RecA-Dependent LexA Cleavage Kinetics Using Acd/Tet FRET

In classical gel-based cleavage or endpoint assays, LexA-RecA association, conformational rearrangement of the cleavage loop, and subsequent autoproteolysis are kinetically indistinguishable and integrated into an apparent steady-state parameter, making the process seem concerted. However, using our dual labeling approach, we can potentially resolve the LexA cleavage dynamics in combination with the above studies of association and conformational change kinetics. Therefore, we generated an active LexA construct, LexAδ_81_τ_86_, to monitor binding and cleavage kinetics (Figure 6F and SI Figure S15). Mixing 15 μM LexAδ_81_τ_86_ with different concentrations of activated RecA* (0, 0.25, 0.5, 1, 2.5, 5, 7.5, 15 μM) shows a strong dependence of LexA cleavage on RecA* concentration. In the absence of RecA* (no ATPγS), the intensity remains essentially constant over the entire time course, indicating negligible spontaneous cleavage (Figure 6F).

At higher RecA* concentrations, the intensity converges to similar plateau levels, suggesting saturation for the RecA*-mediated activation process. This convergence implies that at high RecA*concentrations the later step (i.e., cleavage) in the multistep reaction becomes rate-limiting and largely RecA*-independent. Moreover, the rate constant of the loop stabilization step obtained using LexA*τ*_86_*δ*_175_ is a fast step that does not leave a distinct kinetic signature on the timescale of a plate reader experiment, and therefore, the experimental data best fit to a two-step model (SI Figure S37). Based on reaction simulation, we obtained the cleavage rate as 3.04x10^-3^ s^-1^. Collectively, our Acd and Acd/Tet LexA experiments thus support a unified kinetic model in which RecA* first binds rapidly and reversibly to one of two pre-existing LexA conformations, a further rapid and reversible stabilization of the LexA cleavage loop occurs prior to cleavage, and finally a composite step of cleavage and dissociation completes the reaction in a rate-limiting and irreversible manner.

### Application of LexA Acd/Tet for Inhibitor Screening

Dual-labeled Acd/Tet constructs offer many possible applications beyond detailed biophysical studies, such as use in high-throughput screening. As proof of concept, we next used the dual-labeled LexA*δ*_81_*τ*_86_ construct to screen a panel of 20 small molecules for inhibition of RecA*-mediated LexA cleavage. To enable this approach, we developed a 96-well plate–based fluorescence assay (Figure 7A). The small-molecule library was assembled in an exploratory manner and included commercially available protease inhibitors (compounds 9–20), compounds related to previously published inhibitors (3–7), and polycyclic aromatic compounds isolated from plants (1–2) (Figure S41). Initial fluorescence measurements from the plate-reader assay identified compound **2**, a highly substituted acridone natural product, as a primary hit, as it significantly suppressed the fluorescence change relative to the DMSO-treated control (Figure 7B and see SI Figure S39 for end-point SDS-PAGE analysis).

**Figure 7.**
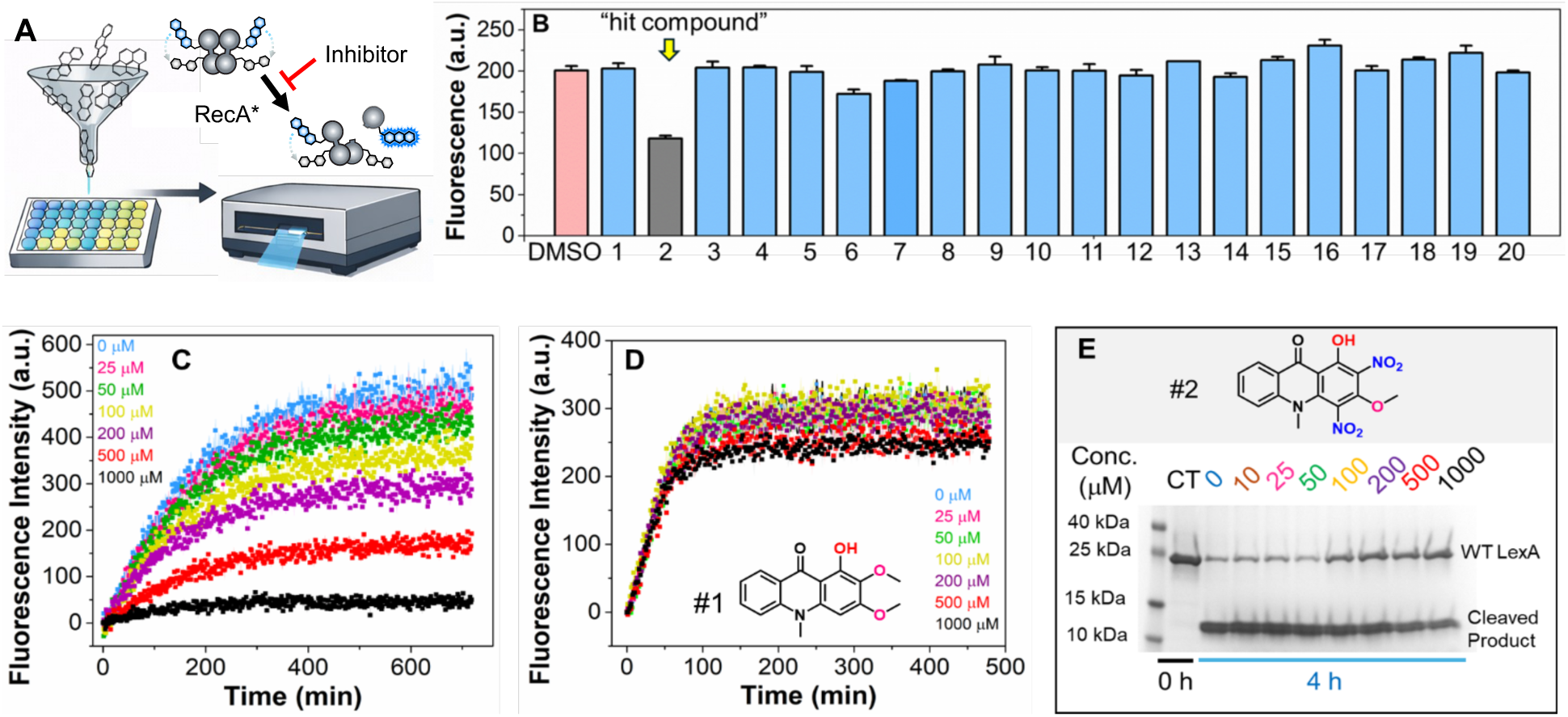
A FRET biosensor for screening LexA inhibitors. (A) Schematic of the 96-well plate–based inhibitor screening workflow using the LexA*δ*_81_*τ*_86_ construct in the presence of RecA*. Fluorescence increase indicates proteolysis, whereas inhibition is marked by the absence of fluorescence increase. (B) Fluorescence intensity from the primary screening of 20 compounds (100 μM each) in a 96-well plate after 8 h incubation of LexA*δ*_81_ *τ*_86_with RecA* at 25 °C. (C) Dose-dependent inhibition of LexA*δ*_81_*τ*_86_ cleavage by hit compound **2** in the presence of RecA* at 25 °C for 12 h. (D) Dose-dependent inhibition of LexA*δ*_81_*τ*_86_ cleavage by negative control compound **1** in the presence of RecA* at 25 °C for 8 h. (E) SDS-PAGE analysis showing dose-dependent inhibition of WT-LexA cleavage by hit compound **2** in the presence of RecA* after incubation at 25 °C for 4 h. Ex/Em for Acd is 385 nm/450 nm.

To further validate this hit, we examined compound **2** in a dose-dependent manner over a concentration range of 0–1000 μM. As shown in Figure 7C, increasing concentrations of **2** led to progressively greater inhibition of LexA cleavage, as evidenced by a corresponding reduction in fluorescence change. To assess whether this effect was structure-specific rather than an assay artifact, we also evaluated the structurally related compound **1**, a natural product analogue differing in substitution, as a negative control. In contrast to **2**, compound **1** showed no detectable inhibition of LexA cleavage across the tested concentration range (Figure 7D), supporting the specificity of compound **2**. The structures of newly isolated natural products **1** and **2** were characterized by ^1^H and ^13^C NMR spectroscopy, high-resolution mass spectrometry (HRMS), and single-crystal X-ray crystallography (SI Figure S42-S45).

Having established compound **2** as a potential LexA inhibitor using the mutant dual-labeled construct, we next sought to validate its activity against WT-LexA. Accordingly, inhibition of RecA*-mediated WT-LexA cleavage was evaluated by SDS–PAGE analysis in a dose-dependent manner (0–1000 μM) after 4 h at 25 °C. As shown in Figure 7E, compound **2** also inhibited cleavage of WT-LexA in a concentration-dependent fashion, confirming that its inhibitory effect is not limited to the engineered fluorescence-reporting construct. Finally, we quantified the inhibitory potency of **2** by determining its IC_50_ values in both assay formats. The IC_50_ values obtained from the dual-labeled LexA fluorescence assay (Figure 7C) and the WT-LexA SDS–PAGE assay (Figure 7E) were 396 μM and 311 μM, respectively (SI Figure S40). We note that by gel analysis, inhibition of WT-LexA reaches about 50% at saturating concentrations, whereas there is no longer any fluorescence change at 1 mM compound **2**. This may indicate that some of the effect in the fluorescence assay is due to quenching by compound **2**, an effect that is more pronounced due to the high IC_50_ of the inhibitor. Nonetheless, these results collectively establish compound **2** as a potential small-molecule inhibitor of RecA*-mediated LexA cleavage and demonstrate the value of Acd/Tet quenching in plate-based screening assays.

## Conclusion

In summary, we present a novel method for monitoring dynamic proteins using fluorescence quenching of Acd by Tet. First, Stern–Volmer and TCSPC lifetime measurements support a quenching mechanism that likely involves a combination of FRET and dynamic, excited-state quenching through shorter-range mechanisms like electron transfer or Dexter energy transfer. Second, peptide “proline rulers” establish the distance dependence of quenching as well as its reversal by either reaction of Tet with sTCO reagents or photolysis. Switching off quenching by sTCO-OH reaction or photolysis provides complementary methods for roughly quantifying FRET efficiency without the need to generate Acd-only control constructs, as we have done in the past.^[13, 24]^ Then, a GCE dual incorporation strategy enables site-specific installation in proteins expressed in *E. coli*. This minimally perturbing approach eliminates the need for post-expression labeling, preserving native protein dynamics and structure for advanced studies. Using orthogonal RS/tRNA pairs, we successfully incorporated these Acd/Tet pairs into three biologically significant proteins: CaM, RecA and LexA. Notably, the Acd/Tet system could be adapted for effective detection of 1) Ca^2+^-induced conformational changes in CaM, 2) RecA assembly on ssDNA to form RecA*, 3) LexA cleavage kinetics and conformational change dynamics, enabling biophysical analysis, and 4) screening of LexA cleavage inhibitors in a simple workflow. Our CaM studies confirm that the de-quenching mechanisms can work in a full-sized protein. The RecA and LexA experiments show how the Acd/Tet pair can probe complex biological mechanisms or enable drug screening assays. Overall, this technology opens new directions for studying dynamic proteins, particularly in cases where labeling with bulkier fluorophores may disrupt protein structure or activity. In our own laboratory, we are already applying this approach both by adapting it to other proteins of interest, such as the Parkinson’s disease associated protein α-synuclein, as well as refining our high-throughput screen for more potent LexA repressor cleavage inhibitors. Finally, parallel efforts seek to understand the mechanism of fluorescence quenching, improve control through chemical and photochemical means, and optimize incorporation of Acd, Tet, and related chromophores to further establish intrinsic probe pairs as robust GCE tools.

## Supporting Information

Peptide and small molecule synthesis and characterization; protein cloning, expression, and characterization; photophysical measurements and calculations; secondary analysis, such as gel imaging; additional references.^[6, 9, 11, 14, 18, 27, 32, 36–47]^

## Acknowledgements

This research was supported by the National Institutes of Health (R01-GM127593 to R.M.K. and E.J.P. supporting LexA/RecA studies, RM1-GM144227 supporting the GCE4All Biomedical Technology Development and Dissemination Center to R.A.M. and R.B.C. for development of the GCE encoding technologies) as well as the National Science Foundation (NSF CHE-2203909 and CHE-1708759 to E.J.P supporting conformational probe development and NSF MCB-2054824 to R.A.M. supporting tetrazine conjugation chemistry development). Instruments supported by the NIH include LC-MS and MALDI-MS instruments (NIH RR-023444 and NIH S10-OD030460). Instruments supported by the NSF include the stopped flow system (NSF CHE-1337449). C.M.H. was supported by the NIH Chemistry Biology Interface Training Program (NIH T32-GM133398) and Z.M.H. by NIH T32-AI118684. C.M.J. was supported by the NIH Structural Biology and Molecular Biophysics Training Program (NIH T32-GM132039).

## Notes

### Competing Interest Statement

The authors have declared no competing interest.

