## Supporting Information for "Genetic Code Expansion for Site-Specific Encoding of a Switchable, Intrinsic Fluorophore-Quencher Pair to Monitor Protein Dynamics"

### Table of Contents


| | |
|---|---|
| General Information | S4 |
| Stern-Volmer Analysis of Tet* Quenching Acd | S5 |
| Fluorescence Lifetime Measurement of Acd in the Presence of Tet* | S6 |
| Synthesis, Purification, and Characterization of Proline Ruler with Tet* and Acd | S6 |
| Proline Ruler FRET Measurements | S12 |
| Dequenching of Proline Ruler Peptides | S15 |
| Cloning of Dual Label Plasmids: CaM, LexA, and RecA | S17 |
| Cloning of Synthetase Plasmids, Sequence Maps, and Primers Used | S19 |
| Reagents and Buffers for Bacterial Culture | S23 |
| Overexpression of CaM, RecA and LexA with Tet and Acd | S26 |
| Overexpression of WT-RecA and WT-LexA | S28 |
| Purification and Characterization of CaM$_{\tau 93 \delta 113}$ | S29 |
| Purification and Characterization of RecA$_{\tau 33 \delta 102}$ | S31 |
| Purification and Characterization of LexA$_{\tau 81 \delta 86}$ | S32 |
| Purification and Characterization of LexA$_{\delta 81 \tau 86}$ | S34 |
| Purification and Characterization of LexA$_{\tau 86 \delta 175}$ (K156A) | S35 |
| Purification and Characterization of LexA$_{\tau 86 \delta 175}$ (QM-L89P, Q92W, E152A, and K156A) | S36 |
| Purification and Characterization of LexA$_{\delta 175}$(K156A) and LexA$d_{175}$(QM) | S37 |
| Oxidation of DihydroTet to Tet by Horseradish Peroxidase (HRP) | S38 |
| Protein Yields and MALDI-MS Analysis of *E. coli* Tet and Acd Expression Systems | S39 |
| Conformational Changes in CaM Upon $Ca^{2+}$ Sensing and Tet Photolysis | S41 |
| LexA Cleavage | S43 |
| RecA Activation Reactions | S45 |
| Fluorescence Anisotropy Measurements | S45 |
| Fluorescence Anisotropy Measurements for LexA$_{\tau 86 \delta 175}$ (QM) | S48 |
| Fluorescence Quenching Measurements (LexA conformational change) | S50 |
| Alternate Models Tested for Conformational Change Step | S53 |
| LexA Cleavage Kinetics Using Fluorescence Intensity Measurement | S58 |
| Control Experiment to Identify the Effect of Surrounding Amino Acids on Acd Fluorescence Quenching | S61 |

| | |
|---|---|
| RecA*-Mediated LexA$\delta_{81}\tau_{86}$ FRET Biosensor Assay | S62 |
| $IC_{50}$ curve for Compound #2 | S63 |
| Chemical Structure of small molecules screened | S63 |
| NMR Spectra of #1 and #2 | S65 |
| Crystal Structure of #1 | S68 |
| Crystal Structure of #2 | S74 |
| Synthetase Sequences | S80 |
| References | S83 |

### General Information

**Materials.** Fluorenylmethoxycarbonyl-Pro-OH was purchased from Novabiochem (currently EMD Millipore, MilliporeSigma; Burlington, MA, USA). Benzotriazol-1-yl-oxy-tris(dimethylamino) phosphonium hexafluorophosphate (BOP) was purchased from Chem-Impex (Wood Dale, IL, USA). Piperidine, 4-(1,2,4,5-Tetrazin-3-yl)benzeneacetic acid (Tet*) and *N,N*-diisopropylethylamine (DIPEA) were purchased from Sigma-Aldrich (St. Louis, MO, USA). All other reagents were purchased from Fisher Scientific (Pittsburg, PA, USA) unless specified otherwise. Milli-Q filtered (18 MΩ) water was used for all solutions (EMD Millipore). *E. coli* BL21 DE3 electrocompetent cells were purchased from MilliporeSigma (CMC0016, 20 x 50 μL). Amicon Ultra centrifugal filter units (3 kDa or 10 kDa MWCO) were purchased from EMD Millipore. The noncanonical amino acids (ncAAs) acridon-2-ylalanine (Acd) and 2-amino-3-(3-(6-methyl-1,2,4,5-tetrazin-3-yl)phenyl)propanoic acid hydrochloride (previously referred to as Tet3.0Me, here simply Tet) were synthesized using previously reported methods.[1-2] DNA sequencing was performed at the University of Pennsylvania sequencing facility unless otherwise noted. The DC protein assay kit was purchased from BioRad (Hercules, CA, USA). Primers were purchased from Integrated DNA Technologies (Coralville, IA, USA). dNTPs and Agarose were purchased from Invitrogen (ThermoFisher, Waltham, MA, USA). Q5 High-Fidelity 2X Master Mix, Gel Green DNA gel stain was purchased from Biotium (Freemont, CA, USA). DNA Clean & Concentrator Kit and Plasmid Miniprep Kit were purchased from Zymo Research (Irvine, CA, USA). Pierce™ Horseradish Peroxidase (Catalog number: 31490) was purchased from Fisher Scientific, USA. BL21(DE3) Electrocompetent Cells (Catalog number: CMC0016) were purchased from MilliporeSigma, USA.

**Instruments.** Proteins were purified with an Agilent 1260 Infinity II Preparative HPLC system and an Agilent 1260 Infinity II Analytical HPLC system (Santa Clara, CA, USA). Protein mass spectrometry data collected Matrix assisted laser desorption ionization (MALDI) mass spectrometry (MS) data were acquired on a Bruker Rapiflex instrument with a standard sinapic acid matrix. Peptide mass spectrometry data were collected with a Bruker MicrofleX (Billerica, MA, USA) MALDI MS using a standard α-cyano-4-hydroxycinnamic acid (CHCA) matrix. Absorbance readings for the DC assay, fluorescence intensity, fluorescence spectral, and fluorescence polarization measurements were made on a Tecan Spark plate reader (Mannedorf, Switzerland). Photolysis of Tet was performed with 7.2 watts USHIO G8T5 Low-Pressure Mercury-Arc lamps that emit radiation peaking at 253.7 nm (UV-C) (Japan). Acd fluorescence was visualized by illuminating the gels using G:Box mini (Syngene), exciting at 365 nm.

### Stern-Volmer Analysis of Tet* Quenching Acd

The experimental procedure for the Stern-Volmer experiments was followed as previously described, with minor modifications.[3] Stocks of 4 mM Acd and 1 M Tet* in 100 mM phosphate buffer, pH 7.0, were used to prepare samples that were uniform in Acd concentration of 100 µM and variable in Tet* concentration (0, 0.1, 0.5, 1, 2, 5, 8, 10, 12, 15, 20 mM). For sTCO-OH modification of Tet*, 50 mM sTCO-OH was added to the above Tet* concentrations and incubated for 15 minutes with shaking (500 rpm) at 37 °C. Fluorescence spectra for each sample were collected at 450 nm with an excitation wavelength of 385 nm in triplicate using Tecan Spark plate reader (5 nm bandwidth, 40 µs integration time, 31000 µm Z-position, 4 sec linear shaking) with a nonsterile Greiner black, flat µClear, 384 well low volume microplate. The data fitted to Equation S1 using GraphPad Prism (version 10.0.0, GraphPad Software, Boston, Massachusetts USA)

$$\frac{F_0}{F} - 1 = K_{SV}[Q] \quad \text{(S1)}$$

$F_0$ is the average fluorescence intensity at 450 nm in the absence of quencher; $F$ is the fluorescence intensity at 450 nm at each Tet* concentration step; $K_{SV}$ is the Stern-Volmer constant in $M^{-1}$; and [Q] is the concentration of the quencher (Tet*) in M (Figure S1).

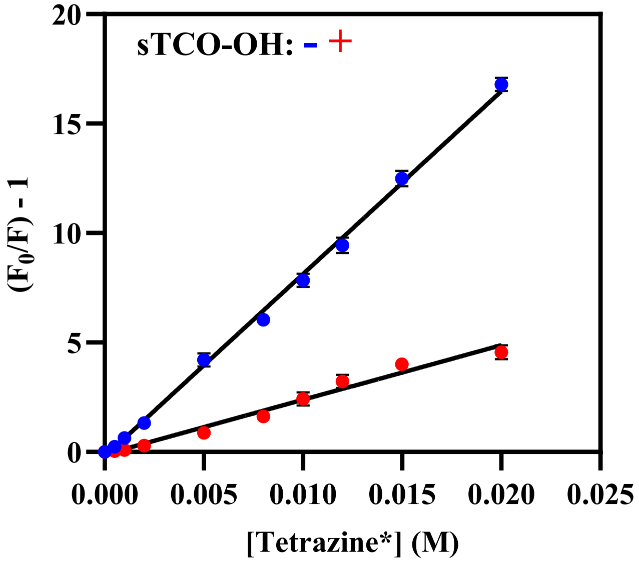


| sTCO-OH | $K_{SV}$ ($M^{-1}$) | $R^2$ |
|---|---|---|
| – | 834.3 ± 8.56 | 0.9967 |
| + | 249.5 ± 7.22 | 0.9859 |

**Figure S1**. Left: Stern-Volmer plots of Tet* quenching Acd. Error bars are calculated from standard error. Right: Stern-Volmer constants and goodness-of-fit values for the fits of Equation S1.

**Fluorescence Lifetime Measurement of Acd in the Presence of Tet***

TCSPC measurements of fluorescence lifetime decays for 5 μM of Acd (in water) in the presence of 0 mM, 2.5 mM, and 5 mM Tet (in water) were collected with Horiba Fluoromax spectrometer using a 340 nm excitation and emission was collected at 485 nm with 20 nm slit widths with a 408 nm long pass filter placed between the sample and the emission monochromator. The instrument response function (IRF) was collected under identical conditions. Data analysis was performed with FluoFit software (PicoQuant GmbH; Berlin, Germany) using an exponential decay model.

**Table S1.** Fluorescence lifetime of Acd at different Tet concentrations.

| Conc.of Tet (mM) | Acd Lifetime (ns) in $H_2O$ | $\chi^2$ |
| --- | --- | --- |
| 0 | 14.56 ± 0.154 | 1.611 |
| 2.5 | 12.49 ± 0.254 | 1.151 |
| 5 | 10.08 ±0.185 | 1.097 |

**Synthesis, Purification, and Characterization of Proline Ruler with Tet* and Acd**

**Peptide Synthesis and Purification.** Tet*-$Pro_n$-Acd, n= 2-14, and Leu-$Pro_4$-Acd were synthesized on a 25 μmol scale on 2-chlorotrityl resin. Tet*-$Pro_n$-Acd, n= 2-8, were synthesized from a common pot of resin (175 μmol); Tet*-$Pro_n$-Acd, n= 9-14, and Leu-$Pro_4$-Acd were synthesized from another common pot of resin (175 μmol). After coupling the appropriate number of proline residues, a 25 μmol portion of resin was removed from the reaction vessel (RV) and transferred to a separate RV for Tet* or Leu coupling. For each synthesis, 2-chlorotrityl chloride resin (100-200 mesh; 1.48 mmol/g; 175 mol) was added to a 6 mL polypropylene syringe (RV). The resin was swollen by incubating it for 45 mins in a (9:1 v/v) mixture of dichloromethane (DCM) and dimethylformamide (DMF) and magnetic stirring. After swelling, the DCM/DMF mixture was removed with vacuum suction, and Fmoc-Acd-OH was coupled to the resin. The amino acid in DMF (2 mL) and DIPEA (4 equiv; 121.8 μL) was added to the RV, and the mixture was allowed to react for two successive 30 min incubations with magnetic stirring. The spent solution was removed with vacuum suction, and the resin beads were cleaned thoroughly with successive DMF and DCM washes. The resin beads were then deprotected by treatment with 20% piperidine in DMF (3 mL) twice for 10 mins each with magnetic stirring. The deprotection solution was drained from the RV, and the beads were rinsed extensively with successive DMF and DCM washes. Subsequent amino acid couplings and deprotections proceeded as described above, with the exception that the Fmoc-Pro-OH and Tet* acid were activated with HBTU (4.8 equiv.) and BOP (1.9 equiv.), respectively, before addition to each reaction. The N-terminal Fmoc group in Leu was removed and acetylated with acetic anhydride (500 μL) and *N*-methyl morpholine (300 μL) in DMF (4.2 mL). The mixture was allowed to react twice for 10 mins each. After the beads were washed extensively with DMF and dried with DCM, peptides were cleaved by 60 mins incubation on a rotisserie with 2 mL of a fresh cleavage cocktail of trifluoracetic acid (TFA) and water (1:1 v/v). The resultant solution was reduced to less than 1 mL by blowing off TFA with argon gas. This solution was then treated with 20 mL of cold ethyl ether to precipitate the peptide. This precipitate was flash frozen with liquid nitrogen and evaporated using a lyophilizer (Labconco; Kansas City, MO, USA).

The crude peptide was diluted in $CH_3CN/H_2O$ (10:90 v/v) and then purified on a Luna®Omega 5 µm PS C18 100 Å, LC semi-preparative column (Phenomenex; Torrance, CA, USA) by HPLC using Gradient 1 for Tet*-$Pro_n$-Acd, n= 2-5 and Leu-$Pro_4$-Acd; Gradient 2 for Tet*-$Pro_n$-Acd, n= 6-8; and Gradient 3 for Tet*-$Pro_n$-Acd, n= 9-14 (Table S2 and S3). MALDI MS and LC-MS were used to confirm peptide identity (Table S4 and Figure S3). Purified peptides were dried on a lyophilizer.

**Scheme S1**. Solid-phase peptide synthesis of Tet*-Pro$_n$-Acd (n = 2-14) on 2-chlorotrityl resin

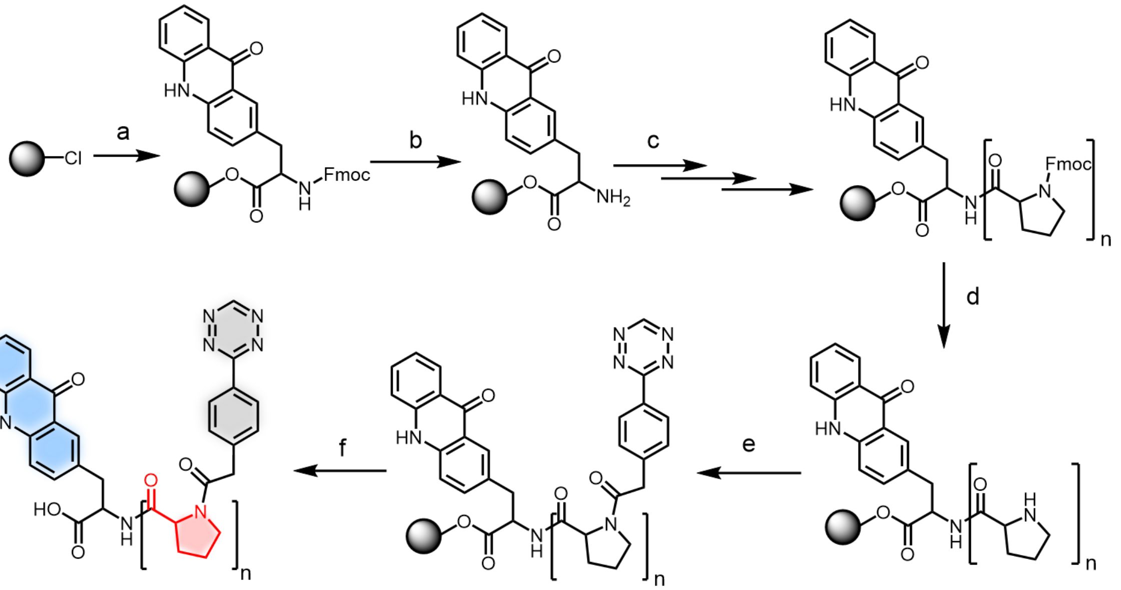


*Reagents and conditions*: (a) Coupling: Fmoc-Acd-OH, HBTU, and DIPEA in dry DMF; (b) Deprotection: 20 % piperidine in DMF; (c) Coupling: Fmoc-Pro-OH, HBTU, and DIPEA in dry DMF; (d) Deprotection: 20 % piperidine in DMF; (e) Coupling: Tet*-COOH or Fmoc-Leu-OH (control), BOP, and DIPEA in dry DMF (f) Cleavage: 50 % Trifluoroacetic acid in water.

**Table S2.** Peptide Purification Methods and Retention Times

| Peptide | Gradient | Retention Time |
|---|---|---|
| Tet*-(Pro)$_2$-Acd | 1 | 34.66 |
| Tet*-(Pro)$_3$-Acd | 1 | 35.92 |
| Tet*-(Pro)$_4$-Acd | 1 | 25.11 |
| Tet*-(Pro)$_5$-Acd | 1 | 26.03 |
| Tet*-(Pro)$_6$-Acd | 2 | 30.74 |
| Tet*-(Pro)$_7$-Acd | 2 | 30.75 |
| Tet*-(Pro)$_8$-Acd | 2 | 31.85 |
| Tet*-(Pro)$_9$-Acd | 3 | 24.10 |
| Tet*-(Pro)$_{10}$-Acd | 3 | 25.33 |
| Tet*-(Pro)$_{11}$-Acd | 3 | 26.79 |
| Tet*-(Pro)$_{12}$-Acd | 3 | 28.12 |
| Tet*-(Pro)$_{13}$-Acd | 3 | 29.69 |
| Tet*-(Pro)$_{14}$-Acd | 3 | 31.23 |
| Leu-(Pro)$_4$-Acd | 1 | 32.19 |

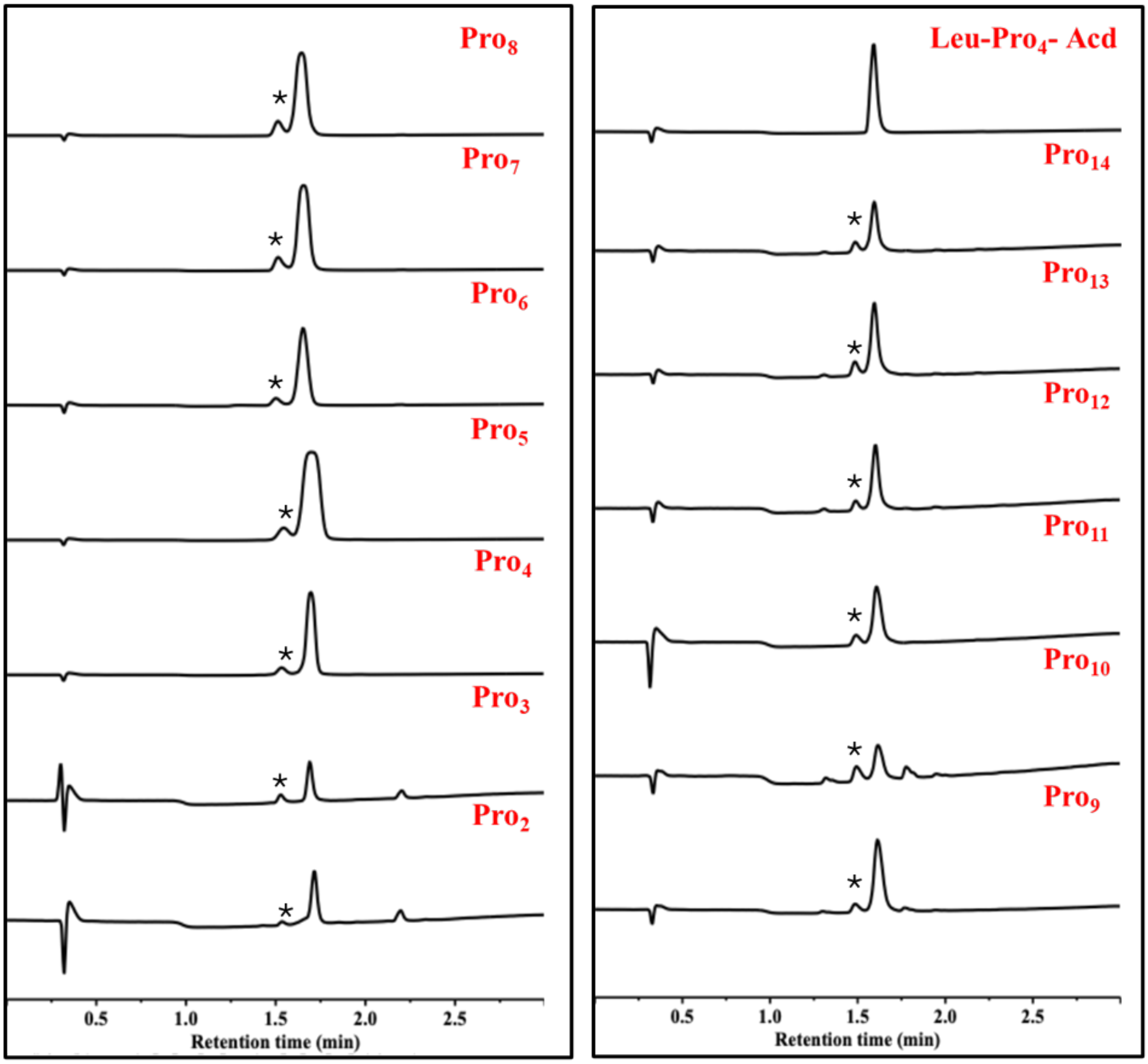


**Figure S2.** LC-MS chromatograms of purified peptides Tet*-Pro$_n$-Acd (n = 2-14) and Leu-Pro$_4$-Acd with * peak corresponding to the reduced form of Tet*.

**Table S3.** HPLC Gradients for Peptide Purification

| | Time (min) | % B | No. | Time (min) | % B | No. | Time (min) | % B |
|---|---|---|---|---|---|---|---|---|
| 1 | 0:00 | 5 | 2 | 0:00 | 5 | 3 | 0:00 | 5 |
| | 5:00 | 5 | | 5:00 | 5 | | 5:00 | 5 |
| | 10:00 | 25 | | 10:00 | 30 | | 10:00 | 40 |
| | 40:00 | 55 | | 40:00 | 60 | | 40:00 | 70 |
| | 45:00 | 100 | | 45:00 | 100 | | 45:00 | 100 |
| | 53:00 | 100 | | 53:00 | 100 | | 53:00 | 100 |
| | 55:00 | 5 | | 55:00 | 5 | | 55:00 | 5 |

**Table S4.** Calculated and Observed Masses of Peptides from LC-MS and MALDI-MS

| Peptide | $[M+H]^+$ | | | | $[M+Na]^+$ | |
|---|---|---|---|---|---|---|
| | Calculated | | Observed | | Calculated | Observed |
| | MALDI-MS | LC-MS | MALDI-MS | LC-MS | MALDI-MS | MALDI-MS |
| Tet*-$(Pro)_2$-Acd | 675.26 | 675.26 | 675.28 | 675.28 | 697.24 | 697.51 |
| Tet*-$(Pro)_3$-Acd | 772.31 | 772.31 | 772.47 | 772.54 | 794.29 | 794.48 |
| Tet*-$(Pro)_4$-Acd | 869.37 | 869.37 | 869.71 | 869.61 | 891.35 | 892.68 |
| Tet*-$(Pro)_5$-Acd | 966.42 | 966.42 | 966.82 | 966.11 | 988.41 | 988.77 |
| Tet*-$(Pro)_6$-Acd | 1063.48 | 1063.48 | 1063.81 | 1063.82 | 1085.46 | 1085.81 |
| Tet*-$(Pro)_7$-Acd | 1160.53 | 1160.53 | 1160.93 | 1160.63 | 1182.51 | 1182.92 |
| Tet*-$(Pro)_8$-Acd | 1257.58 | 1257.58 | 1257.82 | 1257.70 | 1279.57 | 1279.86 |
| Tet*-$(Pro)_9$-Acd | 1354.64 | 1354.64 | 1354.76 | 1354.89 | 1376.62 | 1377.93 |
| Tet*-$(Pro)_{10}$-Acd | 1451.70 | 1451.70 | 1452.54 | 1452.01 | 1473.67 | 1475.27 |
| Tet*-$(Pro)_{11}$-Acd | 1548.76 | 1548.76 | 1549.44 | 1549.01 | 1570.73 | 1572.50 |
| Tet*-$(Pro)_{12}$-Acd | 1645.80 | 1645.80 | 1647.12 | 1646.04 | 1667.77 | 1668.84 |
| Tet*-$(Pro)_{13}$-Acd | 1742.85 | 1742.85 | 1744.30 | 1743.05 | 1764.82 | 1765.91 |
| Tet*-$(Pro)_{14}$-Acd | 1839.90 | 1839.90 | 1841.27 | 1840.25 | 1861.87 | 1862.39 |
| Leu-$(Pro)_4$-Acd | 826.41 | 826.41 | 826.86 | 826.64 | 848.40 | 848.92 |

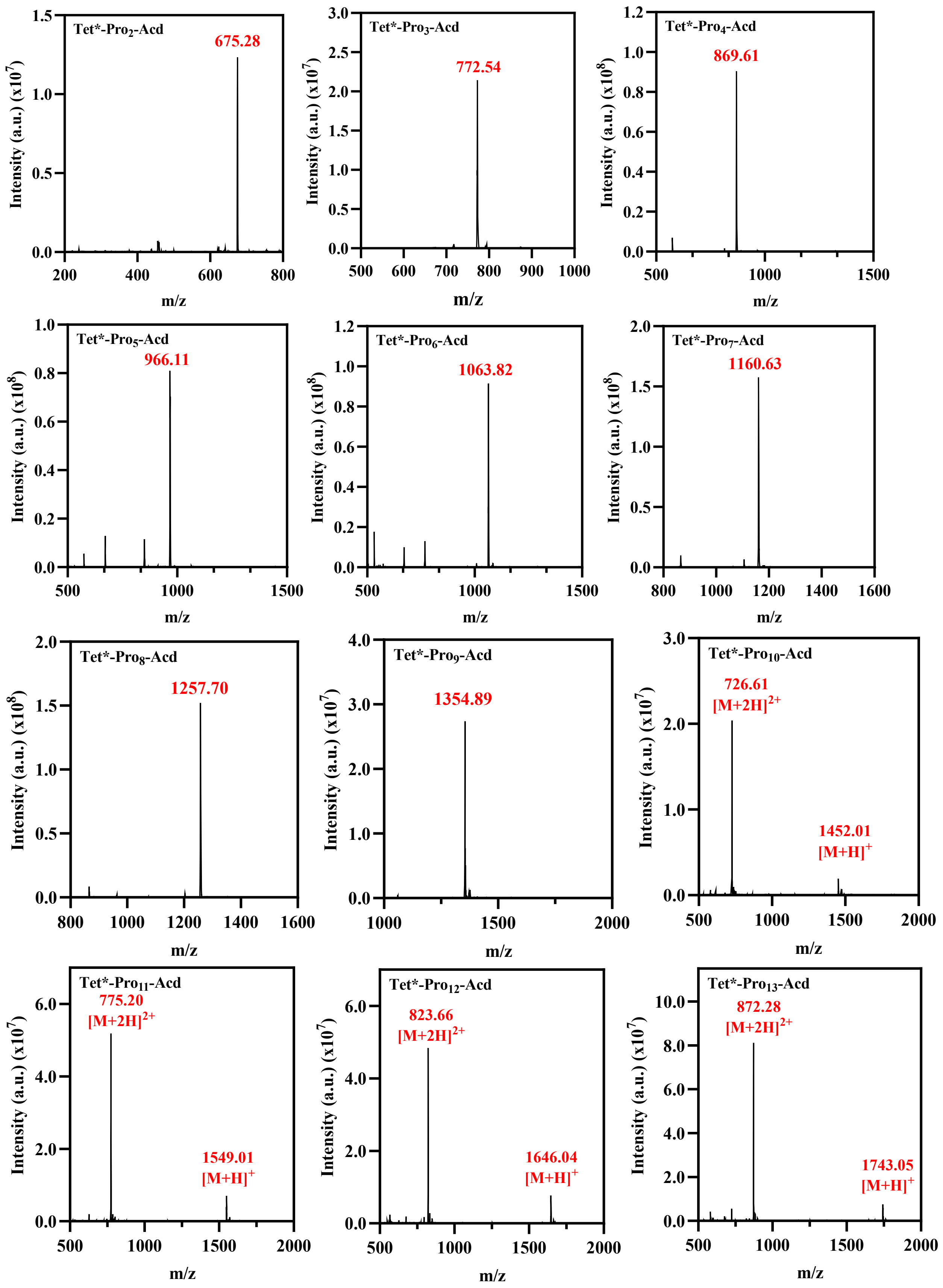
Tet*-Pro2-Acd
675.28
Tet*-Pro3-Acd
772.54
Tet*-Pro4-Acd
869.61
Tet*-Pro5-Acd
966.11
Tet*-Pro6-Acd
1063.82
Tet*-Pro7-Acd
1160.63
Tet*-Pro8-Acd
1257.70
Tet*-Pro9-Acd
1354.89
Tet*-Pro10-Acd
726.61
[M+2H]2+
1452.01
[M+H]+
Tet*-Pro11-Acd
775.20
[M+2H]2+
1549.01
[M+H]+
Tet*-Pro12-Acd
823.66
[M+2H]2+
1646.04
[M+H]+
Tet*-Pro13-Acd
872.28
[M+2H]2+
1743.05
[M+H]+
Intensity (a.u.) (x10^7)
Intensity (a.u.) (x10^8)
m/z

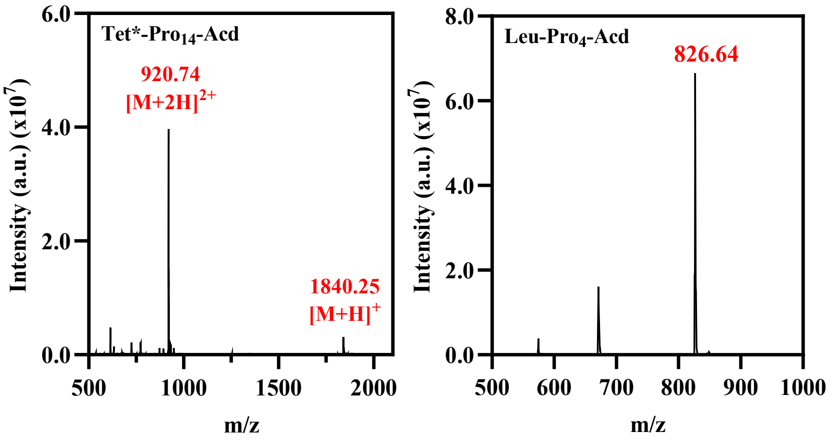


**Figure S3.** Mass spectrum of purified peptides Tet*-Pro$_n$-Acd (n = 2-14) and Leu-Pro$_4$-Acd.

### Proline Ruler FRET Measurements

**Fluorescence Spectroscopy.** Dry proline ruler peptides were diluted in $CH_3CN/H_2O$ (10:90 v/v) and absorbance spectra from 300 to 600 nm were scanned using quartz fluorometer cells with path lengths of 1.00 cm. The peptides were diluted to concentrations of approximately 10 μM in 0.1 M phosphate buffer (57 mM $Na_2HPO_4•7H_2O$, 42 mM $NaH_2PO_4.H_2O$, pH 7.0 adjusted with HCl), as determined by absorbance at 386 nm ($\varepsilon_{386}$ = 5700 $M^{-1}cm^{-1}$).[3] Fluorescence spectra for each peptide were collected at 450 nm with an excitation wavelength of 385 nm in triplicate using Tecan Spark plate reader (5 nm bandwidth, 40 μs integration time, 31000 μm Z-position, 4 sec linear shaking) with a nonsterile Greiner black, flat μClear, 384 well low volume microplate. Also, all Acd containing peptides were excited at 385 nm, with emission measured from 400 nm to 600 nm (30 flashes, 5 nm bandwidth). Examples of fluorescence spectra and associated UV spectra are shown in Figure S4.

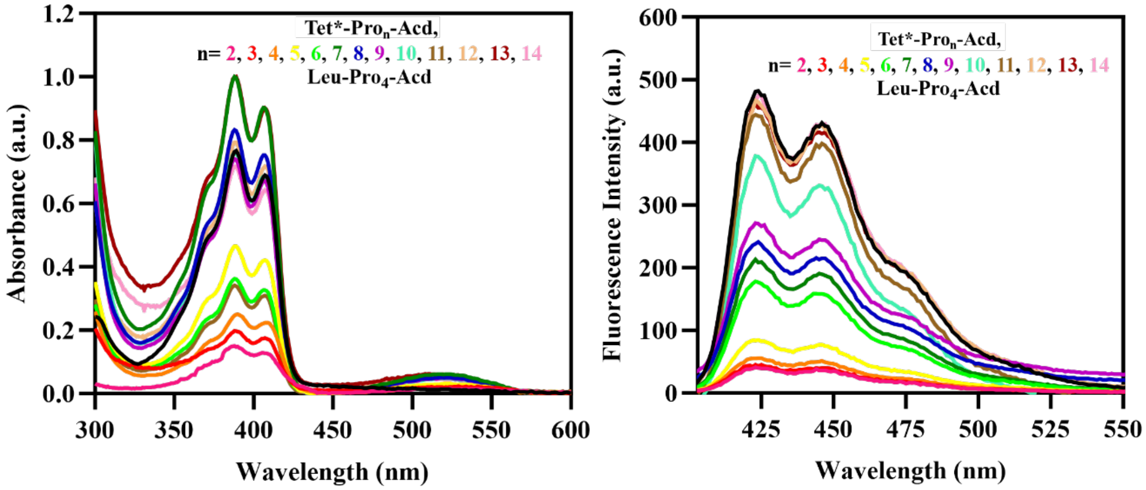


**Figure S4**. Left: UV absorption spectra collected between 300 and 600 nm. Absorbance is normalized. Right: Fluorescence emission spectra collected between 400 and 600 nm with excitation at 385 nm for Pro Series peptides. Emission spectra are colored according to the corresponding absorption spectrum. The acetonitrile concentration after dilution in 0.1 M phosphate buffer is <1% for all peptides.

**Förster Distance Calculation.** The Förster distance, $R_0$, is given in Å by Equation S2[4-5]

$$R_0^6 = \frac{9000(ln10)\kappa^2 Q_D J}{128\pi^5 n^4 N_A} \tag{S2}$$

Where $\kappa^2$ is a geometrical factor that relates the orientation of the donor and acceptor transition moments, $Q_D$ is the quantum yield of the donor, $n$ is the index of refraction of the solvent, $N_A$ A is Avogadro's number, and $J$ is the spectral overlap integral defined in units of $M^{-1}$ •$cm^{-1}$ •$nm^4$ . Combining constants and rearranging gives $R_0$ as

$$R_0 = 0.211(Q_D \kappa^2 n^{-4} J)^{1/6} \tag{S3}$$

*J* is formally defined as

$$J = \int_0^\infty f_D(\lambda)\varepsilon_A(\lambda)\lambda^4 \, d\lambda \tag{S4}$$

Where $\varepsilon_A(\lambda)$ is the molar extinction coefficient of the acceptor at each wavelength λ and $f_D(\lambda)$ is the normalized donor emission spectrum given by

$$f_D(\lambda) = \frac{F_{D\lambda}(\lambda)}{\int_0^\infty F_{D\lambda}(\lambda)d(\lambda)} \tag{S5}$$

where $F_{D\lambda}(\lambda)$ is the fluorescence of the donor at each wavelength λ. Fluorescence spectra of Acd in water were integrated with GraphPad Prism (version 10.0.0, GraphPad Software, Boston, Massachusetts, USA) from 400 to 600 nm to calculate $f_D(\lambda)$. UV-Vis spectra of Tet* were used to determine $\varepsilon_A(\lambda)$. The value of $\varepsilon_{540}$ = 300 $M^{-1}$ •$cm^{-1}$ was used to calculate J, which was found to be $9.51 \times 10^{12} M^{-1} \cdot cm^{-1} \cdot nm^4$. Substituting this result into Equation S2, as well as 0.95 for the quantum yield of Acd in water,[7] 1.33 for the index of refraction of water, and 2/3 for $\kappa^2$ gives the Förster distance of 24 Å for the Tet*-Acd FRET pair.

For the theoretical FRET distance dependence calculation, Equation S6 was used, where $R_0$ the distance was fixed at 24 Å (~ 8 prolines).

$$\frac{F}{F_0} = \left\{1 - \frac{1}{1+\left(\frac{R}{R_0}\right)^6}\right\} \tag{S6}$$

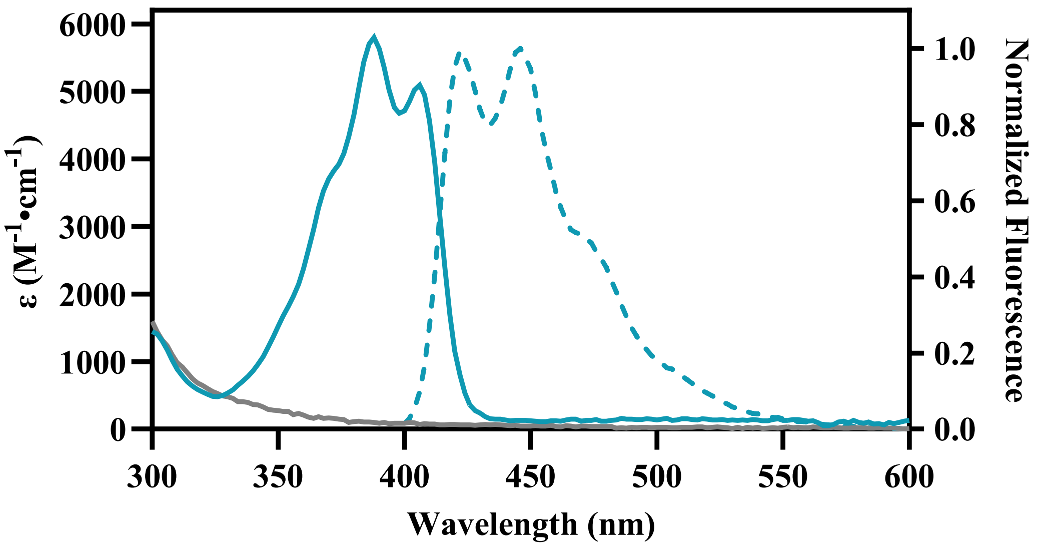


**Figure S5.** Acd/Tet*-sTCO-OH adduct FRET Quenching. Acd/Tet* spectra showing Acd absorbance (solid blue line) and emission from 400 to 600 nm with excitation at 385 nm (dashed blue line) and Tet*-sTCO-OH adduct absorbance (grey solid line).

For Acd and the Tet*-sTCO-OH adduct spectral overlap calculation, UV-Vis spectra of the Tet*-sTCO-OH adduct were used to determine $\varepsilon_A(\lambda)$ which gave J as $1.99 \times 10^{12} M^{-1} \cdot cm^{-1} \cdot nm^4$. Using this J value, an $R_0$ of 18 Å can be calculated for Acd/Tet-sTCO FRET.

**Dequenching of Proline Ruler Peptides**

**sTCO-OH reaction with peptides**. Fresh 15 μM solutions of each peptide were prepared in 0.1 M phosphate buffer. Fluorescence spectra for each peptide were collected at 450 nm with an excitation wavelength of 385 nm in triplicate using Tecan Spark plate reader (5 nm bandwidth, 40 μs integration time, 31000 μm Z-position, 4 sec linear shaking) with a nonsterile Greiner black, flat μClear, 384 well low volume microplate. 150 μM of sTCO-OH was then added to each peptide well. The mixtures were allowed to react for 15 minutes with shaking (500 rpm) at 37 °C. Again, fluorescence spectra for each peptide were collected at 450 nm with an excitation wavelength of 385 nm under the same conditions as unlabeled peptides.

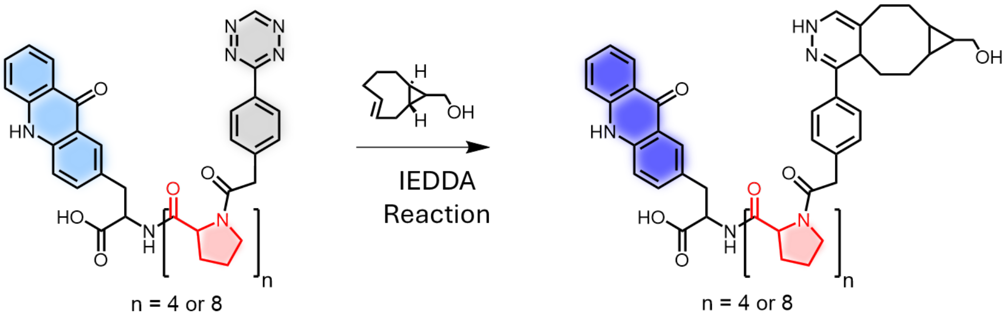


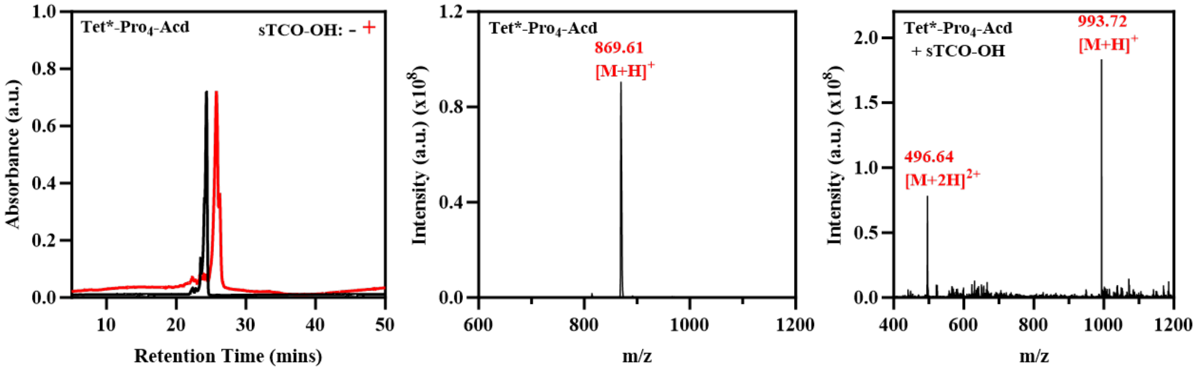


**Figure S6**. Top: Schematic representation of Tet* labeling with sTCO-OH. Bottom Left: Analytical HPLC Chromatograms of Tet*-$Pro_4$-Acd before and after reaction with sTCO-OH. Absorbance is normalized. Peak absorption measured at 215 nm. Bottom Middle: MALDI-MS spectra of Tet*-$Pro_4$-Acd. Bottom Right: MALDI-MS spectra of Tet*-$Pro_4$-Acd after sTCO-OH labeling.

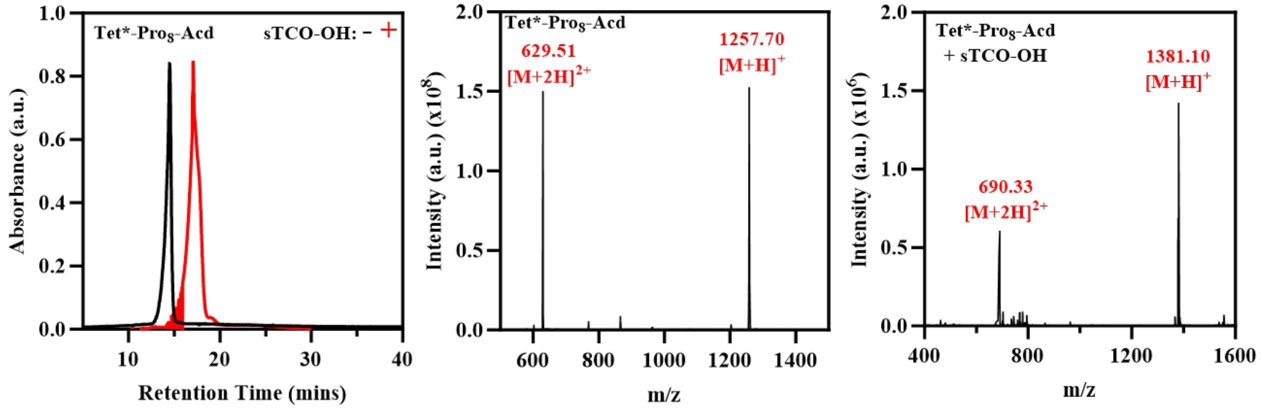


**Figure S7**. Left: Analytical HPLC Chromatograms of Tet*-Pro$_8$-Acd before and after reaction with sTCO-OH. Peak absorbance is normalized that measured at 215 nm. Middle: MALDI-MS spectra of Tet*-Pro$_8$-Acd. Right: MALDI-MS spectra of Tet*-Pro$_8$-Acd after sTCO-OH labeling.

**Photolysis of peptides**. 15 μM of Tet* -Pro$_5$-Acd was prepared in 0.1 M phosphate buffer. These peptides were irradiated with UV light of 254 nm under different time intervals (0, 5, 10, 25, 35, 50, 60) and fluorescence spectra were collected at 450 nm with an excitation wavelength of 385 nm in triplicate using Tecan Spark plate reader under the same conditions as described before.

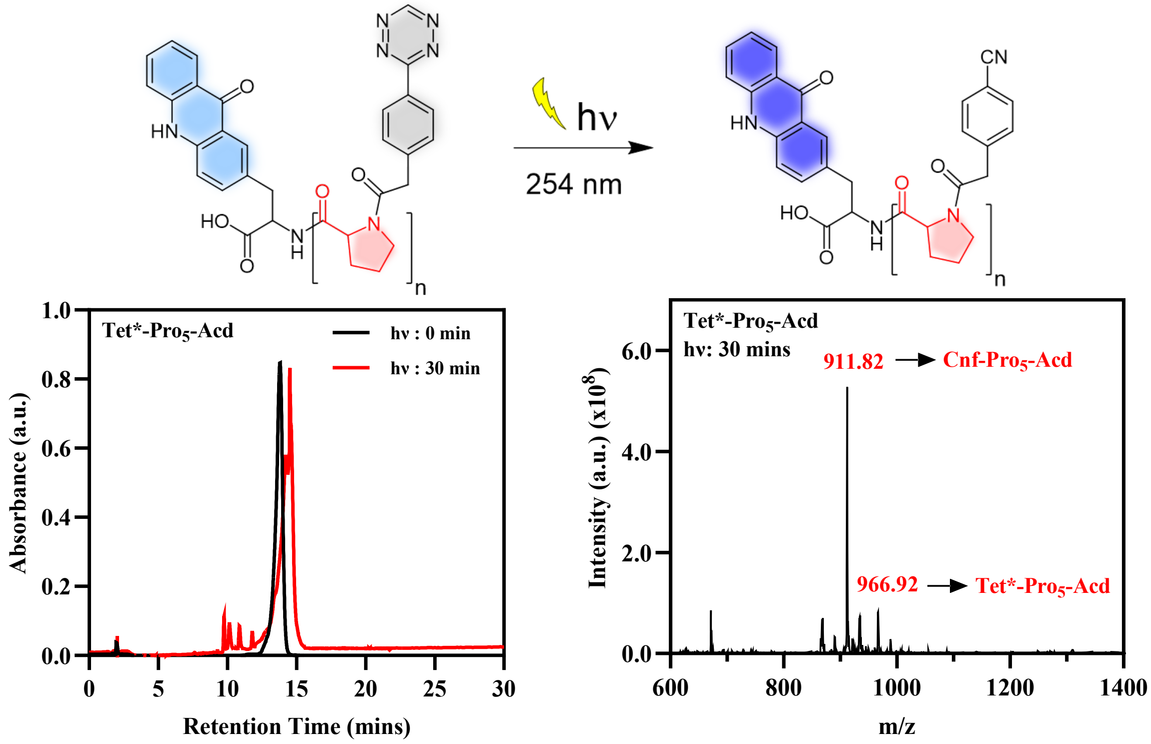


**Figure S8**. Top: Schematic representation of Tet* photolysis at 254 nm leading to dequenching of Acd fluorescence *via* formation of Cnf. Bottom Left: Analytical HPLC Chromatograms of Tet*-Pro$_5$-Acd before and after irradiation. Absorbance is normalized. Peak absorbance measured at 215 nm. Bottom Right: MALDI-MS spectra of Tet*-Pro$_5$-Acd after 30 minutes of irradiation.

**Cloning of Dual Label Plasmids: CaM, LexA, and RecA**

**Previously Cloned constructs**. The gene containing full-length calmodulin (CaM) was previously cloned into the pTXB1 vector containing a C-terminal MxeGyrA intein, followed by a $His_6$ purification tag.[8] The TAG codon for Acd incorporation at position 113 via amber stop codon suppression was previously inserted using QuikChange® PCR, yielding the pTXB1-CaM-$TAG_{113}$-GyrA-$His_6$ plasmid.[9]

**Cloning of CaM plasmid**. All PCR reactions were carried out on a T100 thermocycler from Bio-Rad (Hercules, CA, USA). DNA concentrations were determined with a Tecan Spark plate reader. Agarose gels were visualized on G:Box mini (Syngene).
To incorporate Tet at position 93 of CaM, PCR was performed on the pTXB1-CaM-$TAG_{113}$-GyrA-$His_6$ plasmid using the NEB Q5 Hot Start HiFi Master Mix to mutate the Phe codon to TAA (primers for pTXB1CaM-$TAA_{93}$-$TAG_{113}$-GyrA-$His_6$, Table S6). The resulting DNA was transformed into DH5α cells (NEB) and purified using the Miniprep kit (Qiagen).

**Cloning of LexA dual-label plasmids.** To make pET41-LexA-$TAA_{81}$-$TAG_{86}$-His, the original TAA stop codon from pET41-LexA-$TAG_{86}$-His (reported by Hostetler et al.[12]) was first mutated to TGA using cassette mutagenesis. Briefly, the parent plasmid (pET41-LexA-$TAG_{86}$-His) was digested using AvrII and XhoI (NEB). Digest was assessed by agarose gel, and the desired fragment was obtained by gel purification (Monarch DNA Gel Extraction Kit, NEB). An oligo pair (Table S6) containing the mutation was annealed, phosphorylated using T4 PNK (NEB) in a T100 Thermal Cycler (BioRad), and ligation was completed using T4 DNA ligase (NEB) with overnight cycling ligation. The resulting ligation mixture and negative control were transformed into NEB Turbo competent cells. Plasmid was obtained by miniprep (Qiagen). The mutation was confirmed by Sanger sequencing (Genewiz by Azenta). A second round of cassette mutagenesis was used to mutate the R81 codon to TAA. This cloning was accomplished as described above with the following alterations: a different oligo pair (see Table S6 below) was used to install the TAA mutation, and the pET41-LexA-$TAG_{86}$-His (stop TGA) parent plasmid was digested using SpeI-HF and BmtI-HF (NEB).
For pET41-LexA-$TAG_{81}$-$TAA_{86}$-His, pET41-LexA-$TAA_{86}$-$TAG_{175}$-QM-His, and pET41-LexA-$TAA_{86}$-$TAG_{175}$-K156A-His, the pET41-LexA-$TAG_{86}$-His (stop TGA) parent plasmid was digested using SpeI-HF and HindIII (NEB). Digestion was assessed by agarose gel, and the desired fragment was obtained by gel purification (Monarch DNA Gel Extraction Kit, NEB). Gene fragments containing the desired mutations with flanking SpeI and HindIII restriction enzyme sites were synthesized (Twist Bioscience) (see Table S5 below). The gene fragments were digested with SpeI-HF and HindIII (NEB) and purified by NucleoSpin Gel & PCR Clean-Up Kit (Macherey-Nagel). Ligation was completed using T4 DNA ligase (NEB) with overnight cycling ligation. The resulting ligation mixture and negative control were transformed into NEB Turbo competent cells. Plasmid was obtained by miniprep (Qiagen). The constructs were confirmed by Sanger sequencing (Genewiz by Azenta).
For pET41-LexA-$TAG_{175}$-QM-His, the parent plasmid was pET41-LexA-$TAA_{86}$-$TAG_{175}$_QM-His. Cassette mutagenesis was used as described above with the following alterations: a different oligo pair (see Table S6 below) was used to revert the TAA mutation to the E86 codon, the oligo pair was annealed, but not phosphorylated, and the parent plasmid was digested using BamHI-

HF and EagI-HF (NEB). Additionally, the vector was purified using the NucleoSpin Gel & PCR Clean-Up Kit (Macherey-Nagel). The construct was confirmed by whole-plasmid sequencing (Plasmidsaurus).

For pET41-LexA-$TAG_{175}$-K156A-His, the parent plasmid was pET41- LexA-$TAA_{86}$-$TAG_{175}$-K156A-His. Cassette mutagenesis was used as described above with the following alterations: a different oligo pair (see Table S6 below) was used to revert the TAA mutation to the E86 codon, and the parent plasmid was digested using BamHI-HF and EagI-HF (NEB). Additionally, the vector was purified using the NucleoSpin Gel & PCR Clean-Up Kit (Macherey-Nagel). The construct was confirmed by whole-plasmid sequencing (Plasmidsaurus).

**Cloning of RecA dual-label plasmid.** To make the RecA dual label construct plasmid, a gene fragment containing the sequence of the RecA gene with the desired mutations (codon for R33 changed to TAA and codon for I102 changed to TAG) and flanking SphI and AvrII restriction enzyme sites was synthesized (Twist Bioscience) (see Table S5 below). Both the parent plasmid (pET41-RecA-WT-His)[17] and gene fragment were digested with SphI-HF and AvrII (NEB). For the vector, digestion was assessed by agarose gel, and the desired fragment was obtained by gel purification (Monarch DNA Gel Extraction Kit, NEB). For the insert, the digested gene fragment was purified by NucleoSpin Gel & PCR Clean-Up Kit (Macherey-Nagel). Ligation was completed using T4 DNA ligase (NEB) with overnight cycling ligation. The resulting ligation mixture and negative control were transformed into NEB Turbo competent cells. Plasmid was obtained by miniprep (Qiagen). The construct was confirmed by whole-plasmid sequencing (Plasmidsaurus).

**Table S5: Gene fragments used for plasmid cloning**

| **Construct** | **Gene Fragment Sequence (5'-3')** |
|---|---|
| pET41-LexA-$TAG_{81}$-$TAA_{86}$-His | AGAAGGGTTGCCACTAGTAGGTtagGTGGCGGCCGGTtaaCCACTGCTAGC GCAACAGCATATTGAAGGTCATTACCAGGTGGATCCTTCCTTATTCAAGC CGAATGCTGATTTCCTGCTGCGCGTCAGCGGGATGTCGATGAAAGATATT GGCATTATGGATGGAGATCTGCTGGCAGTGCATAAAACTCAGGACGTCC GTAACGGTCAGGTCGTTGTCGCACGTATTGATGACGAAGTTACCGTTAAA CGTTTGAAAAAACAGGGCAATAAAGTCGAGCTCTTGCCAGAAAATTCCGA GTTTAAACCAATTGTCGTTGACCTTCGTCAGCAAAGCTTCACCATTGAAG G |
| pET41-LexA-$TAA_{86}$-$TAG_{175}$-QM-His | AGAAGGGTTGCCACTAGTAGGTCGTGTGGCGGCCGGTtaaCCACTGccaG CGCAAtggCATATTGAAGGTCATTACCAGGTGGATCCTTCCTTATTCAAGC CGAATGCTGATTTCCTGCTGCGCGTCAGCGGGATGTCGATGAAAGATATT GGCATTATGGATGGAGATCTGCTGGCAGTGCATAAAACTCAGGACGTCC GTAACGGTCAGGTCGTTGTCGCACGTATTGATGACgcaGTTACCGTTgcaC GTTTGAAAAAACAGGGCAATAAAGTCGAGCTCTTGCCAGAAAATTCCGAG TTTtagCCAATTGTCGTTGACCTTCGTCAGCAAAGCTTCACCATTGAAGG |
| pET41-LexA-$TAA_{86}$-$TAG_{175}$-K156A-His | AGAAGGGTTGCCACTAGTAGGTCGTGTGGCGGCCGGTtaaCCACTGCTAG CGCAACAGCATATTGAAGGTCATTACCAGGTGGATCCTTCCTTATTCAAG CCGAATGCTGATTTCCTGCTGCGCGTCAGCGGGATGTCGATGAAAGATAT TGGCATTATGGATGGAGATCTGCTGGCAGTGCATAAAACTCAGGACGTCC GTAACGGTCAGGTCGTTGTCGCACGTATTGATGACGAAGTTACCGTTgca CGTTTGAAAAAACAGGGCAATAAAGTCGAGCTCTTGCCAGAAAATTCCGA GTTTtagCCAATTGTCGTTGACCTTCGTCAGCAAAGCTTCACCATTGAAGG |
| pET41-RecA-$TAA_{33}$-$TAG_{102}$-His | ccgccgccgcaaggaatggtgcatgcaaggagatggcgcccaacagtcccccggccacggggcctgcc accatacccacgccgaaacaagcgctcatgagcccgaagtggcgagcccgatcttccccatcggtgatgt cggcgatataggcgccagcaaccgcacctgtggcgccggtgatgccggccacgatgcgtccggcgtaga |

| | |
|---|---|
| | ggatcgagatcgatctcgatcccgcgaaattaatacgactcactataggggaattgtgagcggataacaatt<br>cccctctagaaataattttgtttaactttaagaaggagatatacgatgGCTATCGACGAAAACAAA<br>CAGAAAGCGTTGGCGGCAGCACTGGGCCAGATTGAGAAACAATTTGGTA<br>AAGGCTCCATCATGCGCCTGGGTGAAGACtaaTCCATGGATGTGGAAACC<br>ATCTCTACCGGTTCGCTTTCACTGGATATCGCGCTTGGGGCAGGTGGTCT<br>GCCGATGGGCCGTATCGTCGAAATCTACGGACCGGAATCTTCCGGTAAA<br>ACCACGCTGACGCTGCAGGTGATCGCCGCAGCGCAGCGTGAAGGTAAAA<br>CCTGTGCGTTTATCGATGCTGAACACGCGCTGGACCCAtagTACGCACGT<br>AAACTGGGCGTCGATATCGACAACCTGCTGTGCTCCCAGCCGGACACCG<br>GCGAGCAAGCTTTGGAAATCTGTGACGCCCTGGCGCGTTCTGGCGCAGT<br>AGACGTTATCGTCGTTGACTCCGTGGCGGCACTGACGCCGAAAGCGGAA<br>ATCGAAGGCGAAATCGGCGACTCTCAtATGGGCCTTGCGGCACGTATGAT<br>GAGCCAGGCGATGCGTAAGCTGGCGGGTAACCTTAAGCAGTCCAACACG<br>CTGCTGATCTTCATCAACCAGATCCGTATGAAAATTGGTGTGATGTTCGG<br>TAACCCGGAAACCACTACCGGTGGTAACGCGCTGAAATTCTACGCTAGC<br>GTTCGTCTCGACATCCGTCGTATCGGCGCGGTGAAAGAGGGCGAAAACG<br>TGGTGGGTAGCGAAACCCGCGTGAAAGTGGTGAAGAACAAAATCGCTGC<br>GCCGTTTAAACAGGCTGAATTCCAGATCCTCTACGGCGAAGGTATCAACT<br>TCTACGGCGAACTGGTTGACCTGGGCGTAAAAGAGAAGCTGATCGAGAA<br>AGCAGGCGCGTGGTACAGCTACAAAGGTGAGAAGATCGGTCAGGGTAAA<br>GCGAATGCGACTGCCTGGCTGAAAGATAACCCGGAAACCGCGAAAGAGA<br>TCGAGAAGAAAGTACGTGAGCTCCTGCTGAGCAACCCGAACTCAACGCC<br>GGATTTCTCTGTAGATGATAGCGAAGGCGTAGCAGAAACTAACGAAGATT<br>TTCACCACCATCACCATCATCACCACTgATTGATTAATACCTAGGcaaggctg<br>ctaaacaaagcc |

**Cloning of Synthetase plasmids:**

pULTRAI-Tet3.0 RS[TAA] was cloned as previously described (Addgene #164580).[13] To create pEVOL-AcdA9 RS[TAG], pEVOL-pAzF[TAG] (Addgene # #31186) was first digested with BglII and PstI.[14] Three PCR amplifications were performed, two of which used pBK-*Mj*-Acd A9[15] as a template to amplify two copies of the *Mj* Acd A9 RS genes with TGA stop codons and the appropriate homology ends for directional recombination into the two RS open reading frames of pEVOL (primers pEVOL-Mj-aaRS1-fwd and pEVOL-Mj-aaRS1(TAA>TGA)-rev; pEVOL-Mj-aaRS2-fwd and pEVOL-Mj-aaRS2(TAA>TGA)-rev). The third PCR reaction amplified the fragment connecting the two RS open reading frames from pEVOL-pAzF[TAG], using primers pEVOL-inter-aaRS-fwd and pEVOL-inter-aaRS-rev. These three PCR fragments and the BglII/PstI digested pEVOL-pAzF[TAG] backbone were gel purified and extracted using a Gel Extraction Kit (Macherey-Nagel). The Seamless Ligation Cloning Extract (SLiCE) cloning method[16] was used to recombine these three PCR fragments with the BglII/PstI digested pEVOL-pAzF[TAG] vector simultaneously. The resulting DNA was transformed into chemically competent DH10b cells and purified using a NucleoSpin Plasmid kit (Macherey-Nagel), and the plasmid sequence was confirmed using Sanger sequencing (GeneWiz by Azenta).

Primers:

| Primer | Sequence |
|---|---|
| pEVOL-Mj-aaRS1-fwd | 5' TGGGCTAACAGGAGGAATTAGATCTATGGACG AATTTGAAATGATAAAGAGAAACAC 3' |
| pEVOL-Mj-aaRS1(TAA>TGA)-rev | 5' CCAAGCTGGAGACCGTTTAAACTCATCATAATC TCTTTCTAATTGGCTCTAAAATCTTTATAAGTTC 3' |
| pEVOL-Mj-aaRS2-fwd | 5' TTGTTTACGCTTTGAGGAATCCCATATGGACGA ATTTGAAATGATAAAGAGAAACA 3' |
| pEVOL-Mj-aaRS2(TAA>TGA)-rev | 5' GGCAATTTAGCGTTTGAAACTGCAGTCATAATC TCTTTCTAATTGGCTCTAAAATCTTTATAAGTTCTT C 3' |
| pEVOL-inter-aaRS-fwd | 5' TGAGTTTAAACGGTCTCCAGCTTGG 3' |
| pEVOL-inter-aaRS-rev | 5' ATGGGATTCCTCAAAGCGTAAACAA 3' |

**Synthetase plasmids and their maps used in this study:**

| Synthetase plasmid | origin of replication | GCE platform | resistance |
|---|---|---|---|
| pULTRAI-**Tet3.0** RS[TAA] | CDF | *M. barkeri* | Streptomycin |
| pEVOL-**Acd**A9 RS[TAG] | p15A | *M. janaschii* | Chloramphenicol |

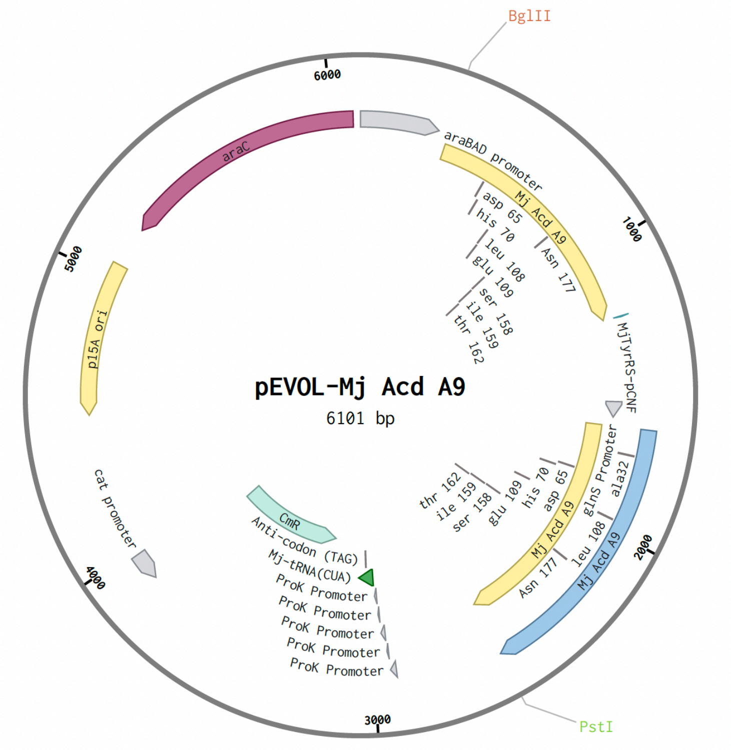


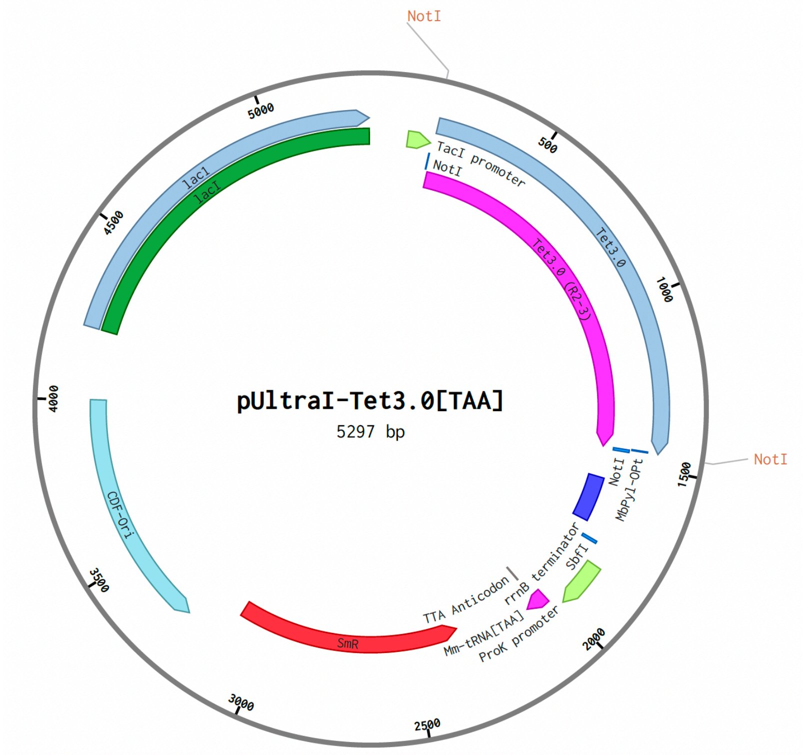


**Plasmids used in this study:**

| Plasmid | Resistance |
|---|---|
| pTXB1-CaM-TAA$_{93}$-TAG$_{113}$-GyrA-His$_{6}$ | Ampicillin |
| pET41-LexA-TAA$_{81}$-TAG$_{86}$-His | Kanamycin |
| pET41-LexA-TAG$_{81}$-TAA$_{86}$-His | Kanamycin |
| pET41-LexA-TAA$_{86}$-TAG$_{175}$-QM-His | Kanamycin |
| pET41-LexA-TAA$_{86}$-TAG$_{175}$-K156A-His | Kanamycin |
| pET41-LexA-TAG$_{175}$-QM-His | Kanamycin |
| pET41-LexA-TAG$_{175}$-K156A-His | Kanamycin |
| pET41-RecA-TAA$_{33}$-TAG$_{102}$-His | Kanamycin |

**Table S6.** Primers used for PCR mutations and cassette mutagenesis

| Primer | Sequence | $T_m$ (˚C) |
|---|---|---|
| CaM(F93TAA) FWD | 5' GTGTAAGACAAGGATGGTAATG 3' | 61 |
| CaM(F93TAA) REV | 5' ACGGAACGCTTCTCTAATTT 3' | |
| LexA(TAA to TGA stop) FWD | 5’TCGAGCACCACCACCACCACCACCACCACTGATTGATTAATAC 3’ | |
| LexA(TAA to TGA stop) REV | 5’CTAGGTATTAATCAATCAGTGGTGGTGGTGGTGGTGGTGGTGC 3’ | |
| LexA(R81 codon to TAA) FWD | 5’CTAGTAGGTTAAGTGGCGGCCGGTTAGCCACTGCTAG 3’ | |
| LexA(R81 codon to TAA) REV | 5’ CAGTGGCTAACCGGCCGCCACTTAACCTA 3’ | |
| LexA(TAA to E86 codon) FWD (to make insert for pET41-LexA-TAG$_{175}$-QM-His) | 5’ GGC CGG TGA ACC ACT GCC TGC GCA ATG GCA TAT TGA AGG TCA TTA CCA GGT G 3’ | |
| LexA(TAA to E86 codon) REV (to make insert for pET41-LexA-TAG$_{175}$-QM-His) | 5’ GAT CCA CCT GGT AAT GAC CTT CAA TAT GCC ATT GCG CAG GCA GTG GTT CAC C 3’ | |
| LexA(TAA to E86 codon) FWD (to make insert for pET41-LexA-TAG$_{175}$-K156A-His) | 5’ GGC CGG TGA ACC ACT GCT TGC GCA ACA GCA TAT TGA AGG TCA TTA CCA GGT G 3’ | |
| LexA(TAA to E86 codon) REV (to make insert for pET41-LexA-TAG$_{175}$-K156A-His) | 5’ GAT CCA CCT GGT AAT GAC CTT CAA TAT GCT GTT GCG CAA GCA GTG GTT CAC C 3’ | |

Tm: Calculated annealing temperature

**Reagents and Buffers for Bacterial Culture**

The following is a general protocol for preparing media for dual ncAA incorporation during protein expression.

**Preparation of starter culture media: non-inducing media (NIM).** Each component in the table below should be prepared separately and sterilized separately. Once sterilized, each component can be stored separately at room temperature indefinitely, provided sterility is maintained. Components should only be mixed when at room temperature and immediately before use.

| | For 500 mL |
|---|---|
| Aspartate (5%, pH 7.5) | 25 mL |
| 18 amino acid mix (25x) | 20 mL |
| 1 M $MgSO_4$ | 1 mL |
| 25x M-Salts | 20 mL |
| 40% (w/v) $\alpha$-D-glucose | 6.25 mL |
| Trace Metals solution (5,000x)* | 100 µl |
| Sterile water (to 500 mL) | 428 mL |

**Expression media: auto-inducing media (AIM).** Each component in the table below should be prepared separately and sterilized separately. Once sterilized, each component can be stored separately at room temperature indefinitely, provided sterility is maintained. *Components should only be mixed when at room temperature and immediately before use*. Note that recipes for all components except the 50x 5052 solution and 20% (w/v) arabinose are in the above section above describing “starter culture media”.

| | For 500 mL |
|---|---|
| Aspartate (5%, pH 7.5) | 25 mL |
| 18 amino acid mix (25x) | 20 mL |
| 1 M $MgSO_4$ | 1 mL |
| 25x M-Salts | 20 mL |
| 50x 5052 solution | 10 mL |
| 20% (w/v) arabinose | 1.25 mL |
| Trace Metals solution (5,000x)* | 100 µL |
| Sterile water (to 500 mL) | 423 mL |

**Preparation of 18 amino acid mix (25x).** Combine the following a total volume of 1 liter. Sterile filter after all amino acids is dissolved. Aliquot into sterile 100-mL bottles or 50 mL conical vials. Stable at 4°C for years w/o contamination, or at -20°C. Final concentration in media for each amino acid is 200 µg/mL.

| | MW | mass (g) |
|---|---|---|
| Glutamic acid, Na salt | 169.1 | 5 |
| Aspartic acid | 133.1 | 5 |

| | | |
|---|---|---|
| Lysine-HCl | 182.6 | 5 |
| Arginine-HCl | 210.7 | 5 |
| Histidine-HCl-$H_2O$ | 209.6 | 5 |
| Alanine | 89.1 | 5 |
| Proline | 115.1 | 5 |
| Glycine | 75.1 | 5 |
| Threonine | 119.1 | 5 |
| Serine | 105.1 | 5 |
| Glutamine | 146.1 | 5 |
| Asparagine-$H_2O$ | 150.1 | 5 |
| Valine | 117.1 | 5 |
| Leucine | 131.2 | 5 |
| Isoleucine | 131.2 | 5 |
| Phenylalanine | 165.2 | 5 |
| Tryptophan | 204.2 | 5 |
| Methionine | 149.2 | 5 |

**Preparation of 25x M-salts.** To prepare the solution, dissolve 88.73 g of anhydrous sodium dibasic ($Na_2HPO_4$) and 85.05 g of anhydrous potassium monobasic ($KH_2PO_4$) in 1 liter of water to achieve 0.625 M concentrations for each phosphate salt. Next, add 66.86 g of $NH_4Cl$ to obtain a 1.25 M solution, followed by 17.75 g of anhydrous sodium sulfate ($Na_2SO_4$) for a 0.125 M solution. Do not adjust the pH of the solution. Sterilize the final solution by autoclaving.

**Preparation of 5000x trace metals.** The following metals are mixed in the ratios mentioned below to prepare the desired 5000x trace metal solution, which is then sterile filtered.

| Amount for 30 ml stock solution | Amount of 30 ml stock solution for 50 ml (5,000x) solution | 1x media concentration |
|---|---|---|
| $CaCl_2 \cdot 2H_2O$ (8.82 g) | 500 µl | 4 µM |
| $MnCl_2 \cdot 4H_2O$ (5.93 g) | 500 µl | 2 µM |
| $ZnSO_4 \cdot 7H_2O$ (8.62 g) | 500 µl | 2 µM |
| $CoCl_2 \cdot 6H_2O$ (1.32 g) | 500 µl | 0.4 µM |
| $CuCl_2$ (807 mg) | 500 µl | 0.4 µM |
| $NiCl_2$ (777 mg) | 500 µl | 0.4 µM |
| $Na_2MoO_4 \cdot 2H_2O$ (1.45 g) | 500 µl | 0.4 µM |
| $Na_2SeO_3$ (1.03 g) | 500 µl | 0.4 µM |
| $H_3BO_3$ (371 mg) | 500 µl | 0.4 µM |
| $FeCl_3$ (486 mg) | 25 ml | 10 µM |
| | Dilute to 50 ml with sterile water | |

**Preparation of 50 x 5052 solution for 500 mL.** To prepare the solution, dissolve 12.5 g of α-D-glucose (2.5% w/v) and 50 g of lactose (10% w/v) in an appropriate volume of Milli-Q (MQ) water. For the glycerol component (25% v/v), either directly add 125 mL of 100% glycerol or prepare a 50% glycerol solution by adding 250 mL of MQ water to a 500 mL graduated cylinder, placing a clean stir bar in the cylinder, and stirring the water while slowly adding 250 mL of 100% glycerol until the total volume reaches 500 mL. Stir for 5–10 minutes to ensure thorough mixing. Note that the lactose may not dissolve immediately; heat the solution in a microwave for 2–3 minutes to dissolve the lactose fully, ensuring it remains in solution before autoclaving. Once all components are dissolved and mixed, autoclave the solution.

**Overexpression of CaM, RecA, and LexA with Tet and Acd**

**Triple plasmid transformation of CaM.** Prepare 1.7 mL Eppendorf tubes containing 1–1.5 µL (~50–200 ng) each of pEVOL-Acd A9 RS[TAG], pULTRAI-Tet3.0 RS[TAA], and the desired pTXB1CaM-$TAA_{93}$-$TAG_{113}$-GyrA-$His_6$ plasmid with a C-terminal intein-$His_6$ fusion gene bearing the amber stop codon and ochre stop codon at the desired position for Acd and Tet incorporation. Thaw 50 µL aliquots of electrocompetent BL21(DE3) cells, gently mix the cells with the plasmids by pipetting, and transfer the mixture into pre-chilled 1 mM electro-cuvettes. Electroporate according to instrument settings, immediately resuspend the cells in 1 mL SOC media, transfer to Eppendorf tubes, and incubate at 37°C with shaking at 250 rpm for 90 minutes. Cells transformed using electroporation were grown on LB/agar plates containing triple antibiotics: for for CaM, 25 µg/mL Chlor, 100 µg/mL Strep, and 50 µg/mL ampicillin (Amp).

**Overexpression of CaM.** For the starter culture, an individual colony was seeded into 5 mL of non-inducing media (NIM) and grown overnight with shaking at 250 rpm at 37°C. Subsequently, 1% of the NIM culture was inoculated into 100 mL of auto-inducing media (AIM) containing 25 µg/mL chloramphenicol, 100 µg/mL streptomycin, and 50 µg/mL ampicillin, and the culture was grown at 37 °C. After 1 hour and 30 minutes of incubation, non-canonical amino acids such as Tet•HCl (0.5 mM final) dissolved in dry DMF and Acd (0.5 mM final) dissolved in water (solubilized by adding an equimolar amount of NaOH from a 5 M stock) were added to the media. The culture was grown until reaching an optical density (OD600) of 1.5 at 37 °C, followed by overnight incubation with shaking at 250 rpm at varied temperatures of 18 °C, 25 °C, or 30 °C.

**Cell lysis and protein purification.** The following day, cells were harvested by centrifugation at 4 °C for 20 min at 2740 RCF (Sorvall GS3 rotor). Pellets were then re-suspended in 15 mL of 40 mM Tris at pH 8.3 for a 100 mL culture that was supplemented with EDTA-free protease inhibitor tablets (Pierce Biotechnology; Waltham, MA, USA) and transferred to a metal cup for sonication. Cells were lysed by sonication on ice with a Q700 probe sonicator (QSonica LLC; Newtown, CT, USA) with the following settings: Amplitude 50, Process Time 5-6 min, Pulse-ON Time 1 s, Pulse-OFF Time 2 s. Crude lysate was then transferred to 50 mL centrifugation tubes and clarified via centrifugation at 23400 RCF for 30 min (Sorvall SS34 rotor). Following centrifugation, the supernatant was removed and transferred to a 50 mL Falcon tube. Then, 5 mL of nickel agarose resin (GoldBio; St. Louis, MO, USA) was added, and the lysate-nickel mixture was incubated with nutation at 4 °C for 1 h. The lysate-nickel mixture was then poured into a 20 mL fritted column, and the flow-through was saved. The remaining resin was then washed with ~20 mL Wash Buffer 1 (50 mM HEPES buffer, pH 7.5), ~20 mL Wash Buffer 2 (50 mM HEPES, 5 mM imidazole, pH 7.5), and eluted with 12 mL Elution Buffer (50 mM HEPES, 300mM imidazole, pH 7.5). The resulting elution was dialyzed against 20 mM Tris, pH 8.0, for 4 h to get rid of the excess imidazole. Then β-mercaptoethanol (Bio-Rad Laboratories; Hercules, CA, USA) was added to crude lysate (200 mM final concentration), and the mixture was allowed to incubate with nutation at room temperature overnight. The resulting cleaved protein was dialyzed against 20 mM Tris, pH 8.0, for 8-10 h. The resulting dialysate was then treated with 5 mL nickel agarose resin and incubated with nutation at 4 °C for 1 h. The mixture was then applied to a 20 mL fritted column, and flow-through containing CaM was collected in a 15 mL Falcon tube. The resulting enriched protein mixture was then purified via FPLC using a 5 mL HiTrap Q-HP column (Cytiva; Marlborough, MA, USA) using the following method: Buffer A: 20 mM Tris, pH 8.0; Buffer B: 20 mM Tris, 1 M NaCl, pH 8.0; Gradient: 0% Buffer B – 5 column volumes, 0 – 10% Buffer B – 5 column volumes, 20-30% Buffer B – 20 column volumes, 30-100% Buffer B – 10 column volumes; flow rate 3 mL/min. The resulting fractions were then assessed for purity via MALDI MS, and pure fractions were combined. Protein was then concentrated, and buffer exchanged into 20 mM Tris pH 8.0 to a final concentration of 100-200 µM via Amicon 3 kDa MWCO filters (Millipore Sigma; St. Louis, MO, USA). Purified protein was aliquoted into 1.5 mL tubes and stored at -80 °C until further use.

**Overexpression of RecA mutant:** In a 1.7 mL eppendorf tube 50 µL aliquot of electrocompetent BL21(DE3) cells were transformed with three plasmids 1–1.5 µL (~50–200 ng) each, one containing the recombinant RecA gene bearing the amber stop codon and ochre stop codon at the desired position for Acd and Tet incorporation respectively, and the other two encoding the aminoacyl tRNA synthetase with improved specificity for Acd and Tet incorporation. The mixture was then transferred into pre-chilled 1 mM electro-cuvettes. After electroporation according to the instrument settings, the cells were immediately resuspended in 1 mL SOC media, transferred to an Eppendorf tube, and incubated at 37 °C with shaking at 250 rpm for 90 minutes. Cells transformed using electroporation were grown on LB/agar plates at 37 °C overnight with 25 µg/mL chloramphenicol, 100 µg/mL streptomycin, and 50 µg/mL kanamycin as required antibiotics. For starter culture, an individual colony was seeded into 5 mL of non-inducing media (NIM) with selective antibiotics and grown at 37°C overnight with shaking at ~250 rpm. To scale up expressions, ~1% inoculum (2 mL) of NIM culture was added to 200 mL of auto-induction media in 1 L baffled flasks with selective antibiotics and grown at 37 °C. After ~1-2 hours, solubilized Acd and Tet were added to a final concentration of 0.5 mM, and continued growth at 37 °C until $OD_{600}$ was ~1.5, and then the temperature was reduced to 30 °C. After growth for 20 hours, cells were harvested and stored at -20 °C.

**Cell lysis and protein purification:** Cells were resuspended in lysis buffer containing 50 mM sodium phosphate, pH 8.0, 500 mM KCl, 500 mM NaCl, 25 mM imidazole, 250 U/mL benzonase nuclease (Sigma), and 0.25 mg/mL lysozyme (Sigma); cells were then lysed using a sonicator. Lysates were then centrifuged (Sorvall SS34 rotor) for 30 mins at 26900 RCF. Clarified lysates were allowed to flow over HisPur cobalt resin (Thermo Fisher) pre-equilibrated with the respective lysis buffer. The resin was then washed with 20 column volumes of RecA cobalt wash buffer (50 mM sodium phosphate pH 8.0, 500 mM KCl, 500 mM NaCl, 100 mM imidazole), followed by elution in 3 column volumes of RecA cobalt elution buffer (50 mM sodium phosphate, pH 8.0, 500 mM KCl, 500 mM NaCl, 400 mM imidazole). After HisPur cobalt resin, for further purification, RecA samples were precipitated by incubation for 30 min at 4 °C using 48% (final) saturation ammonium sulfate, and the pellet was retained after spinning at 3,000 rcf for 30 min. Pellets were resuspended in RecA heparin load/wash buffer (20 mM Tris–HCl, pH 8.0, 20 mM NaCl, 2 mM tris(2-carboxyethyl)phosphine (TCEP)) and dialyzed overnight at 4 °C against the same buffer to remove residual ammonium sulfate. All samples were loaded onto Heparin HiTrap HP 5 ml columns (Cytiva) using an ÄKTAPurifier system. The column was washed with 0% and 10% of wash buffer and eluted with 30% of elution buffer (RecA: 20 mM Tris–HCl, pH 8.0, 1.5 M NaCl, 2 mM TCEP), and all fractions were collected. Fractions were dialyzed overnight at 4 °C into RecA storage buffer (20 mM Tris–HCl, pH 8.0, 200 mM NaCl, 2 mM TCEP, 10% glycerol) and stored at -80 °C.

**Overexpression of LexA mutant:** In a 1.7 mL eppendorf tube 50 μL aliquot of electrocompetent BL21(DE3) cells were transformed with three plasmids 1–1.5 μL (~50–200 ng) each, one containing the recombinant LexA gene bearing the amber stop codon and ochre stop codon at the desired position for Acd and Tet incorporation respectively, and the other two encoding the aminoacyl tRNA synthetase with improved specificity for Acd and Tet incorporation. The mixture was then transferred into pre-chilled 1 mM electro-cuvettes. After electroporation according to the instrument settings, the cells were immediately resuspended in 1 mL SOC media, transferred to an Eppendorf tube, and incubated at 37 °C with shaking at 250 rpm for 90 minutes. Cells transformed using electroporation were grown on LB/agar plates at 37 °C overnight with 25 μg/mL chloramphenicol, 100 μg/mL streptomycin, and 50 μg/mL kanamycin as required antibiotics. For starter culture, an individual colony was seeded into 5 mL of non-inducing media (NIM) with selective antibiotics and grown at 37°C overnight with shaking at ~250 rpm. To scale up expressions, ~1% inoculum (2 mL) of NIM culture was added to 200 mL of auto-induction media in 1 L baffled flasks with selective antibiotics and grown at 37 °C. After ~1-2 hours, solubilized Acd and Tet were added to a final concentration of 0.5 mM, and continued growth at 37 °C until $OD_{600}$ was ~1.5, and then the temperature was reduced to 30 °C. After growth for 24 hours, cells were harvested and stored at -20 °C.

**Cell lysis and protein purification:** Cells were resuspended in lysis buffer containing 20 mM sodium phosphate pH 6.8, 500 mM NaCl, 25 mM imidazole, 250 U/mL benzonase nuclease (Sigma) and 0.25 mg/mL lysozyme (Sigma); cells were then lysed using a sonicator. Lysates were then centrifuged (Sorvall SS34 rotor) for 15 mins at 26900 RCF. Clarified lysates were affinity purified with HisPur cobalt resin (Thermo Fisher) using a 25 to 400 mM imidazole gradient. To further purify the HisPur eluted protein, size exclusion chromatography was done using Superdex™ 200 pg in 50 mM Tris-HCl pH 7.0, 200 mM NaCl, 0.5 mM EDTA, and 10% glycerol. Fractions containing purified Acd/Tet-labeled LexA were stored at -80 °C.

**Overexpression of WT-RecA:** The RecA construct was overexpressed in BL21(DE3) cells containing the recombinant RecA gene. Overnight cultures were grown in LB media with kanamycin at 37 °C with shaking at ~250 rpm and diluted 1:100 into fresh 1L LB media. Cultures were grown to an optical density at 600 nm of 1.2 before induction with 1 mM IPTG. After 5 hours at 37 °C, harvested cells were resuspended in RecA lysis buffer with cOmplete EDTA-free protease inhibitor cocktail tablet (Sigma). Cell lysis and purification were similar to those of previously described RecA mutants.

**Overexpression of WT-LexA:** The LexA construct was overexpressed in BL21(DE3) cells containing the recombinant LexA gene. Overnight cultures were grown in LB media with kanamycin at 37 °C with shaking at ~250 rpm and diluted 1:100 into fresh 1L LB media. Cultures were grown to an optical density at 600 nm of 0.6-0.8 before induction with 1 mM IPTG. After 20 hours at 37 °C, harvested cells were resuspended in LexA lysis buffer with cOmplete EDTA-free protease inhibitor cocktail tablet (Sigma).

Cell lysis and purification were similar to those of previously described LexA mutants.

**Purification and Characterization of CaM$\tau_{93}\delta_{113}$**

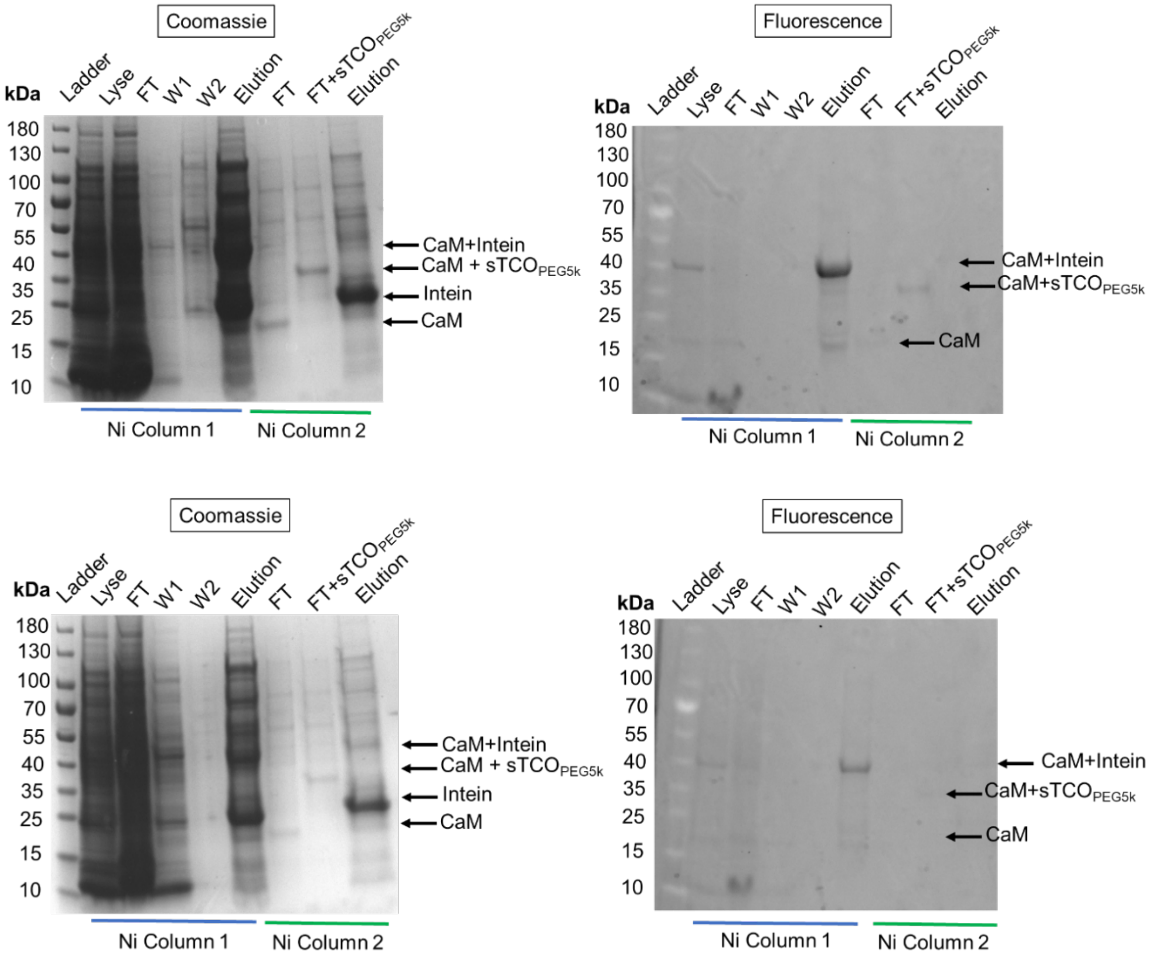


**Figure S9**. Top: SDS-PAGE analysis of expression of CaM$\tau_{93}\delta_{113}$ at 25 ˚C (top) and 30 ˚C (bottom). Left: gel code blue stain and Right: fluorescence. MW = molecular weight in kDa. Lyse = lysate, FT = flow through, W1 = wash 1, and W2 = wash 2. Excitation Wavelength of Acd is 350 nm for fluorescence gel imaging.

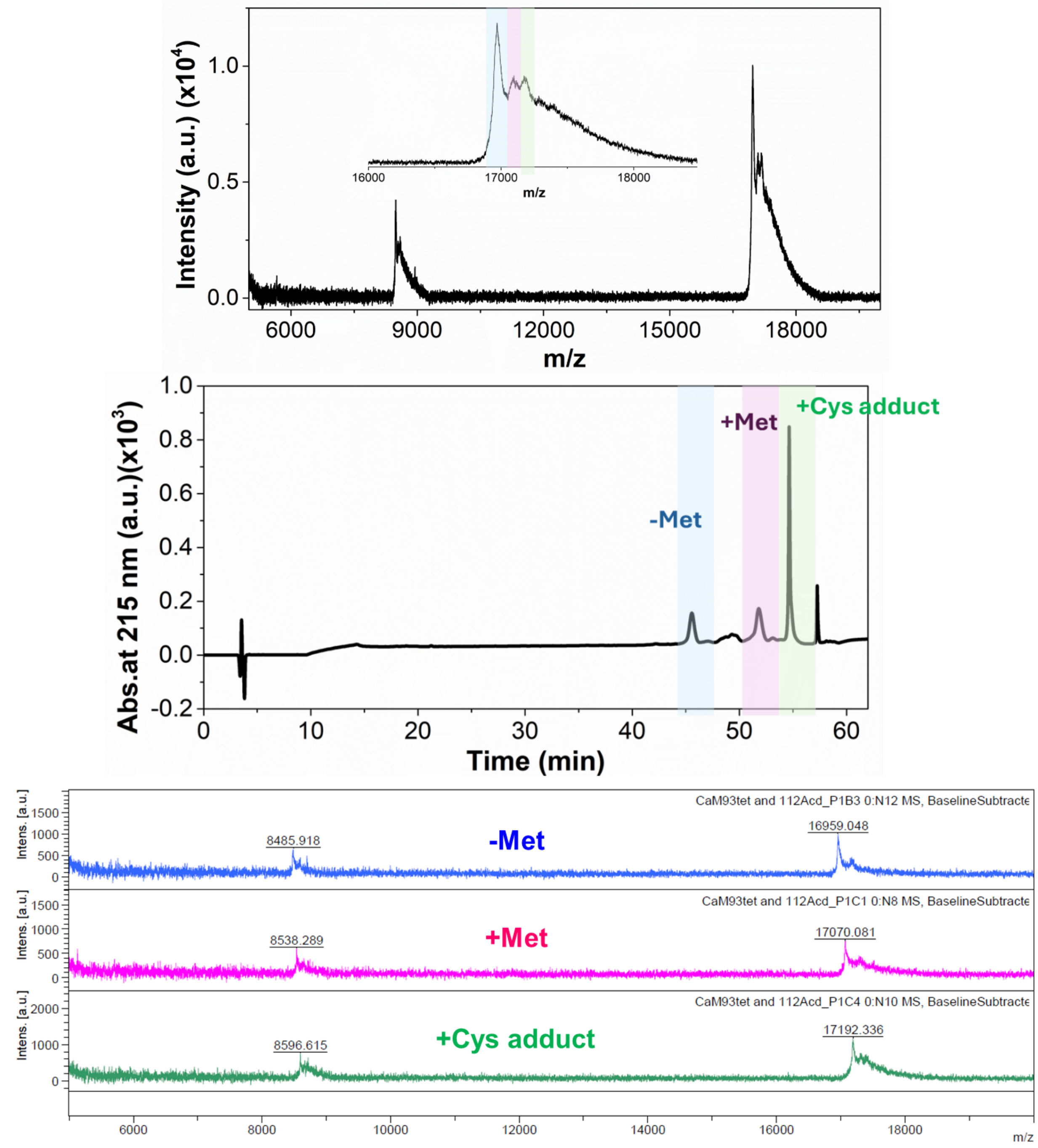


**Figure S10**. Top: MALDI-TOF-MS characterization of FPLC purified $CaM\tau_{93}\delta_{113}$. Middle: Analytical HPLC chromatogram showing the separation profile of $CaM\tau_{93}\delta_{113}$ from Cys adduct of Tet. *Bottom:* MALDI-MS analysis of the individual fractions collected from the analytical HPLC, confirming the purity of full-length $CaM\tau_{93}\delta_{113}$, as well as the presence of its variant, the N-terminal methionine-cleaved form and the cysteine adduct with Tet.

### Purification and Characterization of RecA$\tau_{33}\delta_{102}$

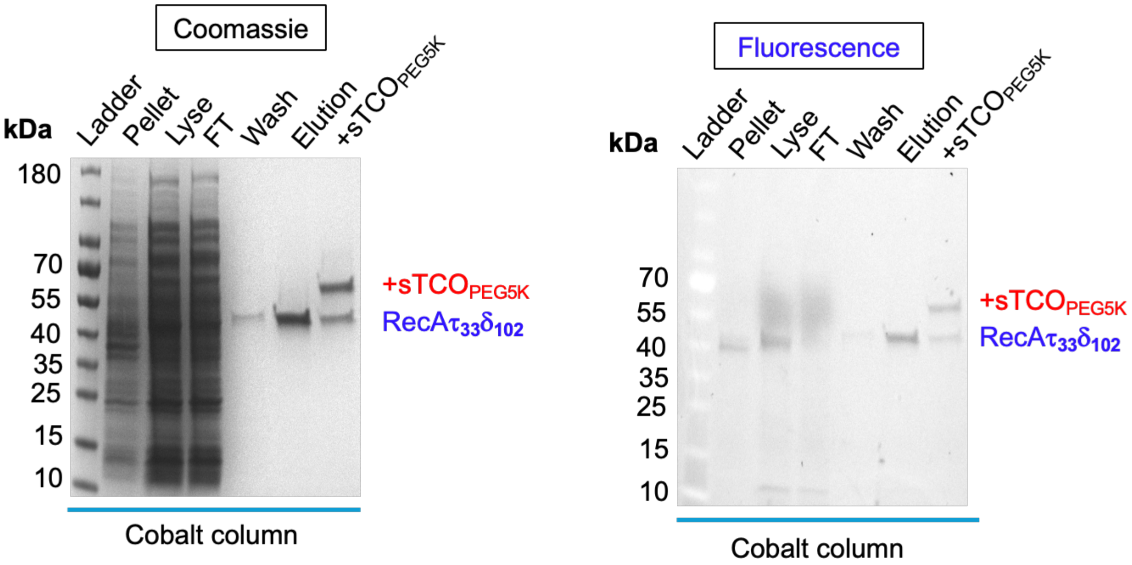


**Figure S11**. SDS-PAGE analysis of expression of RecA$\tau_{33}\delta_{102}$ at 30 ˚C. Left: gel code blue stain, and Right: Fluorescence. MW = molecular weight in kDa. Lyse = lysate, FT = flow through. Excitation wavelength of Acd is 350 nm for fluorescence gel imaging.

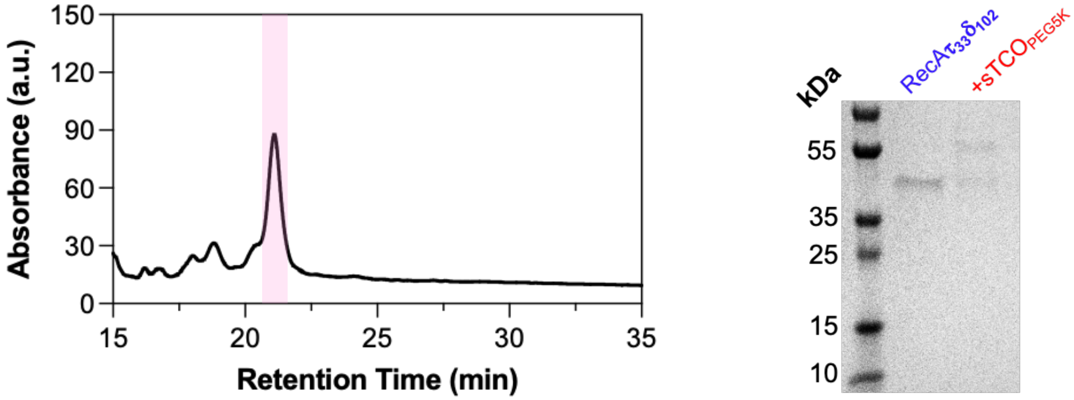


**Figure S12**. Left: Analytical HPLC chromatogram showing the purity of RecA$\tau_{33}\delta_{102}$. Peak absorption measured at 215 nm. Right: Gel analysis of RecA$\tau_{33}\delta_{102}$ after analytical HPLC and confirmation of Tet incorporation by reaction with sTCO$_{PEG5k}$.

### Purification and Characterization of LexA$\tau_{81}\delta_{86}$

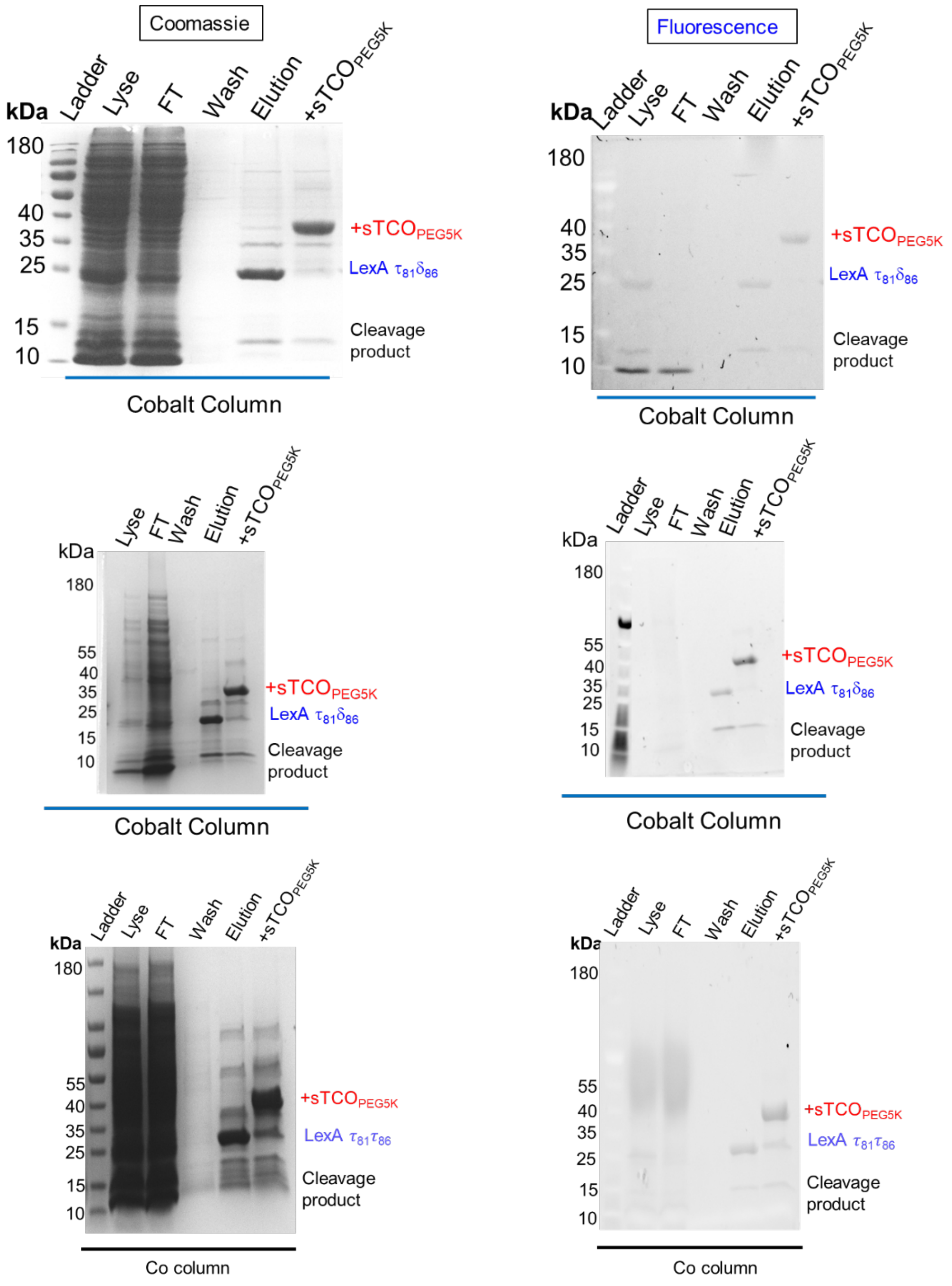


**Figure S13**. SDS-PAGE analysis of expression of LexA$\tau_{81}\delta_{86}$ at 30 ˚C in three independent protein expressions. Left: gel code blue stain, and Right: Fluorescence. MW = molecular weight in kDa. Lyse = lysate, FT = flow through. Excitation wavelength of Acd is 350 nm for fluorescence gel imaging.

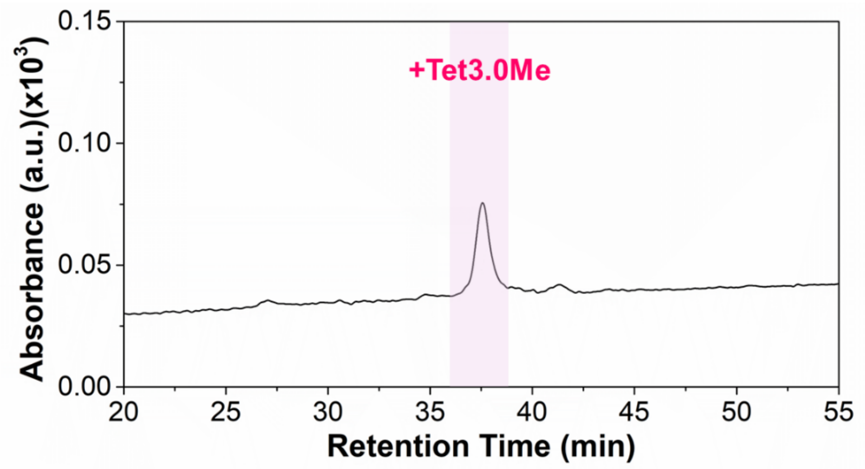


**Figure S14**. Analytical HPLC chromatogram showing the purity of LexA$\tau_{81}\delta_{86}$. Peak absorption measured at 215 nm.

**Purification and Characterization of LexA$\delta_{81}\tau_{86}$**

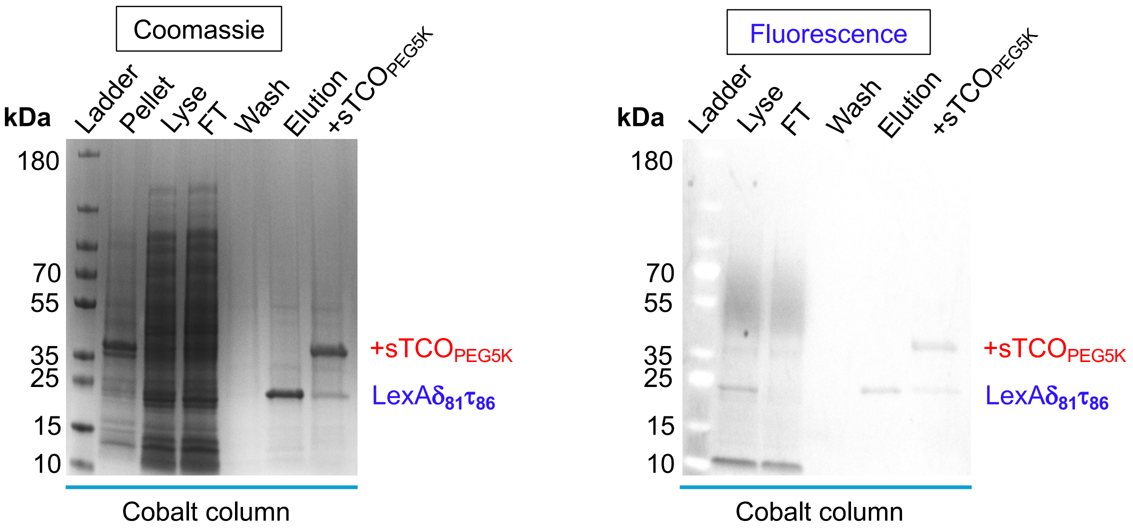


**Figure S15**. SDS-PAGE analysis of expression of LexA$\delta_{81}\tau_{86}$ at 30 ˚C. Left: gel code blue stain, and Right: Fluorescence. MW = molecular weight in kDa. Lyse = lysate, FT = flow through. Excitation wavelength of Acd is 350 nm for fluorescence gel imaging.

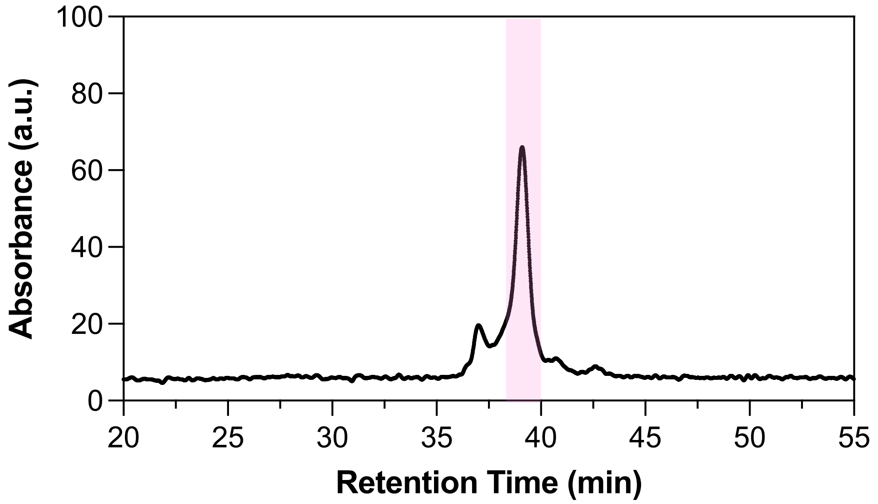


**Figure S16**. Analytical HPLC chromatogram showing the purity of LexA$\delta_{81}\tau_{86}$. Peak absorption measured at 215 nm.

### Purification and Characterization of LexA$\tau_{86}\delta_{175}$ (K156A)

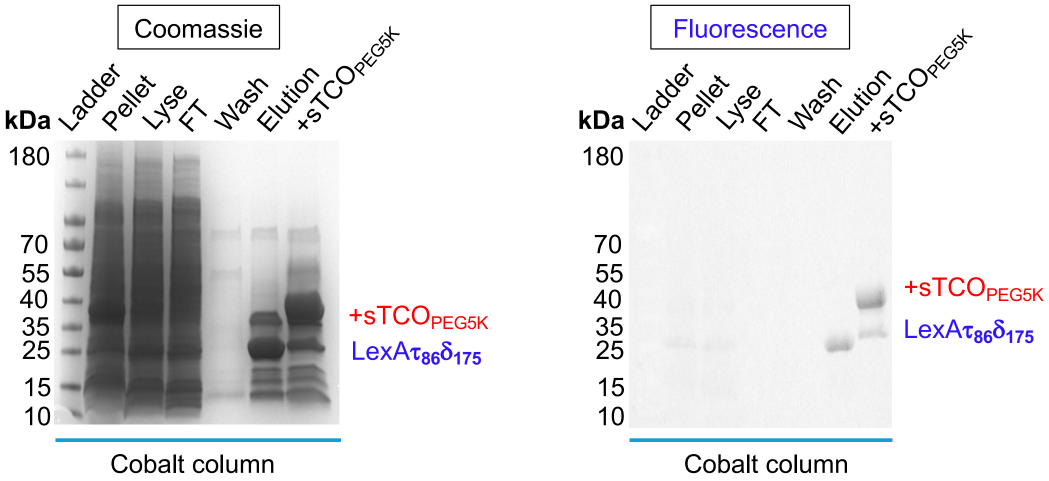


**Figure S17**. SDS-PAGE analysis of expression of LexA$\tau_{86}\delta_{175}$ (K156A) at 30 ˚C. Left: gel code blue stain, and Right: Fluorescence. MW = molecular weight in kDa. Lyse = lysate, FT = flow through. Excitation wavelength of Acd is 350 nm for fluorescence gel imaging.

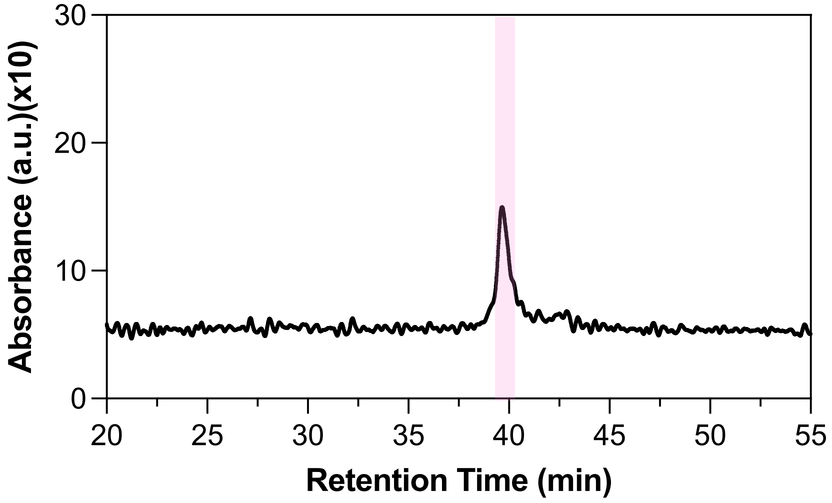


**Figure S18**. Analytical HPLC chromatogram showing the purity of LexA$\tau_{86}\delta_{175}$ (K156A). Peak absorption measured at 215 nm.

**Purification and Characterization of LexA$\tau_{86}\delta_{175}$ (QM-$L_{89}$P, $Q_{92}$W, $E_{152}$A, and $K_{156}$A)**

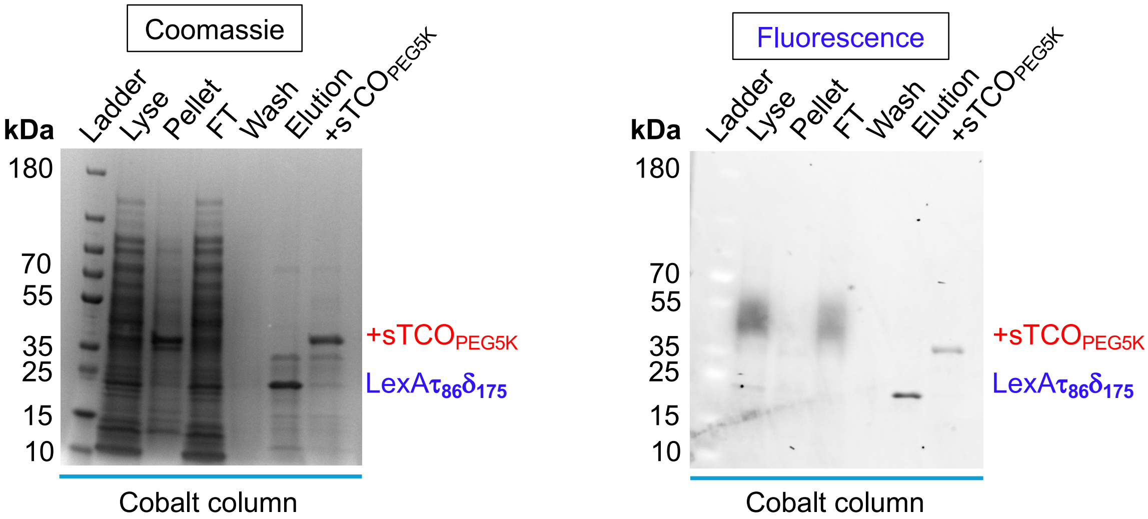


**Figure S19**. SDS-PAGE analysis of expression of LexA$\tau_{86}\delta_{175}$ (QM) at 30 ˚C. Left: gel code blue stain, and Right: Fluorescence. MW = molecular weight in kDa. Lyse = lysate, FT = flow through. Excitation wavelength of Acd is 350 nm for fluorescence gel imaging.

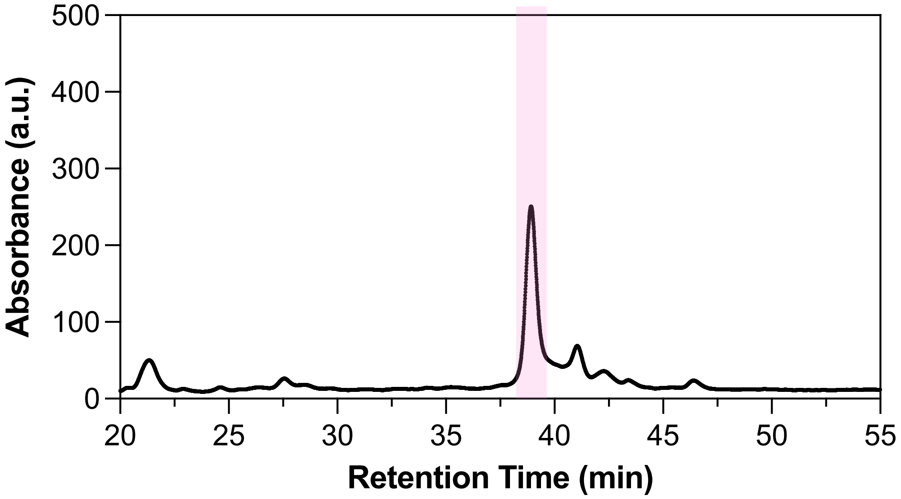


**Figure S20**. Analytical HPLC chromatogram showing the purity of LexA$\tau_{86}\delta_{175}$ (QM). Peak absorption measured at 215 nm.

**Purification of LexAδ175 (K156A) and LexAδ175(QM)**

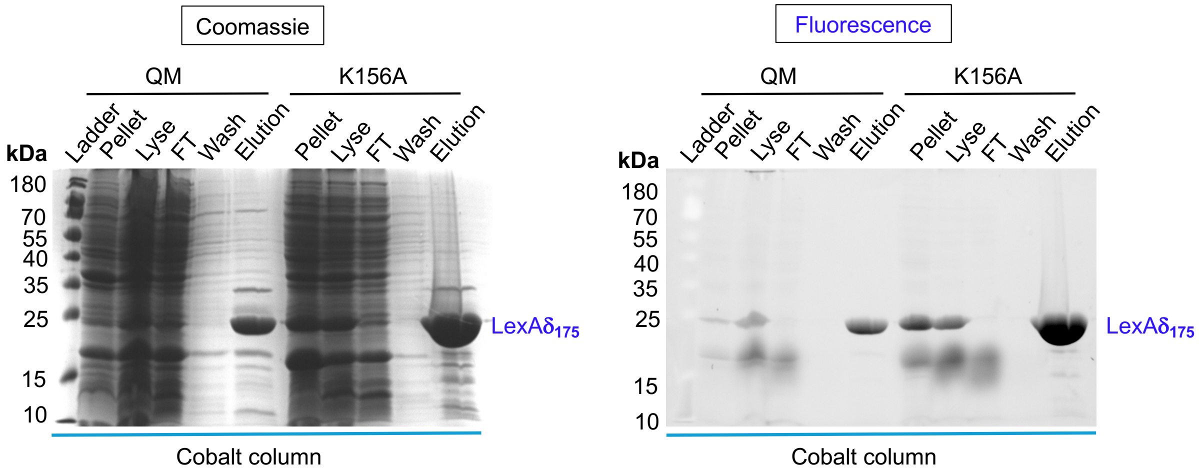


**Figure S21**. SDS-PAGE analysis of expression of LexA$\delta_{175}$ (QM) and LexA$\delta_{175}$ ($K_{156}$A) at 30 ˚C. Left: gel code blue stain, and Right: Fluorescence. MW = molecular weight in kDa. Lyse = lysate, FT = flow through. Excitation wavelength of Acd is 350 nm for fluorescence gel imaging.

**Oxidation of DihydroTet to Tet by Horseradish Peroxidase (HRP).** We prepared a 40 μM stock solution of HRP (Pierce™ Horseradish Peroxidase, Fisher Scientific, USA) in 100 mM phosphate buffer (pH 6.5) and stored it at -80 °C until further use. Separately, we prepared a 25 μM solution of LexA$\tau_{81}\delta_{86}$ in 100 mM phosphate buffer (pH 7.4), ensuring compatibility with HRP activity at pH 7.4. To these solutions, we added varying concentrations of HRP (0, 15 nM, and 1 μM) and incubated them at 37 °C with shaking at 250 rpm using an IKA MS3 control orbital shaker (Wilmington, NC, USA) for 20 minutes. After the incubation, Tet was reacted with either 1 equivalent or 2 equivalents of sTCO$_{PEG5K}$ (10 mM stock solution in $H_2O$). The reaction mixture was further incubated at 37 °C with shaking at 250 rpm for 5 minutes.
After 5 minutes of incubation, 5 μL of loading dye was added to 15 μL of each sample. The samples were then boiled for 5 minutes to denature the proteins. The denatured samples were loaded onto a 4% stacking and 12% resolving SDS-PAGE gel and run at 100–140 volts for 1 hour until the dye front reached the bottom of the gel. The gel was fixed in a destaining solution (40% (v/v) methanol and 10% (v/v) glacial acetic acid in Milli-Q water), followed by staining with GelCode Blue. The gel was subsequently destained with $H_2O$ to remove excess dye and imaged using a G:Box mini (Syngene).

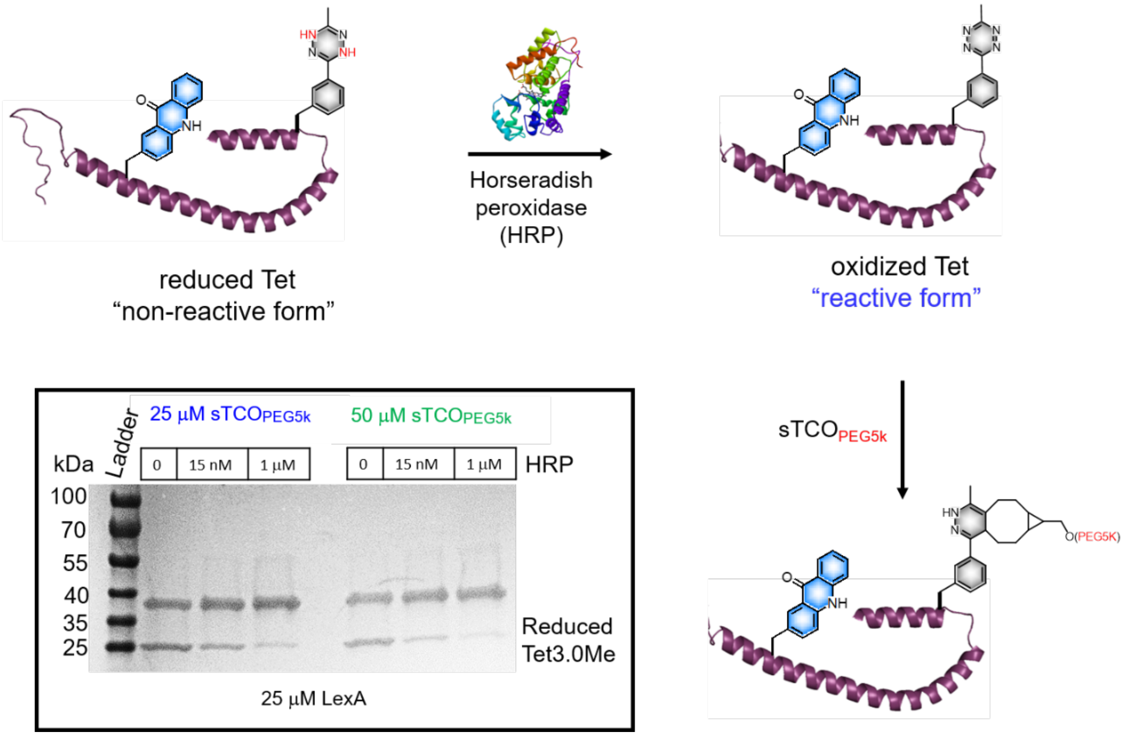


**Figure S22**. Schematic representation of Tet oxidation of LexA$\tau_{81}\delta_{86}$ (25 μM) by HRP and followed by gel shifting assay by labeling with the sTCO$_{PEG5K}$. Inset: SDS-PAGE of LexA$\tau_{81}\delta_{86}$ construct treated with varying concentrations of HRP (0, 15 nM, and 1 μM), followed by labeling with either 1 equiv. or 2 equiv. of sTCO$_{PEG5K}$.

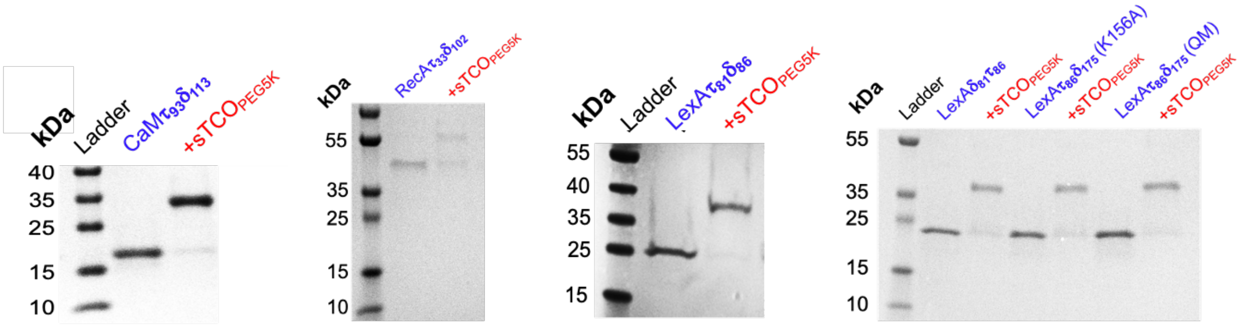


**Figure S23.** SDS-PAGE analysis of Tet IEDDA reaction of CaM, RecA, and LexA dual constructs with $sTCO_{PEG5k}$.

**Table S7. Single and Dual Incorporation of Tet and Acd in Proteins Expressed in *E. coli*.**

| Protein of Interest (POI) | Tet at TAA codon | Acd at TAG codon | Total Yield[a] (mg)/100 mL | Calculated Mass (m/z) | Observed Mass (m/z) |
|---|---|---|---|---|---|
| CaM | 93 | 113 | 3.5 (3.3)[b] | 16950.8 | 16959.0 |
| RecA | 33 | 102 | 2.0 | 39341.7 | 39336.5 |
| LexA | 81 | 86 | 2.4 | 23803.8 | 23804.1 |
| LexA | 86 | 81 | 2.5 | 23839.2 | 23837.6 |
| LexA (K156A) | 86 | 175 | 2.3 | 23810.1 | 23803.6 |
| LexA (QM) | 86 | 175 | 2.1 | 23794.1 | 23785.4 |
| LexA (K156A) | - | 175 | 2.8 | 23679.9 | 23670.7 |
| LexA (QM) | - | 175 | 3.0 | 23663.9 | 23654.3 |

[a]Total protein yield was determined using the DC assay. [b]Yield shown in parentheses corresponds to the pure protein; the difference from the total yield is attributed to cysteine adduct formation.

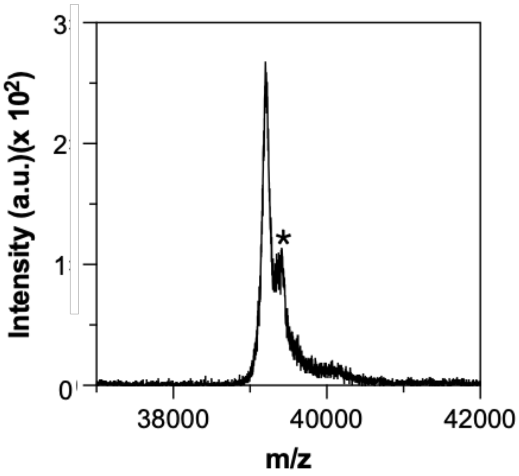


**Figure S24**. MALDI-TOF MS analysis of Tet and Acd incorporation in RecA. The data show masses corresponding to the successful incorporation of Tet and Acd. An asterisk (*) denotes a known matrix adduct.

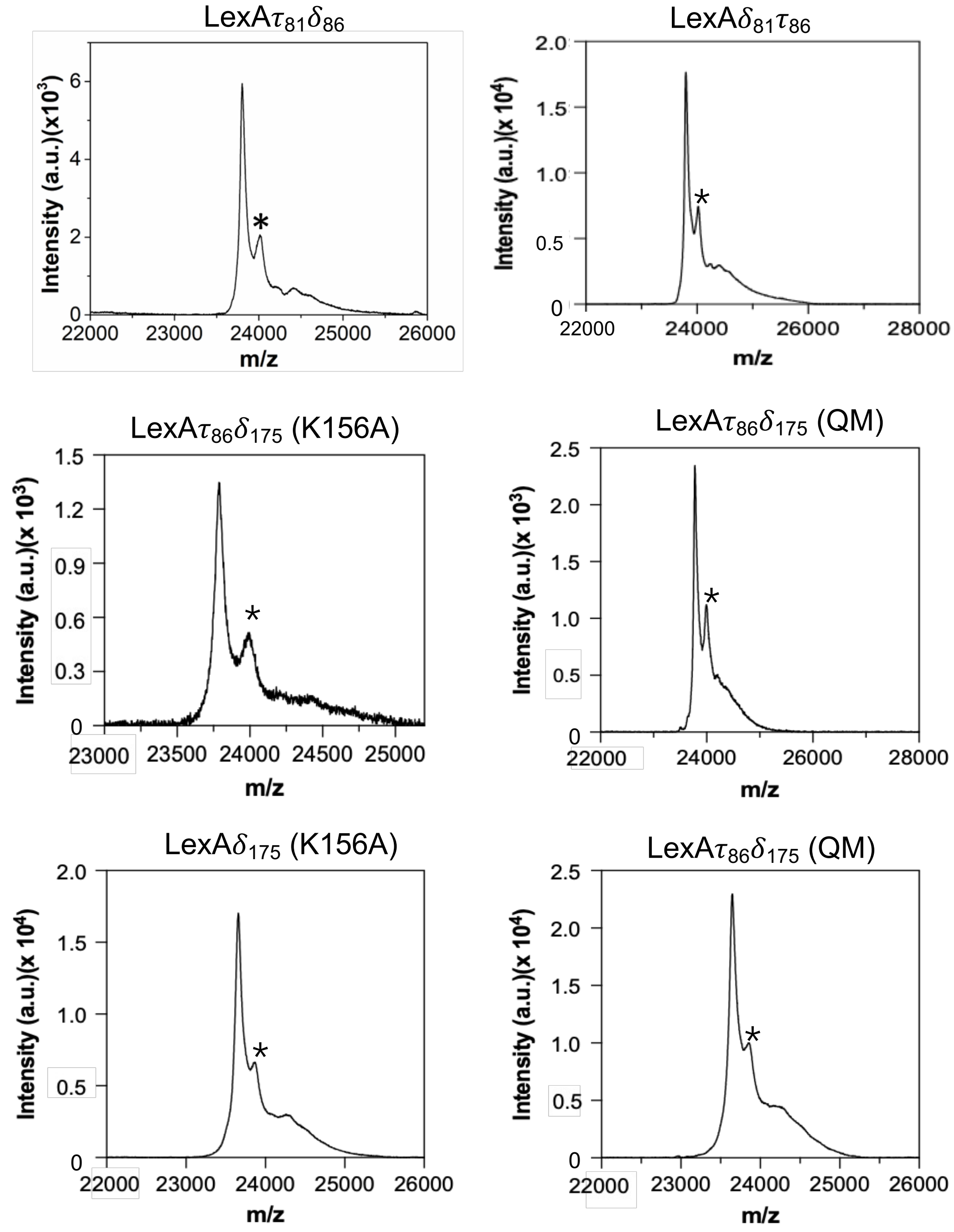


**Figure S25**. MALDI-TOF MS analysis of Tet and Acd incorporation in LexA. The data show masses corresponding to the successful incorporation of Tet and Acd. An asterisk (*) denotes a known matrix adduct.

### Conformational Changes in CaM Upon $Ca^{2+}$ Sensing and Tet Photolysis

**Fluorescence spectral measurements of CaM$\tau_{93}\delta_{113}$ with $Ca^{2+}$ ions.** The dual construct CaM$\tau_{93}\delta_{113}$ (25 μM) in 20 mM Tris buffer (pH 8.0) was titrated with varying concentrations of $Ca^{2+}$ (0, 0.5, 2.5, 7.5, 15, and 25 μM). The mixtures were mixed homogeneously by pipetting in Greiner 96-well flat black half-area plates at room temperature. Fluorescence spectra were recorded using a Tecan Spark plate reader (Männedorf, Switzerland) with an excitation wavelength ($\lambda_{ex}$) of 385 nm using the following parameters: excitation and emission bandwidths of 5 nm, delay time of 0 μs, and integration time of 40 μs.

Similarly, we measured fluorescence intensity to validate the conformational change. The dual construct CaM$\tau_{93}\delta_{113}$ (25 μM) in 20 mM Tris buffer (pH 8.0) was titrated by altering the order of addition of $Ca^{2+}$ (25 μM) and sTCO-OH (50 μM) via an IEDDA reaction. Fluorescence intensity was measured using a Tecan Spark plate reader (Männedorf, Switzerland) with an excitation wavelength ($\lambda_{ex}$) of 385 nm and an emission wavelength of 450 nm. The following parameters were used: excitation and emission bandwidths of 5 nm, delay time of 0 μs, and integration time of 40 μs.

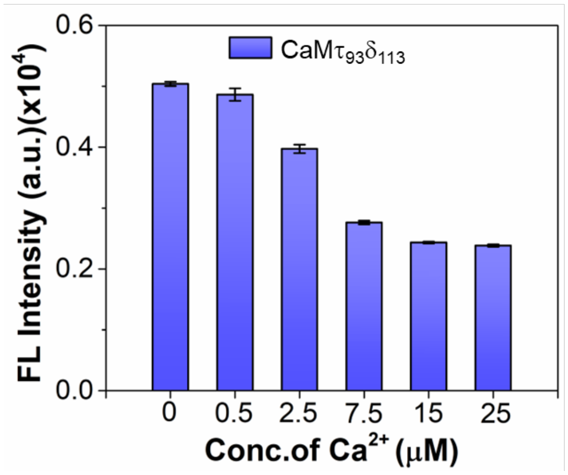


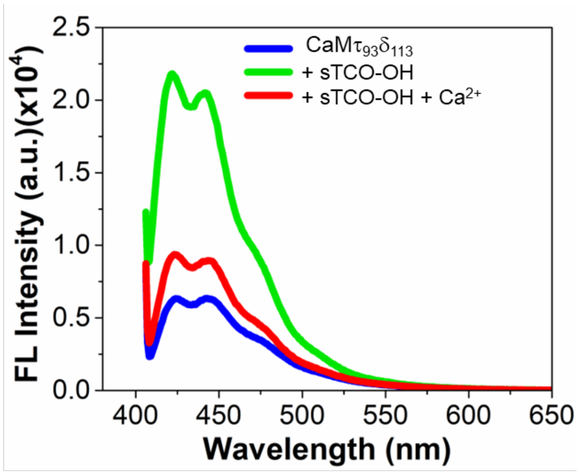


**Figure S26**. Left: Fluorescence spectral changes of CaM$\tau_{93}\delta_{113}$ upon titration with $Ca^{2+}$ ions in 20 mM Tris at pH 8.0, demonstrating the fluorescence response to calcium-induced conformational changes. Right: Validation of conformational changes through the retention of Acd fluorescence, analyzed by the addition of sTCO-OH (*via* the IEDDA reaction) and subsequent addition of $Ca^{2+}$ ions (25 μM) in 20 mM Tris at pH 8.0. Excitation and emission wavelengths for Acd were 385 nm and 450 nm, respectively. Error bars represent the SD of 3 measurements.

**Retention of Acd fluorescence by photolysis of Tet under 254 nm light irradiation.** The fluorescence intensity of the dual construct CaM$\tau_{93}\delta_{113}$ (25 µM) in Greiner 96-well flat black half-area plate was measured in the presence or absence of $Ca^{2+}$ in 20 mM Tris buffer (pH 8.0) at 0 minutes. The plate was then irradiated with a 254 nm light source (USHIO G8T5 low-pressure mercury-arc lamp, 7.2 watts) or 365 nm light source (ANALYTIK JENA, Mfr. No.34-0006-01, 8 watts) placed 5 cm away. Fluorescence intensity was recorded at different intervals of irradiation time.

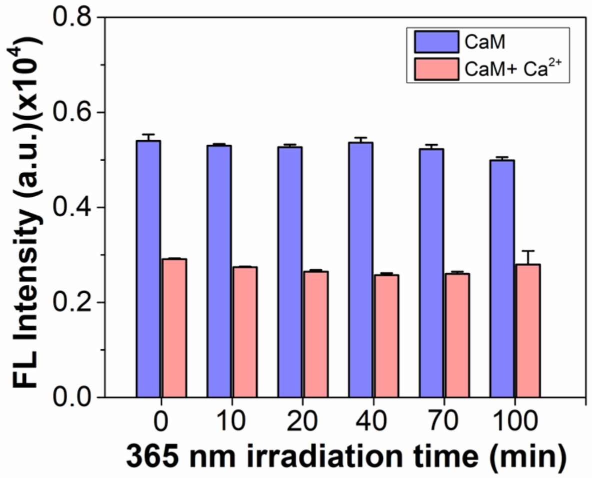


**Figure S27.** Fluorescence intensity changes during the photolysis of CaM$\tau_{93}\delta_{113}$ (25 µM) with and without $Ca^{2+}$ ions (25 µM) upon 365 nm light irradiation. Ex/Em wavelength of Acd is 385 nm/450 nm.

**LexA Cleavage**

**Alkaline-mediated and RecA*-mediated LexA auto-proteolysis assays.** LexA cleavage assays were performed essentially as previously described.[17] For the alkaline-mediated auto-proteolysis assay, 25 μM of His-LexA$\tau_{81}\delta_{86}$ was either mixed with an equal volume of RecA control buffer (70 mM Tris-HCl pH 7.5 or 8.5, 10 mM $MgCl_2$, 2 mM TCEP, 150 mM NaCl, 50 μg/mL BSA) or with an equal volume of alkaline pH reaction buffer (100 mM Tris-glycine-CAPS pH 10.6, 300 mM NaCl). Fluorescence intensity of Acd was monitored at 37 °C using Tecan Spark microplate reader with excitation wavelength at 385 nm and emission wavelength at 450 nm for 12 hours.

For the RecA*-mediated auto-proteolysis assay, RecA was activated by mixing His-RecA and 18-mer poly-GGT ssDNA (5'-GGT GGT GGT GGT GGT GGT-3' from Integrated DNA Technologies, Coralville, Iowa, USA.) in RecA activation buffer (70 mM Tris-HCl, pH 7.5, 10 mM MgCl2, 2 mM TCEP, 150 mM NaCl, 50 μg/mL BSA, and 0.25 mM ATPγS). The reactions were incubated for 3 hours at 25 °C. Activated RecA* was then added 1:1 with 25 μM of His-LexA$\tau_{81}\delta_{86}$, and the fluorescence intensity of Acd was monitored at 25 °C using Tecan Spark microplate reader. A non-activated RecA control was prepared similarly, with the exclusion of ATP-S.

Simultaneously, a gel-based assay was also used to examine His- LexA$\tau_{81}\delta_{86}$ auto-proteolysis in the presence of activated RecA (RecA*) or alkaline pH. At time zero, 5 μL RecA* and LexA were independently added to 10 μL of a stop mixture containing 2x Laemmli buffer (BioRad). For the other time points, 10 μL reaction mixture was added to 10 μL of 2x Laemmli buffer and boiled at 95 °C for 5 minutes. 20 μL of each sample was loaded onto a 15% SDS-PAGE gel, run at 150 V for 60 minutes, and stained with GelCode™ Blue Stain Reagent (Thermo Fisher) to visualize cleavage bands. Following gel image acquisition, protein bands were quantified using ImageJ (NIH). The percentage of cleaved LexA was calculated as the ratio of the C-terminal fragment band over the sum of the full-length and C-terminal fragment bands.

**Gel Analysis.** Samples were collected during cell lysis, nickel column purification, intein cleavage, and the second nickel column purification to confirm expression and monitor protein loss. For SDS-PAGE, 5 μL of loading dye was added to 5 μL of each sample, diluted to 20 μL with water, and boiled for 5 minutes to denature the proteins. The samples were run on a 4% stacking and 12% resolving gel at 60-140 volts for 1.5 hours until the dye front reached the gel bottom. The gel was fixed in a destaining solution (40% (v/v) methanol, 10% (v/v) glacial acetic acid in Milli-Q water) to remove non-covalently bound Acd or Tet. Fluorescent imaging for Acd-labeled proteins was performed using a G:Box mini (Syngene) before overnight staining with Gel Code Blue and subsequent destaining to remove excess dye. Representative gels for each protein construct are shown below.

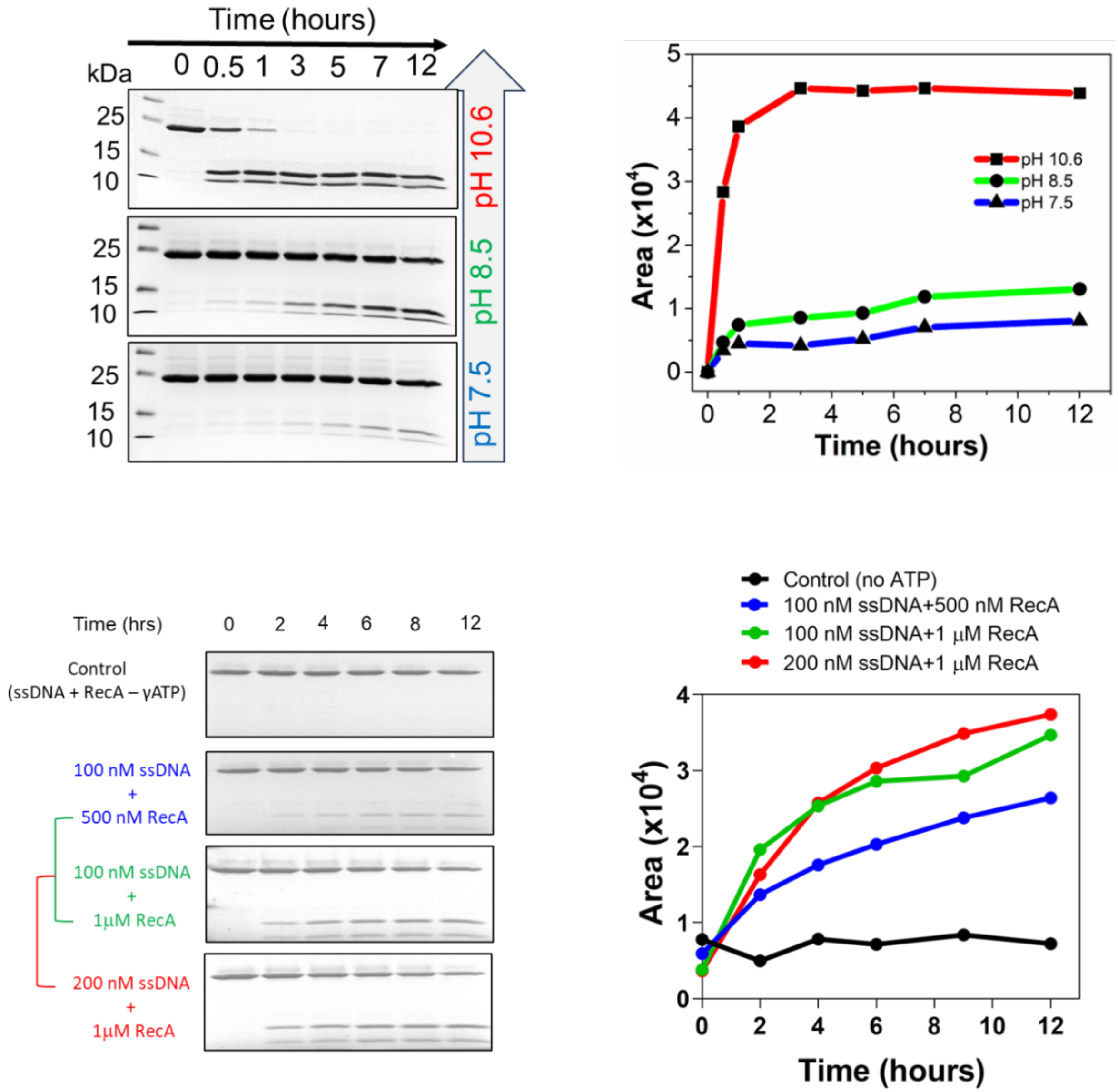


**Figure S28**. Top left: Quantitative validation of LexA (LexA$\tau_{81}\delta_{86}$) cleavage by SDS-PAGE gel analysis under varying pH conditions. Top right: pH-mediated LexA cleavage kinetics analyzed using ImageJ based on the gel bands of SDS-PAGE, correlating with fluorescence-based measurements. Bottom left: Quantitative validation of RecA* (with varying ssDNA or RecA concentrations) induced LexA cleavage by SDS-PAGE gel analysis. Bottom right: RecA*-mediated LexA (LexA$\tau_{81}\delta_{86}$) cleavage kinetics analyzed using ImageJ based on the gel bands of SDS-PAGE, correlating with fluorescence-based measurements.

**RecA activation Reactions**

A 2x concentrated reaction was prepared in activation buffer containing 70 mM Tris-HCl, pH 7.5, 10 mM $MgCl_2$, 2 mM TCEP, 150 mM NaCl, 50 μg/mL BSA, and 0.25 mM ATPγS with His-RecA and 18-mer poly-GGT ssDNA (5'-GGT GGT GGT GGT GGT GGT-3'. The reactions were incubated for 3 hours at 25 °C. For experiments involving concentration series of RecA*, the ssDNA substrate was maintained at a 4:1 nucleotide to RecA monomers. Non-activated RecA control was prepared similarly, with the exclusion of ATPγS.

**Fluorescence Anisotropy Measurements**

All stopped-flow anisotropy measurements were made in a KinTek AutoSF-120 stopped-flow spectrofluorometer, with the following settings. A Xenon/Mercury lamp (Hamamatsu) with a monochromator (Oriel) was set to emit light through a fiber optic at 385 nm with a 3.16 mm diffraction grating. Incident light was plane-polarized using a polarized optical fiber and passed into the observation chamber. Emitted light was filtered with two 440 +/− 40 nm emission filters (Edmund Optics) with polarized films in the vertical or horizontal direction for either photomultiplier tube (PMT). PMT voltages were calibrated, and a grating factor (G-factor) was obtained at the start of each instrument run.

Mixtures of LexA at 0.5 μM and RecA* at different concentrations (0.08, 0.32, 1.28, 5.12, and 15 μM) were monitored for changes in fluorescence anisotropy for 60 seconds. The means at all times for each curve were imported into KinTek Explorer global fitting software. Sigma calculation and normalization for each trace were performed using the built-in afit (analytical fit) and concentration scaling factor analysis, respectively, present in the software. Parameters in a one-step reversible binding model with LexA and RecA in a 1:1 stoichiometry were manually adjusted. The imported data were initially fitted by visual inspection, followed by refinement using the software's built-in regression tools. In brief, the model parameters were adjusted to minimize the $Chi^2$ value, defined as the sum of squared residuals normalized by the standard error (σ) at each time point.

$$\chi^2 = \sum_{i=0}^{N-1} \left( \frac{y_i - y(x_i)}{\sigma_i} \right)^2$$

Subsequently, the optimized parameters were systematically varied to identify ranges that still produced acceptable fits, based on $Chi^2$ analysis. This approach helps assess how well the data constrain the model. A $Chi^2$ threshold of 0.70 was chosen as a moderately strict cutoff to define upper and lower bounds for each parameter. At this threshold, most experimental data points fall within the predicted confidence bounds of the fitted curves upon visual inspection.

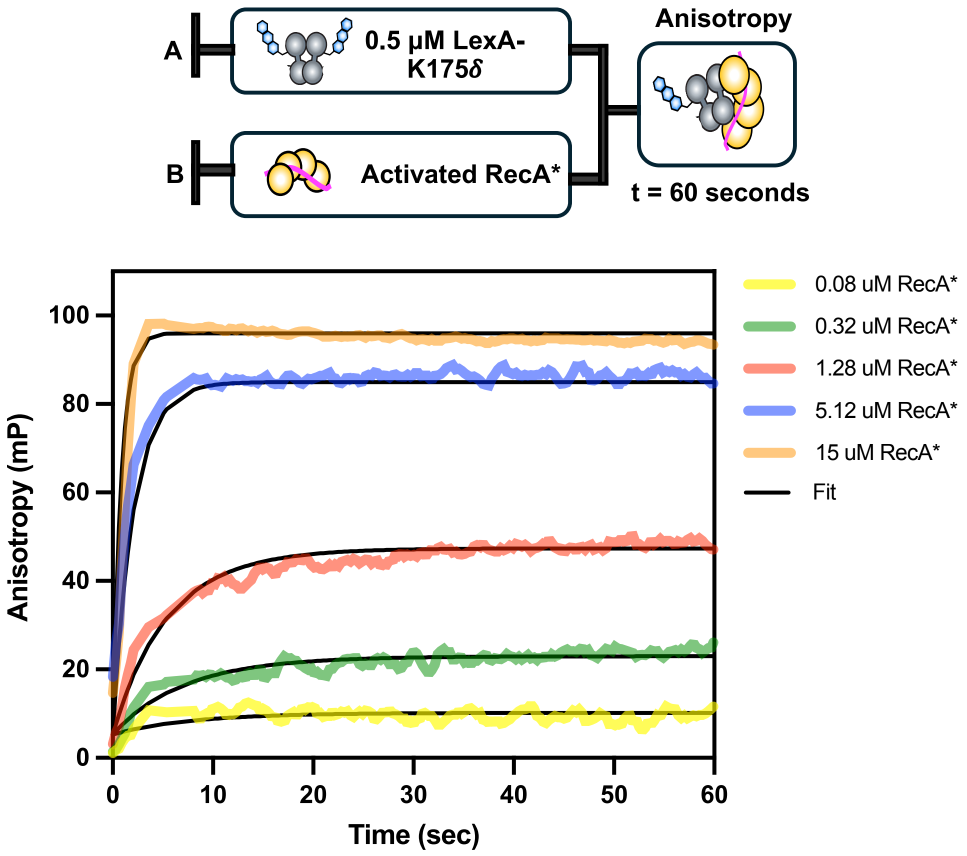


$$\mathrm{LexA} + \mathrm{RecA}^* \underset{0.079\ \mathrm{s}^{-1}}{\overset{0.084\ \mu\mathrm{M}^{-1}\mathrm{s}^{-1}}{\rightleftharpoons}} \mathrm{LexA}^*\!:\mathrm{RecA}^*$$

$$K_D = 0.94\ \mu\mathrm{M}$$

**Figure S29. Association kinetics of LexAδ$_{175}$ (K156A) with RecA**. 500 nM of LexAδ$_{175}$ (dimer) was rapidly mixed with a 1:4 ratio of RecA* and anisotropy was measured for 60 seconds. The best-fit curves from reaction simulations after globally fitting three independent concentration series experiments are shown as solid black lines for each concentration.

$$\mathrm{LexA} + \mathrm{RecA}^* \underset{\mathbf{k_{-1}}}{\overset{\mathbf{k_1}}{\rightleftharpoons}} \mathrm{LexA}^*\!:\mathrm{RecA}^*$$

$$\mathrm{FP} = \frac{\mathbf{a} \times [\mathrm{LexA}] + \mathbf{b} \times [\mathrm{LexA}^*\!:\mathrm{RecA}^*]}{[\mathrm{LexA}] + [\mathrm{LexA}^*\!:\mathrm{RecA}^*]}$$

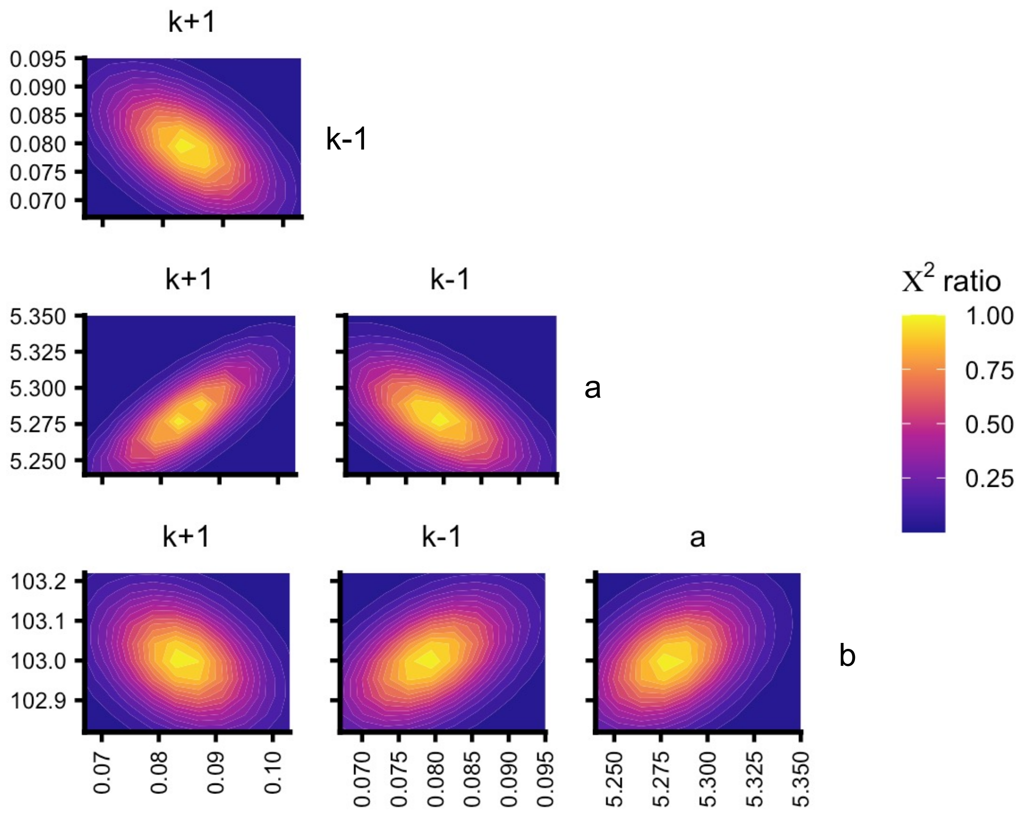


**Figure S30. One-step reversible binding model for LexA:RecA* interaction.** (Top) Fitted parameters include the forward and reverse rate constants, $k_1$ and $k_{-1}$, along with two factors, a and b, used to convert the protein concentrations to simulated fluorescence anisotropy signals. (Bottom) Two-dimensional FitSpace confidence intervals for these fitted parameters, with higher $Chi^2$ ratios corresponding to better fits.

**Table S8. Fitted values for reversible, one-step binding model parameters**

| **Parameter** | **Best fit value** | **Lower, Upper boundaries ($Chi^2$ threshold of 0.70)** |
|---|---|---|
| $k_1$ | 0.0842 | 0.079, 0.087 |
| $k_{-1}$ | 0.079 | 0.0763, 0.0825 |
| a | 5.28 | 5.26, 5.29 |
| b | 103 | 102.9, 103.1 |
| $C_1$ | 1 | N/A |
| $C_2$ | 0.97 | 0.96, 0.98 |
| $C_3$ | 0.95 | 0.94, 0.96 |

**Table S9. Summary of binding model fit results**

| Model characteristic | Result |
|---|---|
| Data points | 10105 |
| Degrees of Freedom (DoF) | 10098 |
| Sigma w.r.t. fit | 0.319 |
| Chi2 | 14644 |
| Chi2/DoF | 1.45 |
| p-value | 0 |
| Chi2 Threshold | 0.998116 |

**Fluorescence Anisotropy Measurements for LexA$\tau_{86}\delta_{175}$ (QM):**
Mixtures of LexA at 0.5 $\mu$M and RecA* at different concentrations (0, 0.08, 0.32, 1.28, 5.12, and 15 $\mu$M) were monitored for changes in fluorescence anisotropy for 60 seconds. Used the previous one-step reversible binding model for fitting the experimental data. Fits were optimized using built-in regression analysis in the software, similar to the previously mentioned fluorescence anisotropy data fitting.

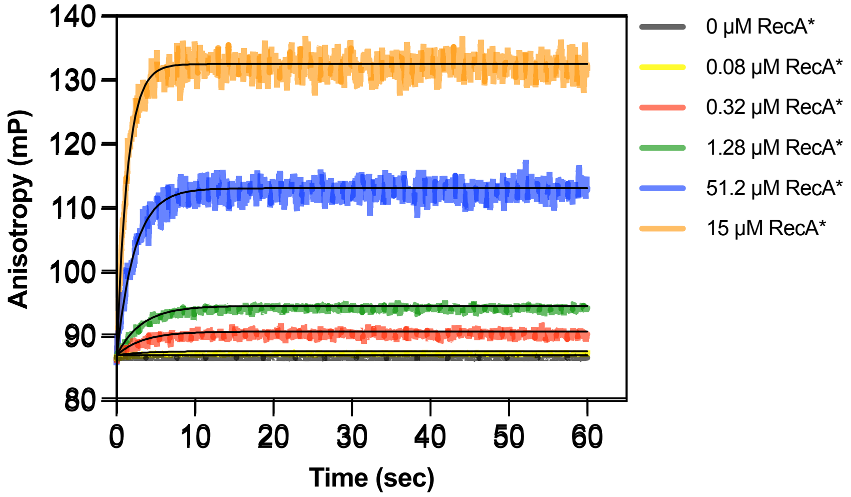


$$\text{LexA} + \text{RecA}^* \underset{0.269\ \text{s}^{-1}}{\overset{0.030\ \mu\text{M}^{-1}\text{s}^{-1}}{\rightleftharpoons}} \text{LexA}^*\text{: RecA}^*$$

$$K_D = 8.9\ \mu\text{M}$$

**Figure S31. Association kinetics of LexA$\delta_{175}$ (QM) with RecA**. 500 nM of LexA$\delta_{175}$ (dimer) was rapidly mixed with a 1:4 ratio of RecA*, and anisotropy was measured for 60 seconds. The best-fit curves from reaction simulations after globally fitting three independent concentration series experiments are shown as solid black lines for each concentration.

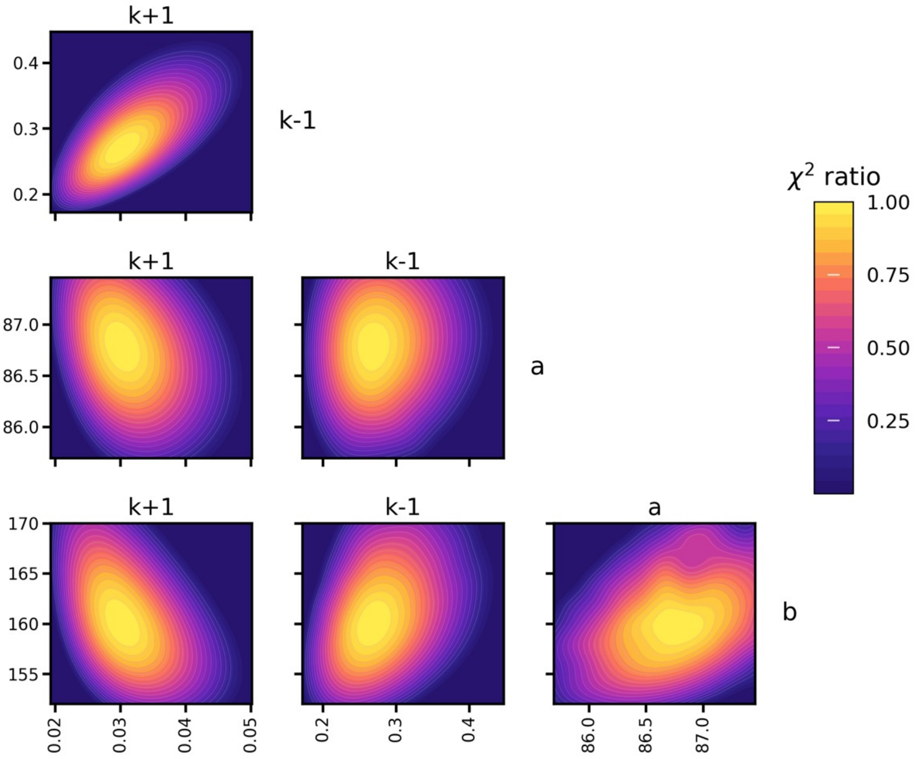


**Figure S32. One-step reversible binding model for LexA(QM):RecA* interaction.** (Top) Fitted parameters include the forward and reverse rate constants, $k_1$ and $k_{-1}$, along with two factors, a and b, used to convert the protein concentrations to simulated fluorescence anisotropy signals. (Bottom) Two-dimensional FitSpace confidence intervals for these fitted parameters, with higher $Chi^2$ ratios corresponding to better fits.

**Table S10. Fitted values for reversible, one-step binding model parameters**

| Parameter | Best fit value | Lower, Upper boundaries ($Chi^2$ threshold of 0.70) |
|---|---|---|
| $k_1$ | 0.030 | 0.024, 0.038 |
| $k_{-1}$ | 0.269 | 0.235, 0.313 |
| a | 86.780 | 86.300, 87.200 |
| b | 160.003 | 155.000, 166.000 |
| $C_1$ | 1 | N/A |
| $C_2$ | 0.98 | 0.97,0.99 |
| $C_3$ | 0.97 | 0.96, 0.98 |

**Table S11. Summary of binding model fit results**

| Model characteristic | Result |
|---|---|
| Data points | 24978 |
| Degrees of Freedom (DoF) | 24974 |
| Sigma w.r.t. fit | 0.143 |
| Chi2 | 35493 |
| Chi2/DoF | 1.42 |
| p-value | 0 |
| Chi2 Threshold | 0.999 |

**Fluorescence Quenching Measurements (LexA conformational change)**

All stopped-flow fluorescence intensity measurements were made in a KinTek AutoSF-120 stopped-flow spectrofluorometer, with the following settings. A Xenon/Mercury lamp (Hamamatsu) with a monochromator (Oriel) was set to emit light through a fiber optic at 385 nm with a 3.16 mm diffraction grating. Emitted light was filtered with a 440 +/− 40 nm emission filter (Edmund Optics) for either photomultiplier tube (PMT).

Mixtures of LexA$\tau_{86}\delta_{175}$ (K156A) at 1.5 $\mu$M and RecA* at different concentrations (1.5, 3, 5, 10, and 15 $\mu$M) were monitored for changes in fluorescence anisotropy for 60 seconds. A three-step kinetic model was used to fit the experimental data. Fits were optimized using built-in regression analysis in the software, similar to the previously mentioned fluorescence anisotropy data fitting.

$$\mathrm{LexA_1} \underset{\mathbf{k_{-1}}}{\overset{\mathbf{k_1}}{\rightleftharpoons}} \mathrm{LexA_2}$$

$$\mathrm{LexA_2} + \mathrm{RecA^*} \underset{\mathbf{k_{-2}}}{\overset{\mathbf{k_2}}{\rightleftharpoons}} \mathrm{LexA_2{:}\,RecA^*}$$

$$\mathrm{LexA_2{:}\,RecA^*} \underset{\mathbf{k_{-3}}}{\overset{\mathbf{k_3}}{\rightleftharpoons}} \mathrm{LexA^*{:}\,RecA^*}$$

$$\mathrm{F} = \frac{\mathbf{a} \times [\mathrm{LexA_1}] + \mathbf{b} \times [\mathrm{LexA_2} + \mathrm{LexA_2{:}\,RecA^*}] + \mathbf{c} \times [\mathrm{LexA^*{:}\,RecA^*}]}{[\mathrm{LexA_1}] + [\mathrm{LexA_2}] + [\mathrm{LexA_2{:}\,RecA^*}] + [\mathrm{LexA^*{:}\,RecA^*}]}$$

Here,
$LexA_1$ = Open state
$LexA_2$ = Closed state
LexA* = RecA* optimized closed state (Cleavable state)

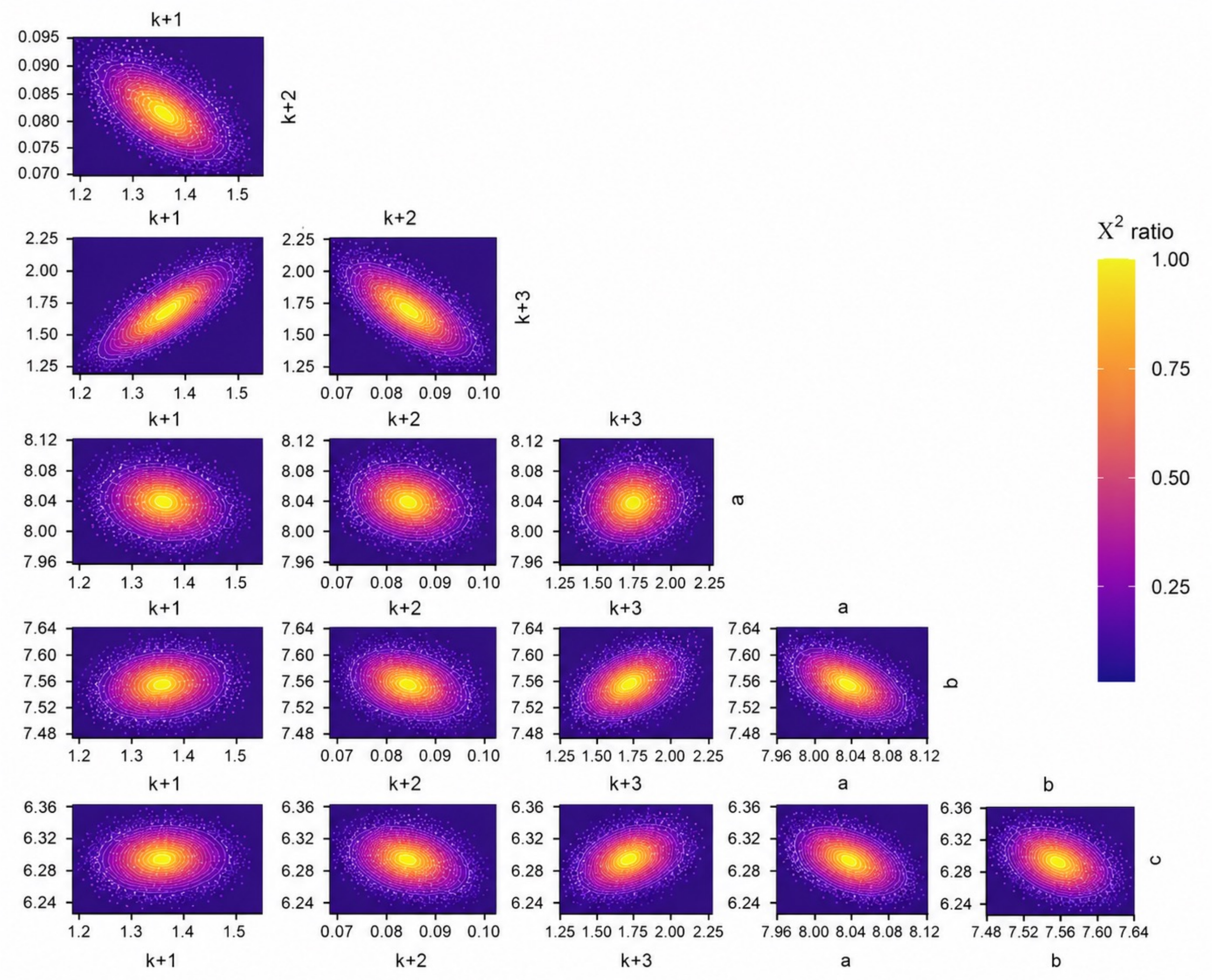


**Figure S33. Three-step hybrid model for LexA conformational change.** (Top) Fitted parameters include the forward equilibrium step, $k_1$, the RecA* binding step, $k_2$, and the RecA* stabilization step, $k_3$, along with three factors, a, b and c, used to convert the protein concentrations to simulated fluorescence signals. (Bottom) Two-dimensional FitSpace confidence intervals for these fitted parameters, with higher $Chi^2$ ratios corresponding to better fits.

**Table S12. Fitted values for the three-step model parameters**

| Parameter | Best fit value | Lower, Upper boundaries ($Chi^2$ threshold of 0.70) |
|---|---|---|
| $k_1$ | 1.360 | 1.190, 1.560 |
| $k_{-1}$ | 0.203 (fixed) | N/A |
| $k_2$ | 0.086 | 0.078, 0.094 |
| $k_{-2}$ | 0.080 | N/A |
| $k_3$ | 1.856 | 1.520, 2.270 |
| $k_{-3}$ | 0.435 (fixed) | N/A |
| a | 8.049 | 7.970, 8.120 |
| b | 7.589 | 7.530, 7.630 |
| c | 6.309 | 6.250, 6.360 |
| $C_1$ | 1 | N/A |
| $C_2$ | 0.99 | 0.98,1.00 |
| $C_3$ | 0.96 | 0.95, 0.97 |

**Table S13. Summary of binding model fit results**

| Model characteristic | Result |
|---|---|
| Data points | 9995 |
| Degrees of Freedom (DoF) | 9991 |
| Sigma w.r.t. fit | 0.019 |
| Chi2 | 12491 |
| Chi2/DoF | 1.250 |
| p-value | 0 |
| Chi2 Threshold | 0.999 |

**Alternative models tested for LexA conformational step**

**Model 1 (Induced Fit):**

$$\mathrm{LexA_1 + RecA^*} \underset{\mathbf{k_{-1}}}{\overset{\mathbf{k_1}}{\rightleftharpoons}} \mathrm{LexA_1{:}\,RecA^*}$$

$$\mathrm{LexA_1{:}\,RecA^*} \underset{\mathbf{k_{-2}}}{\overset{\mathbf{k_2}}{\rightleftharpoons}} \mathrm{LexA_2{:}\,RecA^*}$$

$$\mathrm{F} = \frac{\mathbf{a} \times [\mathrm{LexA_1 + LexA_1{:}\,RecA^*}] + \mathbf{b} \times [\mathrm{LexA_2{:}\,RecA^*}]}{[\mathrm{LexA_1}] + [\mathrm{LexA_2{:}\,RecA^*}] + [\mathrm{LexA_1{:}\,RecA^*}]}$$

Here,
$LexA_1$ = Open state
$LexA_2$ = Closed state

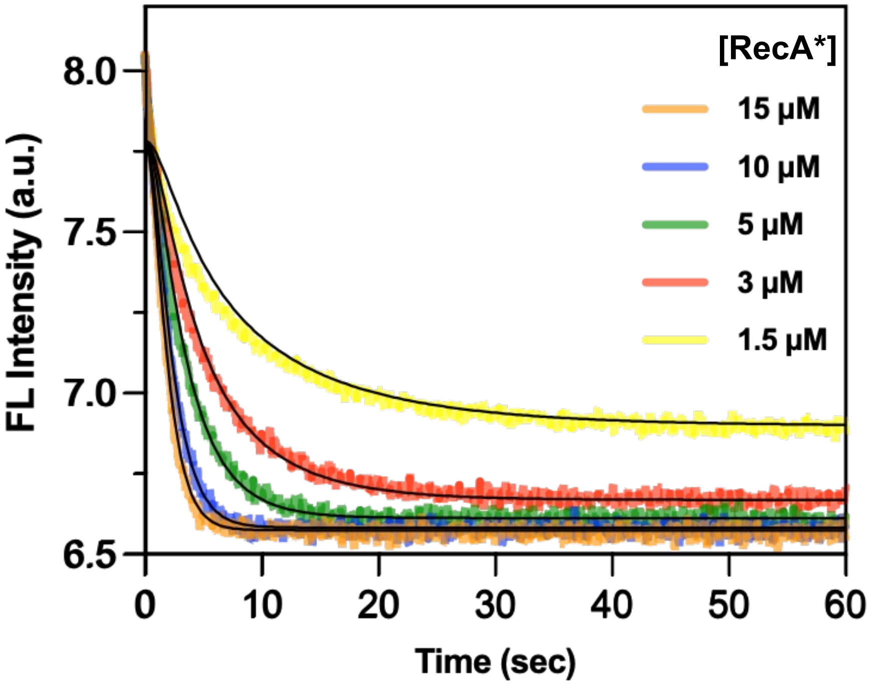


**Figure S34. Fitting to an induced fit model (1)**. (Top) Fitted parameters include the RecA* binding step, $k_1$, and the RecA* induced conformational change step, $k_2$, along with two factors, a and b used to convert the protein concentrations to simulated fluorescence signals. (Bottom) The best-fit curves from reaction simulations after globally fitting three independent concentration series experiments are shown as solid black lines for each concentration.

**Table S14. Fitted values for the two-step model parameters**

| Parameter | Best fit value | Lower, Upper boundaries ($Chi^2$ threshold of 0.70) |
|---|---|---|
| $k_1$ | 0.084 (fixed) | N/A |
| $k_{-1}$ | 0.016 | 0.013, 0.020 |
| $k_2$ | 0.141 | 0.121, 0.176 |
| $k_{-2}$ | 0.754 | 0.675, 0.921 |
| a | 7.783 | 7.730, 7.830 |
| b | $3.847 \times 10^{-6}$ | $3.850 \times 10^{-9}$, $8.850 \times 10^{-5}$ |

**Table S15. Summary of binding model fit results**

| Model characteristic | Result |
|---|---|
| Data points | 9995 |
| Degrees of Freedom (DoF) | 9992 |
| Sigma w.r.t. fit | 0.029 |
| Chi2 | 32567 |
| Chi2/DoF | 3.26 |
| p-value | 0 |
| Chi2 Threshold | 0.999 |

**Model 2 (Conformational Selection):**

$$\mathrm{LexA_1} \underset{\mathbf{k_{-1}}}{\overset{\mathbf{k_1}}{\rightleftharpoons}} \mathrm{LexA_2}$$

$$\mathrm{LexA_2} + \mathrm{RecA^*} \underset{\mathbf{k_{-2}}}{\overset{\mathbf{k_2}}{\rightleftharpoons}} \mathrm{LexA_2\!:RecA^*}$$

$$\mathrm{F} = \frac{\mathbf{a} \times [\mathrm{LexA_1}] + \mathbf{b} \times [\mathrm{LexA_2} + \mathrm{LexA_2\!:RecA^*}]}{[\mathrm{LexA_1}] + [\mathrm{LexA_2}] + [\mathrm{LexA_2\!:RecA^*}]}$$

Here,
$LexA_1$ = Open state
$LexA_2$ = Closed state

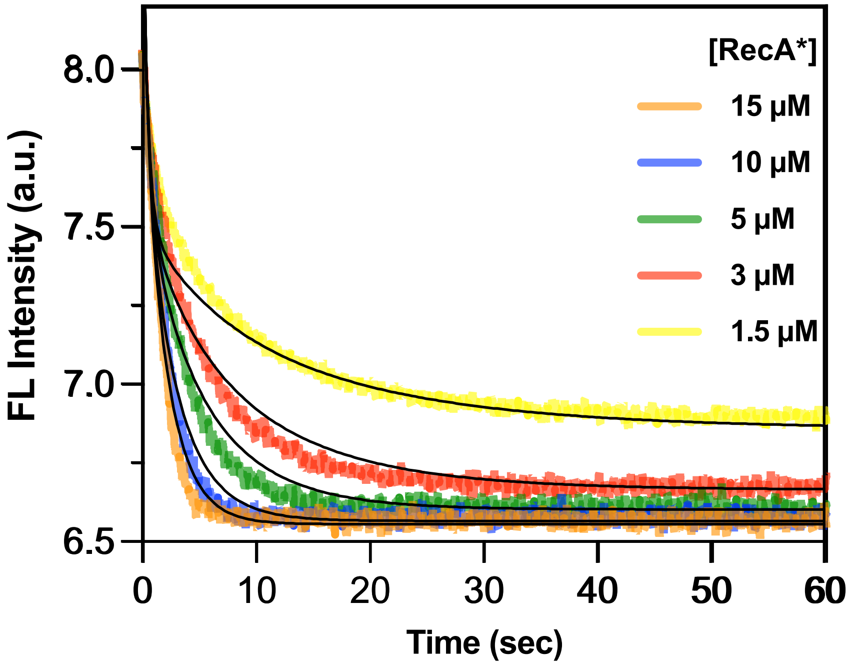


**Figure S35. Fitting to a conformational selection model (2)**. (Top) Fitted parameters include the forward equilibrium step, $k_1$, and the RecA* binding step, along with two factors, a and b used to convert the protein concentrations to simulated fluorescence signals. (Bottom) The best-fit curves from reaction simulations after globally fitting three independent concentration series experiments are shown as solid black lines for each concentration.

**Table S16. Fitted values for the two-step model parameters**

| Parameter | Best fit value | Lower, Upper boundaries (Chi$^2$ threshold of 0.70) |
|---|---|---|
| $k_1$ | 1.175 | 0.940, 1.840 |
| $k_{-1}$ | 1.071 | 0.857, 1.670 |
| $k_2$ | 0.084 (fixed) | N/A |
| $k_{-2}$ | 0.012 | 0.006, 0.019 |
| a | 8.482 | 8.340, 8.680 |
| b | 6.537 | 6.510, 6.560 |

**Table S17. Summary of binding model fit results**

| Model characteristic | Result |
|---|---|
| Data points | 9995 |
| Degrees of Freedom (DoF) | 9992 |
| Sigma w.r.t. fit | 0.040 |
| Chi2 | 61339 |
| Chi2/DoF | 6.14 |
| p-value | 0 |
| Chi2 Threshold | 0.999 |

**Model 3 (Not selected due to over-fitting of experimental data):**

$$\mathrm{LexA_1} \underset{\mathbf{k_{-1}}}{\overset{\mathbf{k_1}}{\rightleftharpoons}} \mathrm{LexA_2}$$

$$\mathrm{LexA_1} + \mathrm{RecA^*} \underset{\mathbf{k_{-2}}}{\overset{\mathbf{k_2}}{\rightleftharpoons}} \mathrm{LexA_1{:}\,RecA^*}$$

$$\mathrm{LexA_1{:}\,RecA^*} \underset{\mathbf{k_{-3}}}{\overset{\mathbf{k_3}}{\rightleftharpoons}} \mathrm{LexA_2{:}\,RecA^*}$$

$$\mathrm{LexA_2} + \mathrm{RecA^*} \underset{\mathbf{k_{-4}}}{\overset{\mathbf{k_4}}{\rightleftharpoons}} \mathrm{LexA_2{:}\,RecA^*}$$

$$F = \frac{\mathbf{a} \times [\text{LexA}_1 + \text{LexA}_1\text{: RecA}^*] + \mathbf{b} \times [\text{LexA}_2 + \text{LexA}_2\text{: RecA}^*]}{[\text{LexA}_1] + [\text{LexA}_2] + [\text{LexA}_2\text{: RecA}^*] + [\text{LexA}_1\text{: RecA}^*]}$$

Here,
$\text{LexA}_1$ = Open state
$\text{LexA}_2$ = Closed state

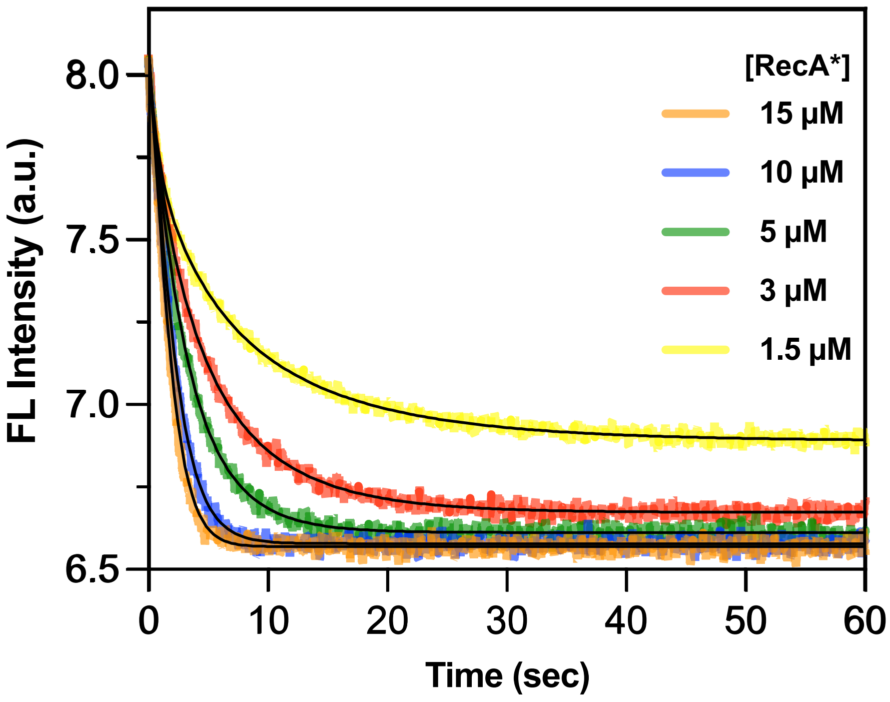


**Figure S36. Fitting to an alternate conformational selection + induced fit model (3)**. (Top) Fitted parameters include the forward equilibrium step, $k_1$, the RecA* binding step, $k_2$, the RecA* induced conformational change step, $k_3$, and the second RecA* binding step, $k_4$, along with two factors, a and b used to convert the protein concentrations to simulated fluorescence signals. (Bottom) The best-fit curves from reaction simulations after globally fitting three independent concentration series experiments are shown as solid black lines for each concentration.

**Table S18. Fitted values for the four-step model parameters**

| Parameter | Best fit value | Lower, Upper boundaries ($Chi^2$ threshold of 0.70) |
|---|---|---|
| $k_1$ | 0.338 | 0.044, 0.661 |
| $k_{-1}$ | 1.012 | 0.648, 1.980 |
| $k_2$ | 0.059 | 0.030, 0.099 |
| $k_{-2}$ | $8.618 \times 10^{-6}$ | $8.620 \times 10^{-9}$, $8.620 \times 10^{-5}$ |
| $k_3$ | 0.901 | 0.151, 1.760 |
| $k_{-3}$ | 0.074 | $1.360 \times 10^{-6}$, 0.816 |
| $k_4$ | 0.106 | 0.000, 0.691 |
| $k_{-4}$ | 0.016 | 0.013, 0.092 |
| a | 8.048 | 7.980, 8.140 |
| b | 6.430 | $5.360 \times 10^{-6}$, 6.56 |

**Table S19. Summary of binding model fit results**

| Model characteristic | Result |
|---|---|
| Data points | 9995 |
| Degrees of Freedom (DoF) | 9986 |
| Sigma w.r.t. fit | 0.015 |
| Chi2 | 6952 |
| Chi2/DoF | 0.69 |
| p-value | 1 |
| Chi2 Threshold | 0.999 |

**LexA Cleavage Kinetics Using Fluorescence Intensity Measurement**

All fluorescence intensity measurements were made using a Tecan Spark plate reader. Fluorescence intensity was measured at 450 nm with an excitation wavelength of 385 nm (5 nm bandwidth, 40 μs integration time, 31000 μm Z-position, and 3 sec linear shaking) using a nonsterile Greiner black, flat, 384-well low-volume microplate.

Mixtures of 0.5 $\mu$M LexA$\delta_{81}\tau_{86}$ and RecA* at different concentrations (0.25, 0.5, 1, 2.5, 5, 7.5, 15 $\mu$M) were monitored for changes in fluorescence intensity for 6 hours. The curves were imported into KinTek Explorer global fitting software. Sigma calculation and normalization for each trace were performed using the built-in afit (analytical fit) and concentration scaling factor analysis, respectively, present in the software. A three-step kinetic model was used to fit the experimental data. Fits were optimized using built-in regression analysis in the software, similar to the previously mentioned fluorescence anisotropy data fitting.

$$\mathrm{LexA} + \mathrm{RecA}^* \underset{\mathbf{k_{-1}}}{\overset{\mathbf{k_1}}{\rightleftharpoons}} \mathrm{LexA}^*\!:\mathrm{RecA}^*$$

$$\mathrm{LexA}^*\!:\mathrm{RecA}^* \xrightarrow{\mathbf{k_2}} \mathrm{LexA}^{**} + \mathrm{RecA}^*$$

$$\mathrm{F} = \frac{\mathbf{a} \times [\mathrm{LexA}] + \mathbf{b} \times [\mathrm{LexA}^{**}]}{[\mathrm{LexA}] + [\mathrm{LexA}^{**}]}$$

The following constraints were applied to better fit the experimental data to the above proposed model :

1. $k_1$ and $k_{-1}$ were fixed as 0.084 and 0.079, respectively
2. $k_{-2}$ fixed as 0

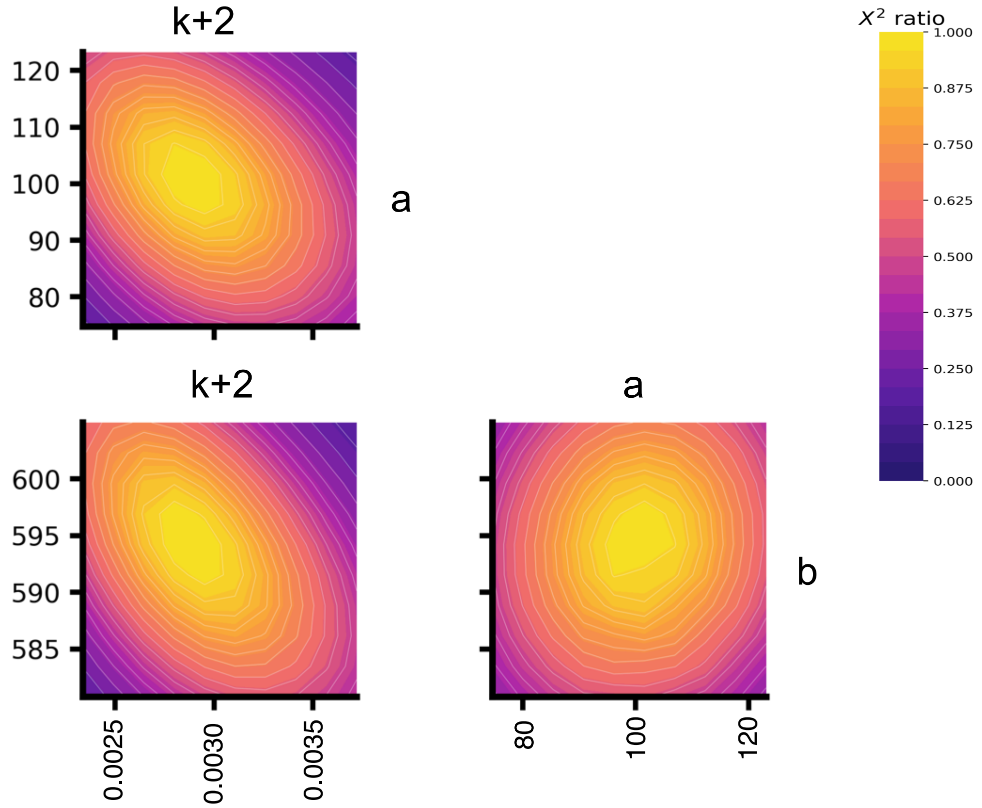


**Figure S37. Two-step model for LexA cleavage.** (Top) Fitted parameters include the forward rate constants for the autoproteolysis step ($k_2$), along with two factors (a and b) used to convert protein concentrations to simulated fluorescence signals. (Bottom) Two-dimensional FitSpace confidence intervals for these fitted parameters, with higher $Chi^2$ ratios corresponding to better fits.

**Table S20. Fitted values for two-step cleavage model parameters**

| Parameter | Best fit value | Lower, Upper boundaries ($Chi^2$ threshold of 0.70) |
|---|---|---|
| $k_1$ | 0.084 (fixed) | N/A |
| $k_{-1}$ | 0.079 (fixed) | N/A |
| $k_2$ | 0.0030 | 0.0027, 0.0031 |
| $k_{-2}$ | 0 (fixed) | N/A |
| a | 103.71 | 90.80, 111.00 |
| b | 595.64 | 589.00, 600.00 |
| $C_1$ | 1.00 | N/A |
| $C_2$ | 0.96 | 0.95, 0.97 |
| $C_3$ | 0.98 | 0.97, 0.99 |

**Table S21. Summary of binding model fit results**

| Model characteristic | Result |
|---|---|
| Data points | 1784 |
| Degrees of Freedom (DoF) | 1781 |
| Sigma w.r.t. fit | 0.1678 |
| Chi2 | 2713.19 |
| Chi2/DoF | 1.52 |
| p-value | 0 |
| Chi2 Threshold | 0.9956 |

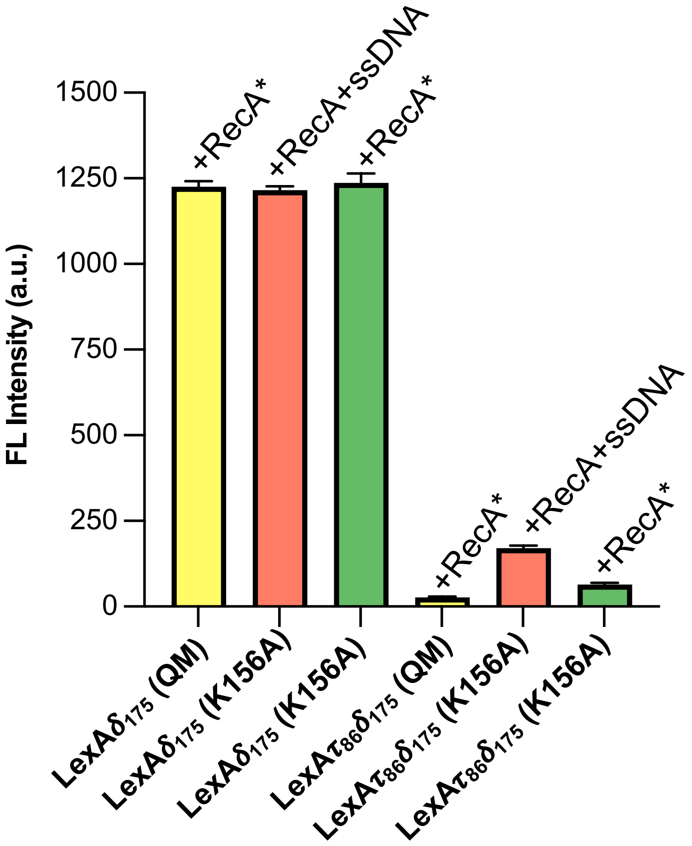


**Figure S38. Control experiment to identify the effect of surrounding amino acids on Acd fluorescence quenching.** Plot for changes in Acd (acridone) fluorescence intensity values on mixing activated RecA (RecA*) with single Acd LexA mutants, LexA$\delta_{175}$ (QM) and LexA$\delta_{175}$ (K156A), and dual Acd-Tet LexA mutants, LexA$\tau_{86}\delta_{175}$ (QM) and LexA$\tau_{86}\delta_{175}$ (K156A). Excitation and emission wavelengths for Acd were 385 nm and 450 nm, respectively. Error bars represent the standard deviation of three replicates.

**RecA*-Mediated LexA$\delta_{81}\tau_{86}$ FRET Biosensor Assay:**
RecA was activated by mixing 1 $\mu$M His-RecA and 2 $\mu$M 18-mer poly-GGT ssDNA (5’-GGT GGT GGT GGT GGT GGT-3’ from Integrated DNA Technologies, Coralville, Iowa, USA.) in RecA activation buffer (70 mM Tris-HCl, pH 7.5, 10 mM $MgCl_2$, 2 mM TCEP, 150 mM NaCl, 50 $\mu$g/mL BSA, and 0.25 mM ATP$\gamma$S). The reactions were incubated for 3 hours at 25 °C.
30 $\mu$M LexA$\delta_{81}\tau_{86}$ was pre-incubated for 10 minutes with 200 $\mu$M different small molecules (#1-20, Figure S41) before activated RecA* was then added 1:1 to give a final concentration of 15 $\mu$M LexA$\delta_{81}\tau_{86}$ and 100 $\mu$M of small molecule. The fluorescence intensity of Acd was monitored at 25 °C using a Tecan Spark microplate reader.

**Gel analysis.** After 18 hours from the time of addition of activated RecA * to LexA and small molecule mixes, 20 $\mu$L reaction mixture was added to 20 $\mu$L of 2x Laemmli buffer and boiled at 95 °C for 5 minutes. 40 $\mu$L of each sample was loaded onto a 15% SDS-PAGE gel, run at 150 V for 60 minutes, and stained with GelCode™ Blue Stain Reagent (Thermo Fisher) to visualize cleavage bands.

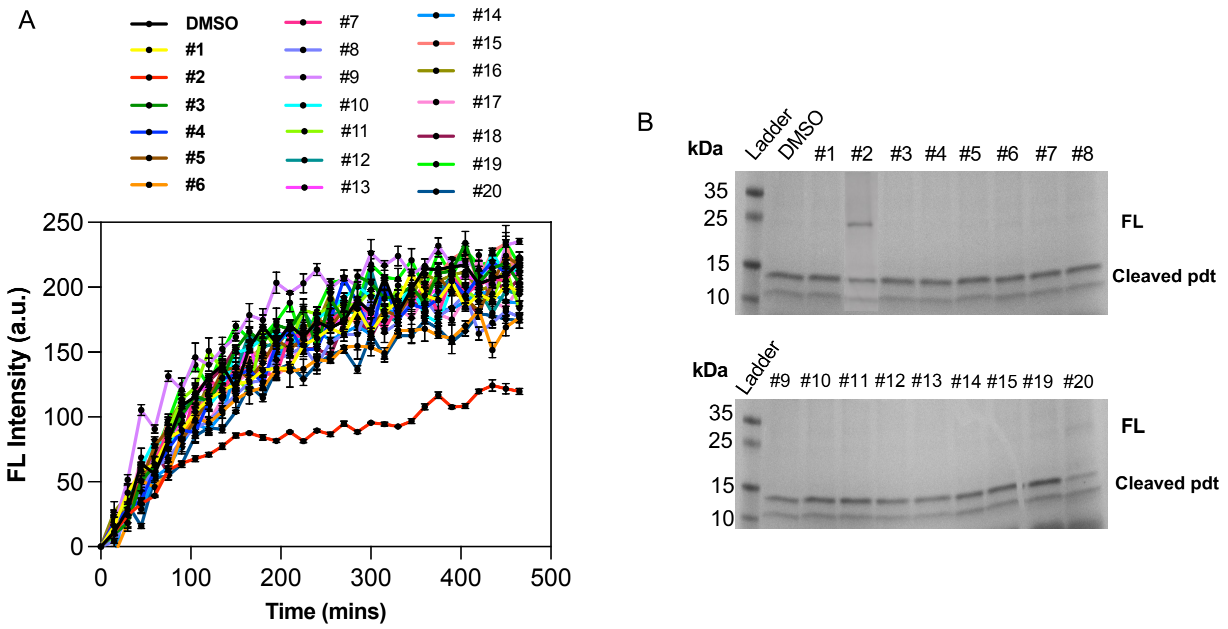


**Figure S39.** (A) Primary screening of small molecules using LexA$\delta_{81}\tau_{86}$ FRET biosensor on a 96-well plate. Error bars represent the standard deviation of three replicates. Excitation and emission wavelengths for Acd were 385 nm and 450 nm, respectively. (B) Gel-based LexA cleavage end point measurement after 18 hours from the addition of activated RecA*. FL=Full-length LexA

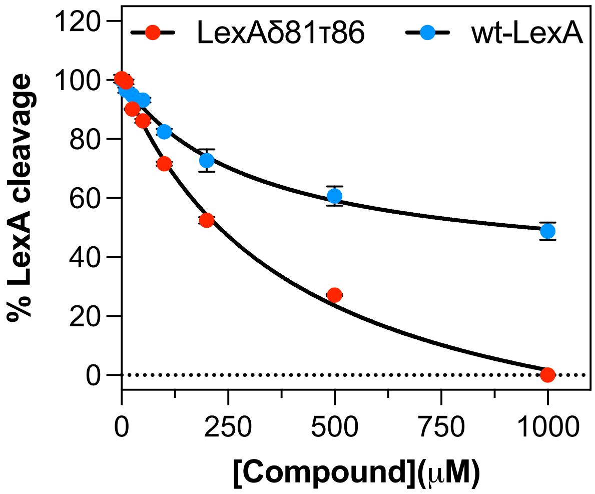


**Figure S40.** $IC_{50}$ curves for inhibition of LexA$\delta_{81}\tau_{86}$ and WT-LexA cleavage by compound **2** evaluated in the 96-well plate–based fluorescence assay (red) and gel-based assay (blue), respectively.

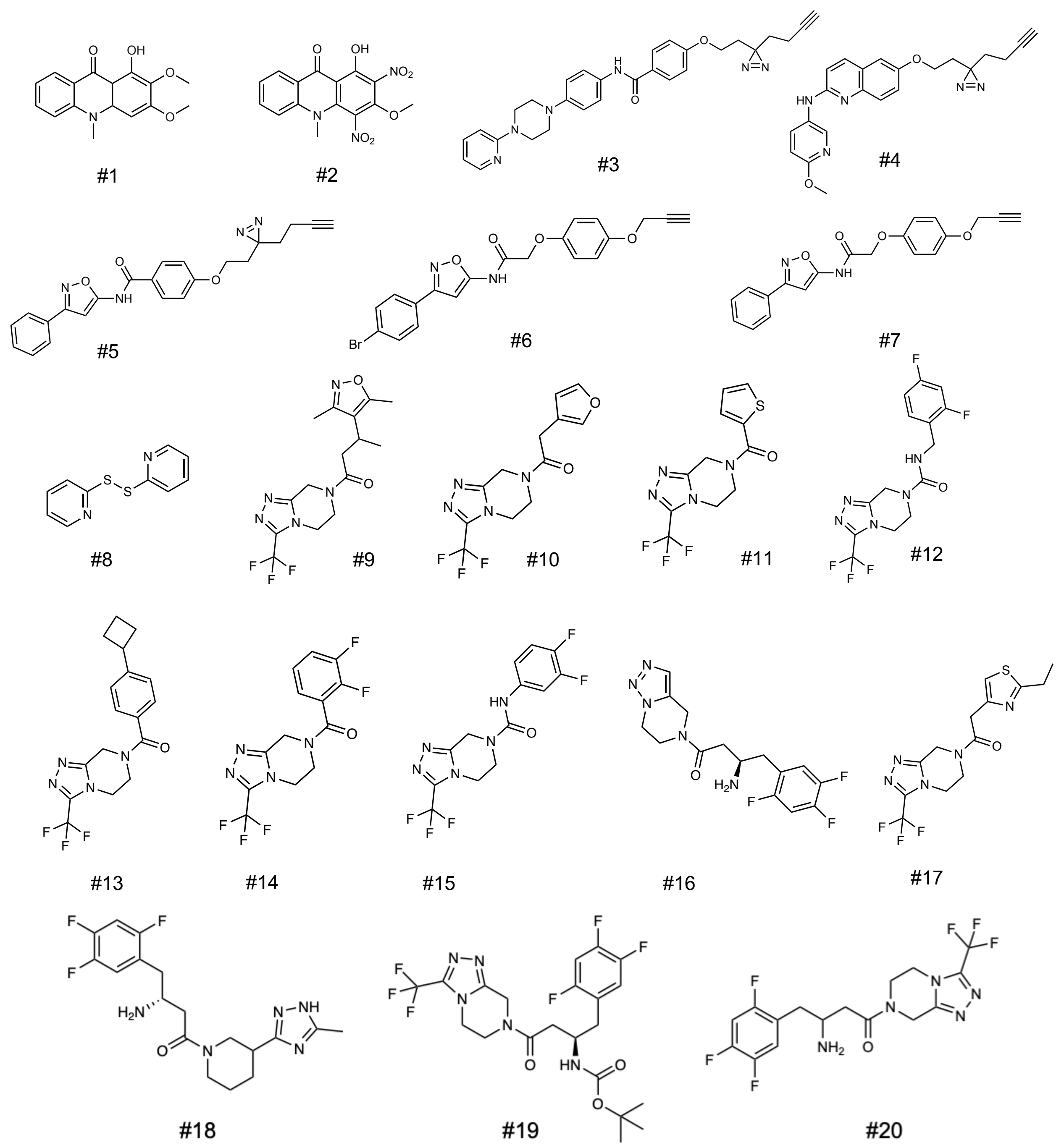


**Figure S41.** Chemical structures of small molecules screened against LexA$\delta_{81}\tau_{86}$ FRET biosensor.

**1-hydroxy-2,3-dimethoxy-10-methylacridin-9(10H)-one (1):** 1H NMR (600 MHz, CDCl3) δ 14.62 (s, 1H), 8.38 (d, J = 8.2 Hz, 1H), 7.76 – 7.69 (m, 1H), 7.47 (d, J = 8.7 Hz, 1H), 7.33 – 7.26 (m, 1H), 6.40 (s, 1H), 4.05 (s, 3H), 3.97 (s, 3H), 3.72 (s, 3H). 13C NMR (151 MHz, CDCl3) δ 181.9, 161.4, 160.1, 145.5, 138.6, 134.4, 129.1, 126.6, 121.7, 121.6, 115.7, 106.3, 93.9, 77.4, 77.2, 77.0, 61.7, 56.5, 40.7. HRMS (ESI$^+$) calcd for. $C_{16}H_{15}NO_4$ [M + H]$^+$, 286.1074; found: 286.1076

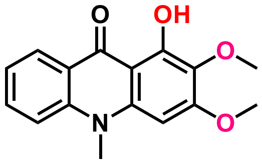


**1-hydroxy-3-methoxy-10-methyl-2,4-dinitroacridin-9(10H)-one (2):** 1H NMR (600 MHz, DMSO) δ 16.69 (s, 1H), 8.47 (d, J = 9.7 Hz, 1H), 7.92 (t, J = 7.9 Hz, 1H), 7.58 (d, J = 8.6 Hz, 1H), 7.54 (t, J = 8.0 Hz, 1H), 7.26 (s, 2H), 4.16 (s, 3H), 3.77 (s, 3H). 13C NMR (151 MHz, DMSO) δ 181.6, 161.0, 154.0, 143.9, 140.0, 136.4, 127.2, 126.9, 126.8, 125.0, 121.7, 117.0, 106.8, 65.6, 39.7. HRMS (ESI$^+$) calcd for. $C_{15}H_{11}N_3O_7$ [M + H]$^+$, 346.0670; found: 346.0670.

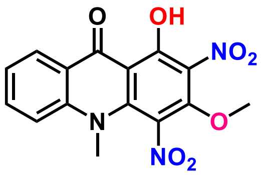

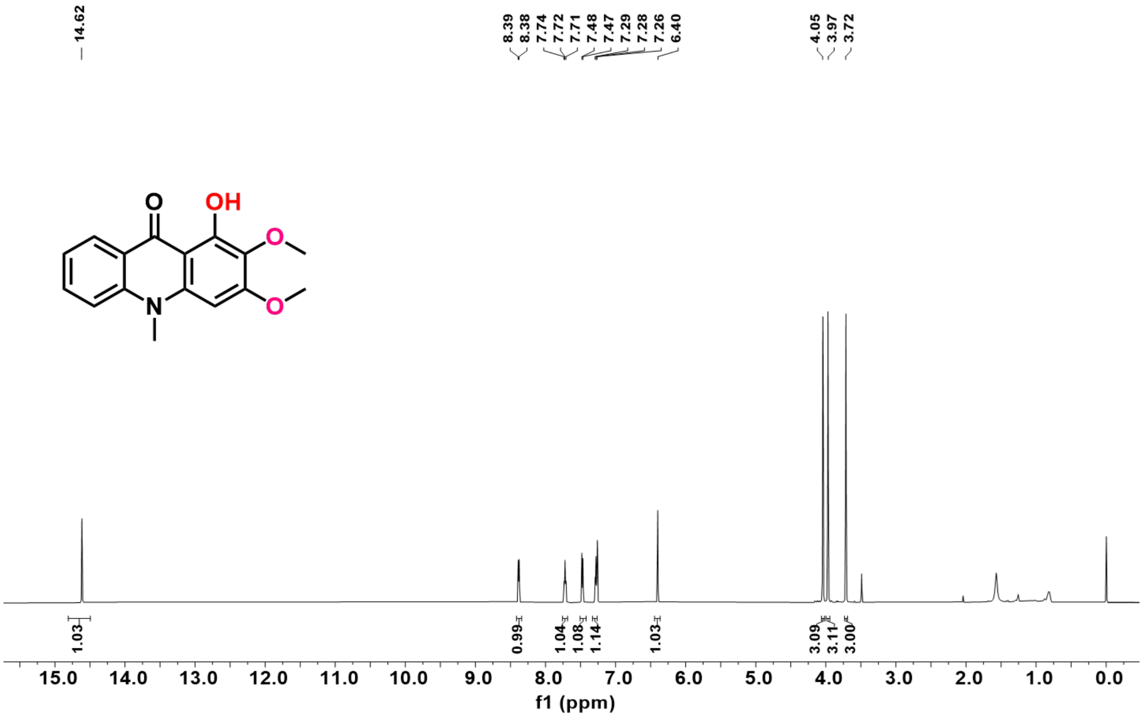


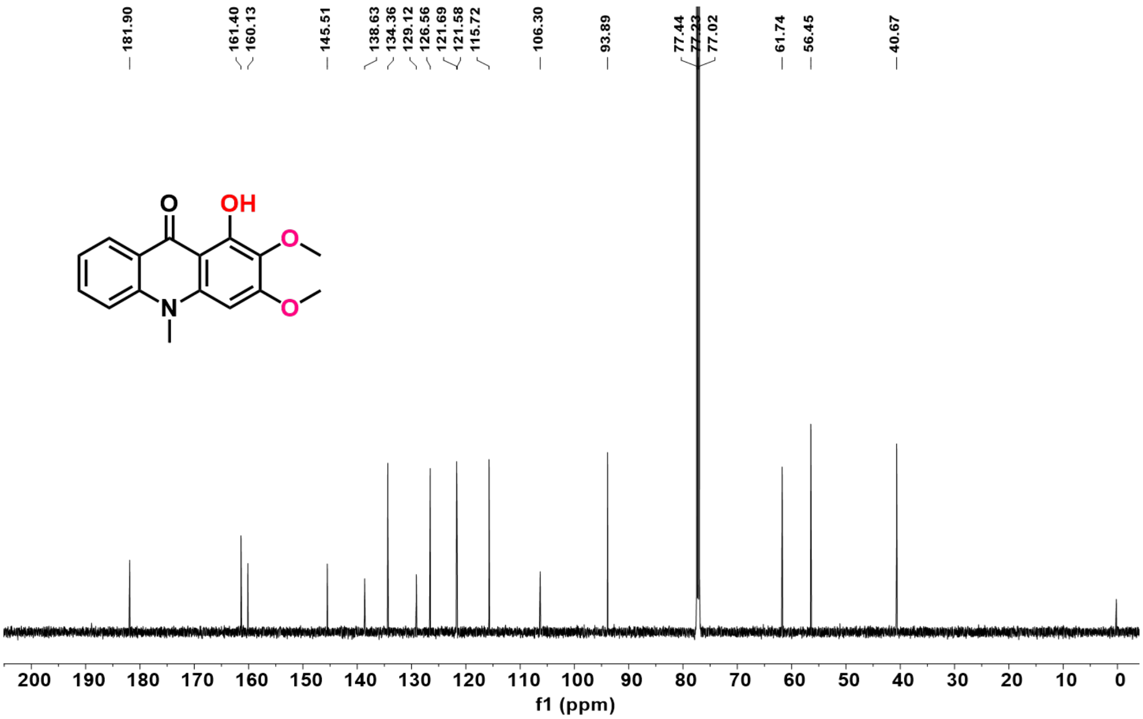


**Figure S42.** $^{1}H$ and $^{13}C$ NMR of the compound **1** in $CDCl_3$.

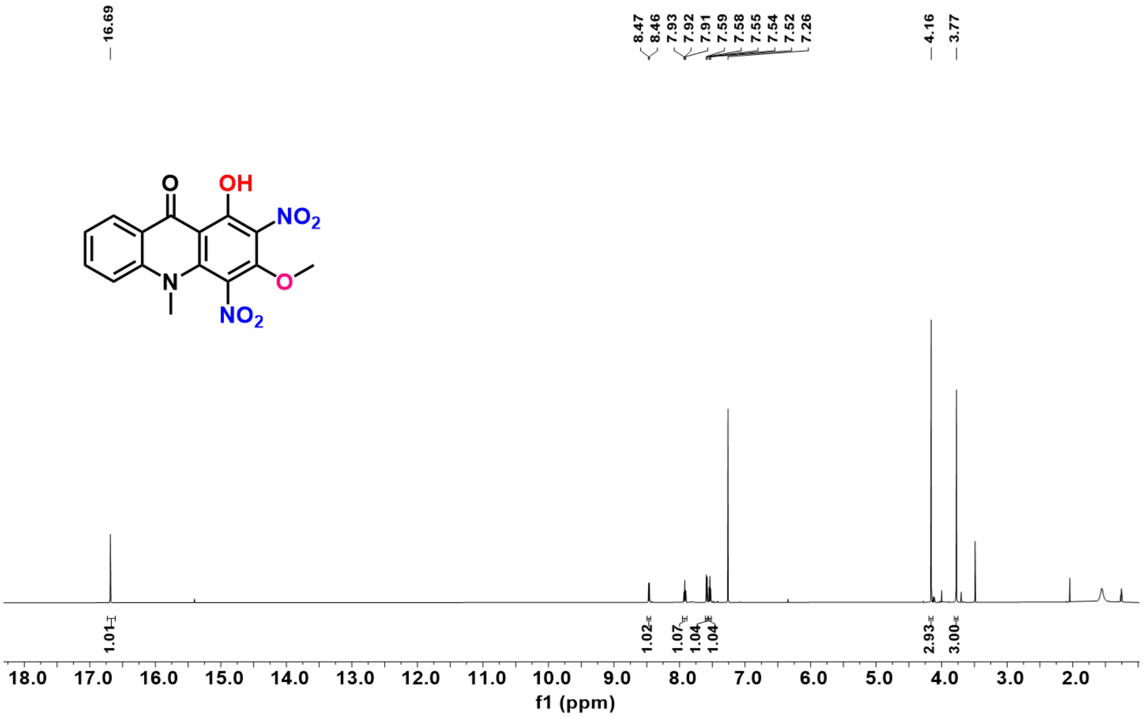

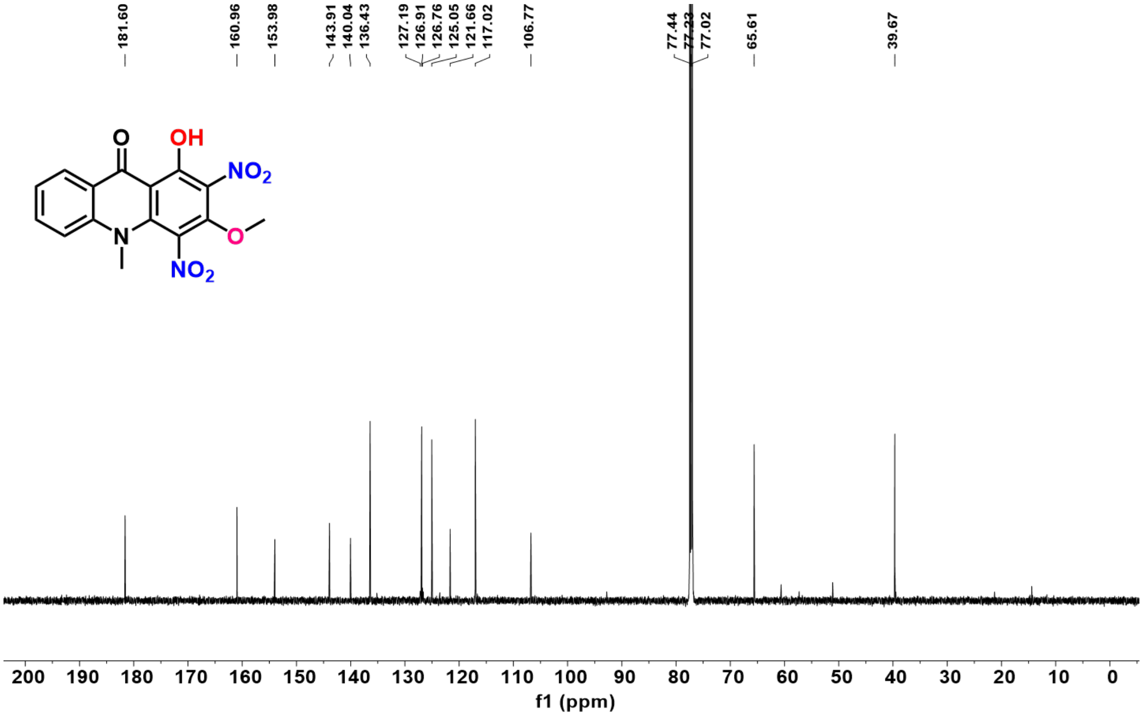

**Figure S43.** $^{1}H$ and $^{13}C$ NMR of the compound **2** in $CDCl_3$.

**X-ray Structure Determination of Compound 1**

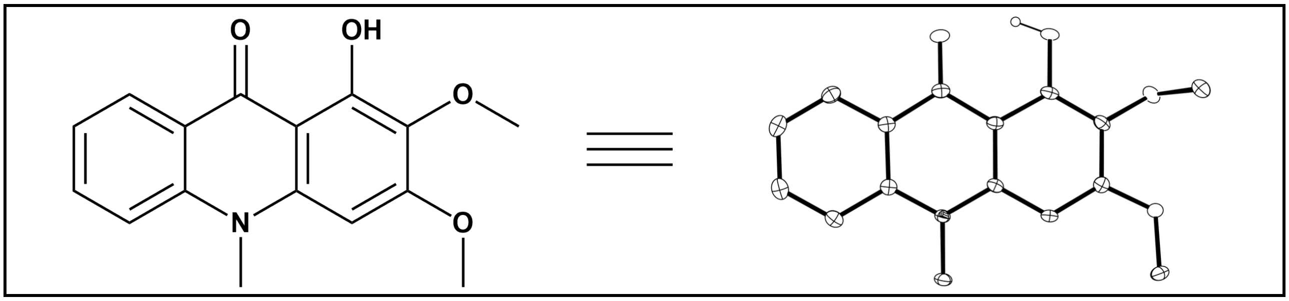


Compound **1**, $C_{16}H_{15}NO_4$, crystallizes in the orthorhombic space group Pbca (systematic absences hk0: h=odd, 0kl: k=odd, h0l: l=odd) with a=14.6718(6)Å, b=7.3756(3)Å, c=23.5504(8)Å, α=90°, β=90°, γ=90°, V=2548.47(17)$Å^3$, Z=8, and $d_{calc}$=1.487 g/$cm^3$. X-ray intensity data were collected on a Rigaku XtaLAB Synergy-S diffractometer [1] equipped with an HPC area detector (Dectris Pilatus3 R 200K) and employing confocal multilayer optic-monochromated Mo-Kα radiation (λ=0.71073 Å) at a temperature of 100K. Preliminary indexing was performed from a series of thirty 0.5° rotation frames with exposures of 10 seconds. A total of 1130 frames (10 runs) were collected employing ω scans with a crystal to detector distance of 34.0 mm, rotation widths of 0.5° and exposures of 50 seconds.

Rotation frames were integrated using CrysAlisPro [2], producing a listing of unaveraged $F^2$ and $\sigma(F^2)$ values. A total of 45565 reflections were measured over the ranges $4.436 \leq 2\theta \leq 56.556°$, $-17 \leq h \leq 19$, $-9 \leq k \leq 9$, $-31 \leq l \leq 31$ yielding 3148 unique reflections ($R_{int}$ = 0.0550). The intensity data were corrected for Lorentz and polarization effects and for absorption using SCALE3 ABSPACK [3] (minimum and maximum transmission 0.48380, 1.00000). The structure was solved by dual space methods - SHELXT [4]. Refinement was by full-matrix least squares based on $F^2$ using SHELXL [5]. All reflections were used during refinement. The weighting scheme used was $w=1/[\sigma^2(F_o^2)+(0.0636P)^2+1.8077P]$ where $P=(F_o^2+2F_c^2)/3$. Non-hydrogen atoms were refined anisotropically and hydrogen atoms were refined using a riding model. Refinement converged to R1=0.0436 and wR2=0.1155 for 2713 observed reflections for which $F > 4\sigma(F)$ and R1=0.0509 and wR2=0.1201 and GOF =1.027 for all 3148 unique, non-zero reflections and 194 variables. The maximum Δ/σ in the final cycle of least squares was 0.001 and the two most prominent peaks in the final difference Fourier were +0.41 and -0.26 e/$Å^3$.

Table 1. lists cell information, data collection parameters, and refinement data. Final positional and equivalent isotropic thermal parameters are given in Tables 2. and 3. Anisotropic thermal parameters are in Table 4. Tables 5. and 6. list bond distances and bond angles. Figure 1. is an ORTEP representation of the molecule with 50% probability thermal ellipsoids displayed.

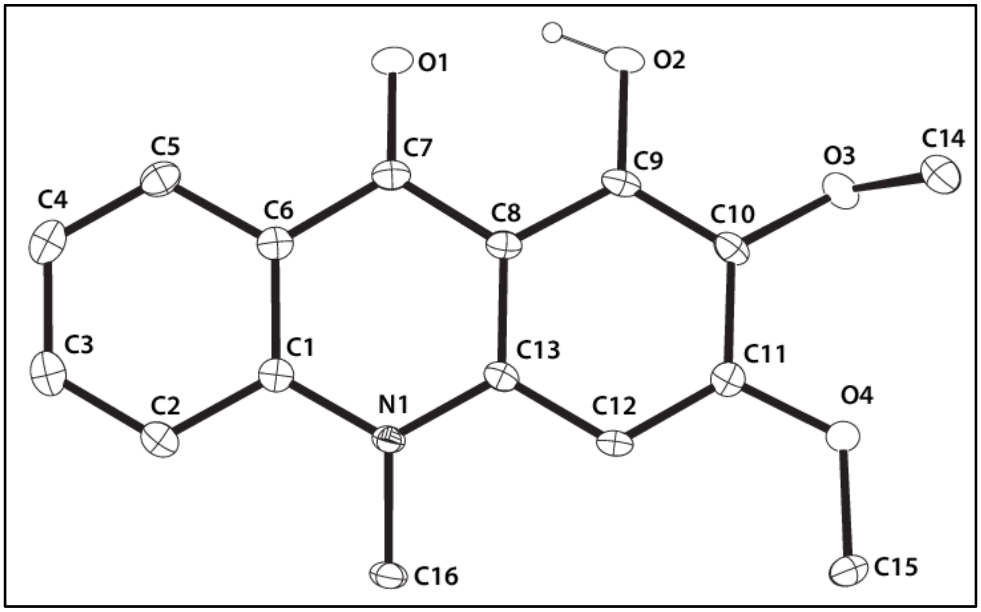


**Figure S44.** ORTEP drawing of the title compound **1** with 50% thermal ellipsoids.

**Table S22.** Summary of Structure Determination of Compound **1**

| | |
|---|---|
| Empirical formula | $C_{16}H_{15}NO_4$ |
| Formula weight | 285.29 |
| Diffractometer | Rigaku XtaLAB Synergy-S (Dectris Pilatus3 R 200K) |
| Temperature/K | 100(2) |
| Crystal system | orthorhombic |
| Space group | Pbca |
| a | 14.6718(6)Å |
| b | 7.3756(3)Å |
| c | 23.5504(8)Å |
| α | 90° |
| β | 90° |
| γ | 90° |
| Volume | 2548.47(17)$Å^3$ |
| Z | 8 |
| $d_{calc}$ | 1.487 g/$cm^3$ |
| μ | 0.108 $mm^{-1}$ |
| F(000) | 1200.0 |
| Crystal size, mm | 0.497 × 0.068 × 0.057 |
| 2θ range for data collection | 4.436 - 56.556° |
| Index ranges | -17 ≤ h ≤ 19, -9 ≤ k ≤ 9, -31 ≤ l ≤ 31 |
| Reflections collected | 45565 |
| Independent reflections | 3148[R(int) = 0.0550] |
| Data/restraints/parameters | 3148/0/194 |
| Goodness-of-fit on $F^2$ | 1.027 |
| Final R indexes [I>=2σ (I)] | $R_1$ = 0.0436, $wR_2$ = 0.1155 |
| Final R indexes [all data] | $R_1$ = 0.0509, $wR_2$ = 0.1201 |
| Largest diff. peak/hole | 0.41/-0.26 e$Å^{-3}$ |

**Table S23.** Refined Positional Parameters for Compound **1**.

| Atom | *x* | *y* | *z* | U(eq) |
|---|---|---|---|---|
| O1 | 0.45718(6) | 0.31790(14) | 0.53937(4) | 0.0174(2) |
| O2 | 0.49033(6) | 0.20360(14) | 0.63814(4) | 0.0179(2) |
| O3 | 0.42615(7) | 0.07127(14) | 0.73929(4) | 0.0175(2) |
| O4 | 0.24816(6) | 0.10849(14) | 0.76466(4) | 0.0167(2) |
| N1 | 0.18227(7) | 0.30781(15) | 0.57317(5) | 0.0120(2) |
| C1 | 0.21240(9) | 0.37955(17) | 0.52197(5) | 0.0123(3) |
| C2 | 0.15023(9) | 0.44433(18) | 0.48080(6) | 0.0146(3) |
| C3 | 0.18161(10) | 0.51609(19) | 0.43044(6) | 0.0169(3) |
| C4 | 0.27519(10) | 0.52620(19) | 0.41834(6) | 0.0178(3) |
| C5 | 0.33655(9) | 0.46440(18) | 0.45797(6) | 0.0156(3) |
| C6 | 0.30668(9) | 0.39129(17) | 0.51020(5) | 0.0128(3) |
| C7 | 0.37291(9) | 0.32348(17) | 0.55085(6) | 0.0129(3) |
| C8 | 0.33813(8) | 0.26664(17) | 0.60504(5) | 0.0122(3) |
| C9 | 0.39956(9) | 0.20823(18) | 0.64859(6) | 0.0135(3) |
| C10 | 0.36756(9) | 0.15172(18) | 0.70078(6) | 0.0142(3) |
| C11 | 0.27293(9) | 0.16063(18) | 0.71163(5) | 0.0132(3) |
| C12 | 0.21142(8) | 0.21613(17) | 0.67041(5) | 0.0124(3) |
| C13 | 0.24309(8) | 0.26443(17) | 0.61610(5) | 0.0113(3) |
| C14 | 0.44292(10) | 0.1712(2) | 0.79069(6) | 0.0211(3) |
| C15 | 0.15480(9) | 0.1363(2) | 0.78077(6) | 0.0207(3) |
| C16 | 0.08438(9) | 0.27124(19) | 0.58126(6) | 0.0147(3) |

**Table S24.** Positional Parameters for Hydrogens in Compound **1.**

| Atom | *x* | *y* | *z* | U(eq) |
|---|---|---|---|---|
| H2 | 0.500362 | 0.240016 | 0.604932 | 0.027 |
| H2A | 0.086559 | 0.438346 | 0.487897 | 0.018 |
| H3 | 0.138995 | 0.559676 | 0.403304 | 0.02 |
| H4 | 0.295783 | 0.575036 | 0.383286 | 0.021 |
| H5 | 0.399978 | 0.470947 | 0.450124 | 0.019 |
| H12 | 0.148151 | 0.22149 | 0.678829 | 0.015 |
| H14A | 0.478992 | 0.279473 | 0.781958 | 0.032 |
| H14B | 0.476499 | 0.094634 | 0.817514 | 0.032 |
| H14C | 0.3847 | 0.207486 | 0.807621 | 0.032 |
| H15A | 0.115411 | 0.057176 | 0.758073 | 0.031 |
| H15B | 0.138051 | 0.263152 | 0.774117 | 0.031 |
| H15C | 0.147239 | 0.107543 | 0.821126 | 0.031 |
| H16A | 0.076779 | 0.165814 | 0.606137 | 0.022 |
| H16B | 0.056028 | 0.246433 | 0.544387 | 0.022 |
| H16C | 0.055229 | 0.377156 | 0.598621 | 0.022 |

**Table S25.** Refined Thermal Parameters (U's) for Compound **1.**

| Atom | $U_{11}$ | $U_{22}$ | $U_{33}$ | $U_{23}$ | $U_{13}$ | $U_{12}$ |
|---|---|---|---|---|---|---|
| O1 | 0.0098(4) | 0.0227(5) | 0.0199(5) | 0.0001(4) | 0.0035(4) | -0.0012(4) |
| O2 | 0.0069(4) | 0.0282(6) | 0.0187(5) | -0.0003(4) | 0.0010(4) | 0.0007(4) |
| O3 | 0.0136(5) | 0.0214(5) | 0.0176(5) | 0.0005(4) | -0.0038(4) | 0.0040(4) |
| O4 | 0.0109(5) | 0.0248(5) | 0.0144(5) | 0.0042(4) | 0.0012(4) | 0.0024(4) |
| N1 | 0.0082(5) | 0.0148(5) | 0.0128(5) | -0.0006(4) | -0.0006(4) | -0.0005(4) |
| C1 | 0.0135(6) | 0.0099(6) | 0.0135(6) | -0.0026(4) | 0.0000(5) | -0.0002(4) |
| C2 | 0.0138(6) | 0.0137(6) | 0.0163(6) | -0.0027(5) | -0.0021(5) | 0.0015(5) |
| C3 | 0.0225(7) | 0.0133(6) | 0.0150(6) | -0.0018(5) | -0.0030(5) | 0.0028(5) |
| C4 | 0.0250(7) | 0.0142(6) | 0.0141(6) | -0.0001(5) | 0.0027(5) | -0.0020(5) |
| C5 | 0.0164(6) | 0.0137(6) | 0.0169(6) | -0.0018(5) | 0.0030(5) | -0.0023(5) |
| C6 | 0.0133(6) | 0.0113(6) | 0.0137(6) | -0.0031(5) | 0.0005(5) | -0.0008(5) |
| C7 | 0.0114(6) | 0.0109(6) | 0.0162(6) | -0.0035(5) | 0.0012(5) | -0.0015(4) |
| C8 | 0.0097(6) | 0.0123(6) | 0.0146(6) | -0.0024(5) | 0.0002(5) | -0.0009(4) |
| C9 | 0.0082(6) | 0.0145(6) | 0.0178(6) | -0.0035(5) | -0.0007(5) | 0.0008(5) |
| C10 | 0.0112(6) | 0.0155(6) | 0.0158(6) | -0.0009(5) | -0.0029(5) | 0.0025(5) |
| C11 | 0.0142(6) | 0.0118(6) | 0.0137(6) | -0.0007(5) | 0.0003(5) | 0.0006(5) |
| C12 | 0.0076(5) | 0.0137(6) | 0.0158(6) | -0.0008(5) | 0.0008(4) | 0.0005(4) |
| C13 | 0.0099(6) | 0.0094(6) | 0.0144(6) | -0.0023(4) | -0.0014(5) | -0.0001(4) |
| C14 | 0.0158(7) | 0.0296(8) | 0.0180(7) | -0.0010(6) | -0.0035(5) | -0.0012(6) |
| C15 | 0.0126(7) | 0.0312(8) | 0.0184(7) | 0.0073(6) | 0.0048(5) | 0.0029(6) |
| C16 | 0.0081(6) | 0.0195(7) | 0.0167(6) | -0.0007(5) | -0.0003(5) | -0.0005(5) |

**Table S26.** Bond Distances in Compound **1, Å**

| | | | | | |
|---|---|---|---|---|---|
| O1-C7 | 1.2663(16) | O2-C9 | 1.3547(15) | O3-C10 | 1.3834(16) |
| O3-C14 | 1.4383(17) | O4-C11 | 1.3563(16) | O4-C15 | 1.4361(16) |
| N1-C1 | 1.3891(16) | N1-C13 | 1.3859(16) | N1-C16 | 1.4737(16) |
| C1-C2 | 1.4143(18) | C1-C6 | 1.4134(17) | C2-C3 | 1.3780(19) |
| C3-C4 | 1.404(2) | C4-C5 | 1.375(2) | C5-C6 | 1.4128(18) |
| C6-C7 | 1.4528(18) | C7-C8 | 1.4371(18) | C8-C9 | 1.4317(18) |
| C8-C13 | 1.4186(17) | C9-C10 | 1.3801(19) | C10-C11 | 1.4133(18) |
| C11-C12 | 1.3872(18) | C12-C13 | 1.4067(17) | | |

**Table S27.** Bond Angles in Compound **1, °**

| | | | | | |
|---|---|---|---|---|---|
| C10-O3-C14 | 116.00(11) | C11-O4-C15 | 117.28(10) | C1-N1-C16 | 119.47(11) |
| C13-N1-C1 | 121.08(11) | C13-N1-C16 | 119.41(11) | N1-C1-C2 | 121.23(12) |
| N1-C1-C6 | 120.33(11) | C6-C1-C2 | 118.43(12) | C3-C2-C1 | 120.29(12) |
| C2-C3-C4 | 121.45(13) | C5-C4-C3 | 119.01(12) | C4-C5-C6 | 120.97(13) |
| C1-C6-C7 | 120.29(12) | C5-C6-C1 | 119.84(12) | C5-C6-C7 | 119.84(12) |
| O1-C7-C6 | 121.57(12) | O1-C7-C8 | 121.79(12) | C8-C7-C6 | 116.62(11) |
| C9-C8-C7 | 120.03(12) | C13-C8-C7 | 121.02(11) | C13-C8-C9 | 118.94(12) |
| O2-C9-C8 | 119.76(12) | O2-C9-C10 | 119.25(12) | C10-C9-C8 | 120.97(12) |
| O3-C10-C11 | 120.79(12) | C9-C10-O3 | 120.13(12) | C9-C10-C11 | 118.77(12) |
| O4-C11-C10 | 114.61(11) | O4-C11-C12 | 123.62(12) | C12-C11-C10 | 121.75(12) |
| C11-C12-C13 | 119.74(12) | N1-C13-C8 | 119.76(11) | N1-C13-C12 | 120.60(11) |
| C12-C13-C8 | 119.64(11) | | | | |

This report has been created with Olex2 [6], compiled on 2022.04.07 svn.rca3783a0 for OlexSys.

### X-ray Structure Determination of Compound 2

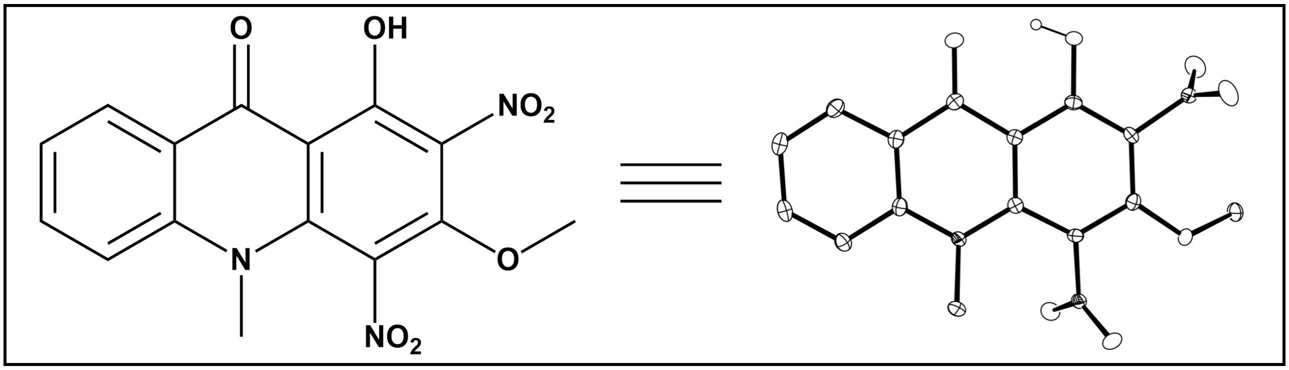


Compound **2**, $C_{15}H_{11}N_3O_7$, crystallizes in the monoclinic space group $P2_1/c$ (systematic absences 0k0: k=odd and h0l: l=odd) with a=9.4121(4)Å, b=14.5547(6)Å, c=10.3081(5)Å, α=90°, β=99.071(4)°, γ=90°, V=1394.45(11)$Å^3$, Z=4, and $d_{calc}$=1.645 g/$cm^3$. X-ray intensity data were collected on a Rigaku XtaLAB Synergy-S diffractometer [1] equipped with an HPC area detector (Dectris Pilatus3 R 200K) and employing confocal multilayer optic-monochromated Mo-Kα radiation (λ=0.71073 Å) at a temperature of 100(K. Preliminary indexing was performed from a series of thirty 0.5° rotation frames with exposures of 1.25 seconds. A total of 1194 frames (13 runs) were collected employing ω scans with a crystal to detector distance of 34.0 mm, rotation widths of 0.5° and exposures of 13.25 seconds.

Rotation frames were integrated using CrysAlisPro [2], producing a listing of unaveraged $F^2$ and $σ(F^2)$ values. A total of 27961 reflections were measured over the ranges 4.382 ≤ 2θ ≤ 56.56°, -12 ≤ h ≤ 12, -19 ≤ k ≤ 19, -13 ≤ l ≤ 13 yielding 3459 unique reflections ($R_{int}$ = 0.0588). The intensity data were corrected for Lorentz and polarization effects and for absorption using SCALE3 ABSPACK [3] (minimum and maximum transmission 0.61353, 1.00000). The structure was solved by dual space methods - SHELXT [4]. Refinement was by full-matrix least squares based on $F^2$ using SHELXL [5]. All reflections were used during refinement. The weighting scheme used was $w=1/[σ^2(F_o^2)+(0.0487P)^2+0.5769P]$ where $P = (F_o^2 + 2F_c^2)/3$. Non-hydrogen atoms were refined anisotropically and hydrogen atoms were refined using a riding model. Refinement converged to R1=0.0357 and wR2=0.0948 for 2962 observed reflections for which F > 4σ(F) and R1=0.0433 and wR2=0.0984 and GOF =1.039 for all 3459 unique, non-zero reflections and 229 variables. The maximum Δ/σ in the final cycle of least squares was 0.000 and the two most prominent peaks in the final difference Fourier were +0.41 and -0.28 e/$Å^3$.

Table 1. lists cell information, data collection parameters, and refinement data. Final positional and equivalent isotropic thermal parameters are given in Tables 2. and 3. Anisotropic thermal parameters are in Table 4. Tables 5. and 6. list bond distances and bond angles. Figure 1. is an ORTEP representation of the molecule with 50% probability thermal ellipsoids displayed.

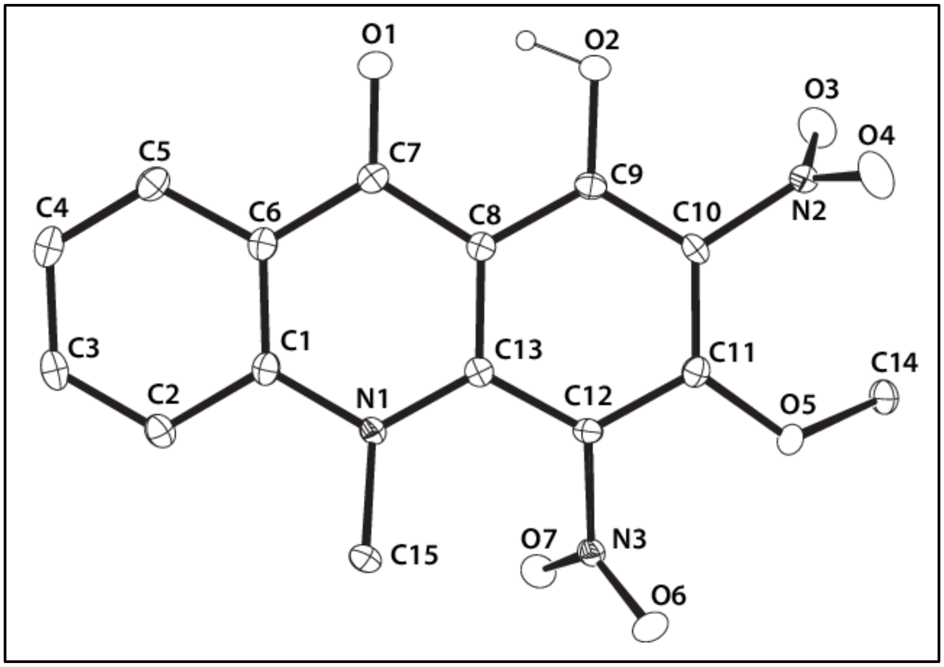


**Figure S45.** ORTEP drawing of the title compound **2** with 50% thermal ellipsoids.

**Table S28.** Summary of Structure Determination of Compound **2**

| | |
|---|---|
| Empirical formula | $C_{15}H_{11}N_3O_7$ |
| Formula weight | 345.27 |
| Diffractometer | Rigaku XtaLAB Synergy-S (Dectris Pilatus3 R 200K) |
| Temperature/K | 100(2) |
| Crystal system | monoclinic |
| Space group | $P2_1/c$ |
| a | 9.4121(4)Å |
| b | 14.5547(6)Å |
| c | 10.3081(5)Å |
| α | 90° |
| β | 99.071(4)° |
| γ | 90° |
| Volume | 1394.45(11)$Å^3$ |
| Z | 4 |
| $d_{calc}$ | 1.645 $g/cm^3$ |
| μ | 0.134 $mm^{-1}$ |
| F(000) | 712.0 |
| Crystal size, mm | 0.148 × 0.121 × 0.058 |
| 2θ range for data collection | 4.382 - 56.56° |
| Index ranges | $-12 \leq h \leq 12$, $-19 \leq k \leq 19$, $-13 \leq l \leq 13$ |
| Reflections collected | 27961 |
| Independent reflections | 3459[R(int) = 0.0588] |
| Data/restraints/parameters | 3459/0/229 |
| Goodness-of-fit on $F^2$ | 1.039 |
| Final R indexes [I>=2σ (I)] | $R_1$ = 0.0357, $wR_2$ = 0.0948 |
| Final R indexes [all data] | $R_1$ = 0.0433, $wR_2$ = 0.0984 |
| Largest diff. peak/hole | 0.41/-0.28 $eÅ^{-3}$ |

**Table S29.** Refined Positional Parameters for Compound **2.**

| Atom | *x* | *y* | *z* | U(eq) |
|---|---|---|---|---|
| O1 | 0.28892(9) | 0.62651(6) | 0.58474(8) | 0.01562(19) |
| O2 | 0.11485(9) | 0.64351(6) | 0.37915(8) | 0.01440(19) |
| O3 | -0.03180(9) | 0.59684(7) | 0.11192(10) | 0.0215(2) |
| O4 | 0.00712(10) | 0.74100(7) | 0.0791(1) | 0.0230(2) |
| O6 | 0.58249(9) | 0.68236(6) | 0.00349(8) | 0.0159(2) |
| O7 | 0.61864(9) | 0.54913(6) | 0.09872(9) | 0.0160(2) |
| O5 | 0.29516(9) | 0.66072(6) | -0.02924(8) | 0.01397(19) |
| N1 | 0.62303(10) | 0.63492(7) | 0.37252(9) | 0.0105(2) |
| N2 | 0.04674(10) | 0.66430(7) | 0.11766(10) | 0.0131(2) |
| N3 | 0.55912(10) | 0.62453(7) | 0.08430(9) | 0.0112(2) |
| C1 | 0.65068(12) | 0.61198(8) | 0.50591(11) | 0.0113(2) |
| C2 | 0.79246(13) | 0.59287(8) | 0.56609(12) | 0.0138(2) |
| C3 | 0.82043(13) | 0.57456(8) | 0.69931(12) | 0.0150(2) |
| C4 | 0.71145(14) | 0.57574(8) | 0.77700(12) | 0.0155(2) |
| C5 | 0.57145(13) | 0.59182(8) | 0.71829(12) | 0.0139(2) |
| C6 | 0.53944(12) | 0.60869(8) | 0.58216(11) | 0.0117(2) |
| C7 | 0.39191(12) | 0.62438(8) | 0.51989(11) | 0.0114(2) |
| C8 | 0.36822(12) | 0.63475(8) | 0.37896(11) | 0.0106(2) |
| C9 | 0.22463(12) | 0.64253(8) | 0.31258(11) | 0.0111(2) |
| C10 | 0.19798(12) | 0.65283(8) | 0.17694(12) | 0.0114(2) |
| C11 | 0.30979(12) | 0.65403(8) | 0.10240(11) | 0.0110(2) |
| C12 | 0.45104(12) | 0.64499(8) | 0.16862(11) | 0.0106(2) |
| C14 | 0.17003(13) | 0.62357(9) | -0.11303(12) | 0.0151(2) |
| C15 | 0.74413(12) | 0.67173(9) | 0.31328(12) | 0.0148(2) |
| C13 | 0.48427(12) | 0.63785(8) | 0.30693(11) | 0.0100(2) |

**Table S30.** Positional Parameters for Hydrogens in Compound **2.**

| Atom | *x* | *y* | *z* | U(eq) |
|---|---|---|---|---|
| H2 | 0.145976 | 0.639386 | 0.459914 | 0.022 |
| H2A | 0.868504 | 0.592549 | 0.515453 | 0.017 |
| H3 | 0.916126 | 0.560839 | 0.739018 | 0.018 |
| H4 | 0.733425 | 0.565614 | 0.869025 | 0.019 |
| H5 | 0.496285 | 0.591502 | 0.769919 | 0.017 |
| H14A | 0.193622 | 0.611935 | -0.200825 | 0.023 |
| H14B | 0.0907 | 0.667778 | -0.119342 | 0.023 |
| H14C | 0.14124 | 0.565905 | -0.07552 | 0.023 |
| H15A | 0.79433 | 0.621055 | 0.277277 | 0.022 |
| H15B | 0.811107 | 0.703941 | 0.380575 | 0.022 |
| H15C | 0.707355 | 0.71465 | 0.242666 | 0.022 |

**Table S31.** Refined Thermal Parameters (U's) for Compound **2**

| Atom | $U_{11}$ | $U_{22}$ | $U_{33}$ | $U_{23}$ | $U_{13}$ | $U_{12}$ |
|---|---|---|---|---|---|---|
| O1 | 0.0134(4) | 0.0217(5) | 0.0127(4) | 0.0004(3) | 0.0049(3) | 0.0016(3) |
| O2 | 0.0102(4) | 0.0218(5) | 0.0118(4) | 0.0000(3) | 0.0038(3) | 0.0013(3) |
| O3 | 0.0143(4) | 0.0237(5) | 0.0254(5) | 0.0016(4) | -0.0006(4) | -0.0070(4) |
| O4 | 0.0172(4) | 0.0207(5) | 0.0294(5) | 0.0054(4) | -0.0019(4) | 0.0061(4) |
| O6 | 0.0178(4) | 0.0176(4) | 0.0131(4) | 0.0030(3) | 0.0054(3) | -0.0019(3) |
| O7 | 0.0152(4) | 0.0144(4) | 0.0189(5) | -0.0006(3) | 0.0041(3) | 0.0035(3) |
| O5 | 0.0122(4) | 0.0205(4) | 0.0085(4) | 0.0013(3) | -0.0005(3) | -0.0029(3) |
| N1 | 0.0082(4) | 0.0127(5) | 0.0104(5) | -0.0007(4) | 0.0008(3) | -0.0010(4) |
| N2 | 0.0104(5) | 0.0180(5) | 0.0110(5) | 0.0008(4) | 0.0019(4) | 0.0006(4) |
| N3 | 0.0094(4) | 0.0137(5) | 0.0103(5) | -0.0014(4) | 0.0010(4) | -0.0013(4) |
| C1 | 0.0137(5) | 0.0090(5) | 0.0106(5) | -0.0013(4) | -0.0002(4) | -0.0012(4) |
| C2 | 0.0117(5) | 0.0135(6) | 0.0156(6) | 0.0000(4) | 0.0003(4) | -0.0013(4) |
| C3 | 0.0138(5) | 0.0129(6) | 0.0163(6) | -0.0008(4) | -0.0040(4) | -0.0003(4) |
| C4 | 0.0208(6) | 0.0132(6) | 0.0112(5) | 0.0002(4) | -0.0017(5) | -0.0002(5) |
| C5 | 0.0182(6) | 0.0121(6) | 0.0117(6) | -0.0004(4) | 0.0029(4) | -0.0002(4) |
| C6 | 0.0137(5) | 0.0098(5) | 0.0113(5) | -0.0009(4) | 0.0008(4) | 0.0001(4) |
| C7 | 0.0137(5) | 0.0095(5) | 0.0114(5) | -0.0008(4) | 0.0027(4) | -0.0002(4) |
| C8 | 0.0109(5) | 0.0098(5) | 0.0110(5) | -0.0005(4) | 0.0018(4) | -0.0001(4) |
| C9 | 0.0109(5) | 0.0097(5) | 0.0131(5) | -0.0008(4) | 0.0036(4) | 0.0000(4) |
| C10 | 0.0077(5) | 0.0124(5) | 0.0137(6) | 0.0002(4) | 0.0000(4) | 0.0007(4) |
| C11 | 0.0131(5) | 0.0091(5) | 0.0105(5) | 0.0006(4) | 0.0010(4) | -0.0007(4) |
| C12 | 0.0093(5) | 0.0117(5) | 0.0114(5) | -0.0004(4) | 0.0033(4) | -0.0004(4) |
| C14 | 0.0130(6) | 0.0195(6) | 0.0116(5) | -0.0025(5) | -0.0018(4) | -0.0020(5) |
| C15 | 0.0093(5) | 0.0211(6) | 0.0142(6) | 0.0003(5) | 0.0022(4) | -0.0036(4) |
| C13 | 0.0109(5) | 0.0075(5) | 0.0115(5) | -0.0007(4) | 0.0012(4) | -0.0005(4) |

**Table S32.** Bond Distances in Compound **2,** Å

| | | | | | |
|---|---|---|---|---|---|
| O1-C7 | 1.2616(14) | O2-C9 | 1.3275(14) | O3-N2 | 1.2247(14) |
| O4-N2 | 1.2234(14) | O6-N3 | 1.2282(13) | O7-N3 | 1.2304(13) |
| O5-C11 | 1.3457(14) | O5-C14 | 1.4507(14) | N1-C1 | 1.3989(15) |
| N1-C15 | 1.4760(15) | N1-C13 | 1.3734(14) | N2-C10 | 1.4674(14) |
| N3-C12 | 1.4689(14) | C1-C2 | 1.4079(16) | C1-C6 | 1.4056(16) |
| C2-C3 | 1.3828(17) | C3-C4 | 1.3973(18) | C4-C5 | 1.3805(17) |
| C5-C6 | 1.4096(16) | C6-C7 | 1.4530(16) | C7-C8 | 1.4425(16) |
| C8-C9 | 1.4198(16) | C8-C13 | 1.4149(15) | C9-C10 | 1.3891(16) |
| C10-C11 | 1.3976(16) | C11-C12 | 1.4017(16) | C12-C13 | 1.4145(16) |

**Table S33.** Bond Angles in Compound **2**, °

| | | | | | |
|---|---|---|---|---|---|
| C11-O5-C14 | 121.18(9) | C1-N1-C15 | 117.21(9) | C13-N1-C1 | 120.42(9) |
| C13-N1-C15 | 121.43(10) | O3-N2-C10 | 117.87(10) | O4-N2-O3 | 124.41(10) |
| O4-N2-C10 | 117.71(10) | O6-N3-O7 | 124.49(10) | O6-N3-C12 | 118.50(9) |
| O7-N3-C12 | 117.00(9) | N1-C1-C2 | 119.78(10) | N1-C1-C6 | 121.21(10) |
| C6-C1-C2 | 119.00(11) | C3-C2-C1 | 119.63(11) | C2-C3-C4 | 121.66(11) |
| C5-C4-C3 | 119.14(11) | C4-C5-C6 | 120.36(11) | C1-C6-C5 | 120.09(11) |
| C1-C6-C7 | 119.46(10) | C5-C6-C7 | 120.44(11) | O1-C7-C6 | 122.02(11) |
| O1-C7-C8 | 121.43(11) | C8-C7-C6 | 116.52(10) | C9-C8-C7 | 118.55(10) |
| C13-C8-C7 | 121.48(10) | C13-C8-C9 | 119.97(10) | O2-C9-C8 | 120.74(10) |
| O2-C9-C10 | 119.17(10) | C10-C9-C8 | 120.05(10) | C9-C10-N2 | 116.16(10) |
| C9-C10-C11 | 121.51(10) | C11-C10-N2 | 122.32(10) | O5-C11-C10 | 126.09(10) |
| O5-C11-C12 | 115.87(10) | C10-C11-C12 | 118.02(10) | C11-C12-N3 | 115.03(10) |
| C11-C12-C13 | 122.67(10) | C13-C12-N3 | 121.19(10) | N1-C13-C8 | 119.58(10) |
| N1-C13-C12 | 122.71(10) | C12-C13-C8 | 117.71(10) | | |

This report has been created with Olex2 [6], compiled on 2022.04.07 svn.rca3783a0 for OlexSys.

**Synthetase Sequences**
**pULTRAI-Tet3.0 RS (RS sequence underlined).**
accaccctgaattgactctcttccgggcgctatcatgccataccgcgaaaggttttgcgccattcgatggtgtccgggatctcgacgctct
cccttatgcgactcctgcattagggagctgttgacaattaatcatcggctcgtataatgtgtggaattgtgagcggataacaatttcacaa
aggaggtgcggccgcatggataaaaaaccgctggatgtgctgattagcgcgaccggcctgtggatgagccgtaccggcaccctgc
ataaaatcaaacatcatgaagtgagccgcagcaaaatctatattgaaatggcgtgcggcgatcatctggtggtgaacaacagccgta
gctgccgtaccgcgcgtgcgtttcgtcatcataaataccgcaaaacctgcaaacgttgccgtgtgagcgatgaagatatcaacaacttt
ctgacccgtagcaccgaaagcaaaaacagcgtgaaagtgcgtgtggtgagcgcgccgaaagtgaaaaaagcgatgccgaaaa
gcgtgagccgtgcgccgaaaccgctggaaaatagcgtgagcgcgaaagcgagcaccaacaccagccgtagcgttccgagcccg
gcgaaaagcaccccgaacagcagcgttccggcgtctgcgccggcaccgagcctgacccgcagccagctggatcgtgtggaagcg
ctgctgtctccggaagataaaattagcctgaacatggcgaaaccgtttcgtgaactggaaccggaactggtgacccgtcgtaaaaac
gattttcagcgcctgtataccaacgatcgtgaagattatctgggcaaactggaacgtgatatcaccaaatttttgtggatcgcggctttct
ggaaattaaaagcccgattctgattccggcggaatatgtggaacgtatgggcattaacaacgacaccgaactgagcaaacaaatttt
ccgtgtggataaaaacctgtgcctgcgtccgatgctggccccgaccggttataactatttgcgtaaactggatcgtattctgccgggtccg
atcaaaatttttgaagtgggcccgtgctatcgcaaagaaagcgatggcaaagaacacctggaagaattcaccatggttggttttgctca
aatgggcagcggctgcacccgtgaaaacctggaagcgctgatcaaagaattcctggattatctggaaatcgacttcgaaattgtggg
cgatagctgcatggtgtatggcgataccctggatattatgcatggcgatctggaactgagcagcgcggtggtgggtccggttagcctgg
atcgtgaatggggcattgataaaccgtggattggcgcgggttttggcctggaacgtctgctgaaagtgatgcatggcttcaaaaacatta
aacgtgcgagccgtagcgaaagctactataacggcattagcacgaacctgtaagcggccgcgtttaaacggtctccagcttggctgtt
ttggcggatgagagaagattttcagcctgatacagattaaatcagaacgcagaagcggtctgataaaacagaatttgcctggcggca
gtagcgcggtggtcccacctgaccccatgccgaactcagaagtgaaacgccgtagcgccgatggtagtgtggggtctccccatgcg
agagtagggaactgccaggcatcaaataaaacgaaaggctcagtcgaaagactgggccttgtttgtgagctcccggtcatcaatcat
ccccataatccttgttagcctgcaggtaattccgcttcgcaacatgtgagcaccggtttattgactaccggaagcagtgtgaccgtgtgctt
ctcaaatgcctgaggccagtttgctcaggctctccccgtggaggtaataattgacgatatgatcagtgcacggctaactaagcggcctg
ctgactttctcgccgatcaaaaggcattttgctattaagggattgacgagggcgtatctgcgcagtaagatgcgccccgcattcggaaa
cctgatcatgtagatcgaacggactttaaatccgttcagccgggttagattcccggggtttccgccaaattcgaaaagcctgctcaacg
agcaggcttttttgcatgctcgagcagctcagggtcgaatttgggtaccaccggcgcctcaggcatttgagaagcacacggtcacact
gcttccggtagtcaataaaccggtaaaccagcaatagacataagcggctatttaacgaccctgccctgaaccgacgaccgggtcatc
gtggccggatcttgcggcccctcggcttgaacgaattgttagacattatttgccgactaccttggtgatctcgcctttcacgtagtggacaa
attcttccaactgatctgcgcgcgaggccaagcgatcttcttcttgtccaagataagcctgtctagcttcaagtatgacgggctgatactg
ggccggcaggcgctccattgcccagtcggcagcgacatccttcggcgcgattttgccggttactgcgctgtaccaaatgcgggacaa
cgtaagcactacatttcgctcatcgccagcccagtcgggcggcgagttccatagcgttaaggtttcatttagcgcctcaaatagatcctgt
tcaggaaccggatcaaagagttcctccgccgctggacctaccaaggcaacgctatgttctcttgcttttgtcagcaagatagccagatc
aatgtcgatcgtggctggctcgaagatacctgcaagaatgtcattgcgctgccattctccaaattgcagttcgcgcttagctggataacg
ccacggaatgatgtcgtcgtgcacaacaatggtgacttctacagcgcggagaatctcgctctctccaggggaagccgaagtttccaa
aaggtcgttgatcaaagctcgccgcgttgtttcatcaagccttacggtcaccgtaaccagcaaatcaatatcactgtgtggcttcaggcc
gccatccactgcggagccgtacaaatgtacggccagcaacgtcggttcgagatggcgctcgatgacgccaactacctctgatagttg
agtcgatacttcggcgatcaccgcttccctcatactcttcctttttcaatattattgaagcatttatcagggttattgtctcatgagcggatacat
atttgaatgtatttagaaaaataaacaaatagctagctcactcggtcgctacgctccgggcgtgagactgcggcgggcgctgcggaca
catacaaagttacccacagattccgtggataagcaggggactaacatgtgaggcaaaacagcagggccgcgccggtggcgtttttc
cataggctccgcccctcctgccagagttcacataaacagacgcttttccggtgcatctgtgggagccgtgaggctcaaccatgaatctga
cagtacgggcgaaacccgacaggacttaaagatccccaccgtttccggcgggtcgctccctcttgcgctctcctgttccgaccctgccg
tttaccggatacctgttccgcctttctcccttacgggaagtgtggcgctttctcatagctcacacactggtatctcggctcggtgtaggtcgtt
cgctccaagctgggctgtaagcaagaactccccgttcagcccgactgctgcgccttatccggtaactgttcacttgagtccaacccgga

aaagcacggtaaaacgccactggcagcagccattggtaactgggagttcgcagaggatttgtttagctaaacacgcggttgctcttga
agtgtgcgccaaagtccggctacactggaaggacagatttggttgctgtgctctgcgaaagccagttaccacggttaagcagttcccc
aactgacttaaccttcgatcaaaccacctccccaggtggtttttcgtttacagggcaaaagattacgcgcagaaaaaaaggatctcaa
gaagatcctttgatcttttctactgaaccgctctagatttcagtgcaatttatctcttcaaatgtagcacctgaagtcagccccatacgatata
agttgtaattctcatgttagtcatgccccgcgcccaccggaaggagctgactgggttgaaggctctcaagggcatcggtcgagatccc
ggtgcctaatgagtgagctaacttacattaattgcgttgcgctcactgcccgctttccagtcgggaaacctgtcgtgccagctgcattaat
gaatcggccaacgcgcggggagaggcggtttgcgtattgggcgccagggtggtttttcttttcaccagtgagacgggcaacagctgat
tgcccttcaccgcctggccctgagagagttgcagcaagcggtccacgctggtttgccccagcaggcgaaaatcctgtttgatggtggtt
aacggcgggatataacatgagctgtcttcggtatcgtcgtatcccactaccgagatgtccgcaccaacgcgcagcccggactcggta
atggcgcgcattgcgcccagcgccatctgatcgttggcaaccagcatcgcagtgggaacgatgccctcattcagcatttgcatggtttgt
tgaaaaccggacatggcactccagtcgccttcccgttccgctatcggctgaatttgattgcgagtgagatatttatgccagccagccaga
cgcagacgcgccgagacagaacttaatgggcccgctaacagcgcgatttgctggtgacccaatgcgaccagatgctccacgccca
gtcgcgtaccgtcttcatgggagaaaataatactgttgatgggtgtctggtcagagacatcaagaaataacgccggaacattagtgca
ggcagcttccacagcaatggcatcctggtcatccagcggatagttaatgatcagcccactgacgcgttgcgcgagaagattgtgcacc
gccgctttacaggcttcgacgccgcttcgttctaccatcgacaccaccacgctggcacccagttgatcggcgcgagatttaatcgccgc
gacaatttgcgacggcgcgtgcagggccagactggaggtggcaacgccaatcagcaacgactgtttgcccgccagttgttgtgccac
gcggttgggaatgtaattcagctccgccatcgccgcttccactttttcccgcgttttcgcagaaacgtggctggcctggttcaccacgcgg
gaaacggtctgataagagacaccggcatactctgcgacatcgtataacgttactggtttcacattc

**pEVOL-AcdA9 RS (RS sequence underlined).**
aagaaaccaattgtccatattgcatcagacattgccgtcactgcgtcttttactggctcttctcgctaaccaaaccggtaaccccgcttatt
aaaagcattctgtaacaaagcgggaccaaagccatgacaaaaacgcgtaacaaaagtgtctataatcacggcagaaaagtccac
attgattatttgcacggcgtcacactttgctatgccatagcatttttatccataagattagcggatcctacctgacgctttttatcgcaactctct
actgtttctccatacccgtttttttgggctaacaggaggaattagatctatggacgaatttgaaatgataaagagaaacacatctgaaatt
atcagcgaggaagagttaagagaggttttaaaaaaagatgaaaaatctgctgcgataggttttgaaccaagtggtaaaatacatttag
ggcattatctccaaataaaaaagatgattgatttacaaaatgctggatttgatataattatagatttggctgatttacacgcctatttaaacc
agaaaggagagttggatgagattagaaaaataggagattataacaaaaaagtttttgaagcaatggggttaaaggcaaaatatgttt
atggaagtgaactggagcttgataaggattatacactgaatgtctatagattggctttaaaaactaccttaaaaagagcaagaaggagt
atggaacttatagcaagagaggatgaaaatccaaaggttgctgaagttatctatccaataatgcaggttaattctattcattatactggcg
ttgatgttgcagttggagggatggagcagagaaaaataaacatgttagcaagggagcttttaccaaaaaaggttgtttgtattcacaac
cctgtcttaacgggtttggatggagaaggaaagatgagttcttcaaaagggaattttatagctgttgatgactctccagaagagattagg
gctaagataaagaaagcatactgcccagctggagttgttgaaggaaatccaataatggagatagctaaatacttccttgaatatccttta
accataaaaaggccagaaaaatttggtggagatttgacagttaatagctatgaggagttagagagtttatttaaaaataaggaattgca
tccaatggatttaaaaaatgctgtagctgaagaacttataaagattttagagccaattagaaagagattatgatgagtttaaacggtctcc
agcttggctgttttggcggatgagagaagattttcagcctgatacagattaaatcagaacgcagaagcggtctgataaaacagaatttg
cctggcggcagtagcgcggtggtcccacctgaccccatgccgaactcagaagtgaaacgccgtagcgccgatggtagtgtggggtc
tccccatgcgagagtagggaactgccaggcatcaaataaaacgaaaggctcagtcgaaagactgggccttgtttgtgagctcccggt
catcaatcatccccataatccttgttagattatcaattttaaaaaactaacagttgtcagcctgtcccgctttaatatcatacgccgttatacg
ttgtttacgctttgaggaatcccatatggacgaatttgaaatgataaagagaaacacatctgaaattatcagcgaggaagagttaagag
aggttttaaaaaaagatgaaaaatctgctgcgataggttttgaaccaagtggtaaaatacatttagggcattatctccaaataaaaaag
atgattgatttacaaaatgctggatttgatataattatagatttggctgatttacacgcctatttaaaccagaaaggagagttggatgagatt
agaaaaataggagattataacaaaaaagtttttgaagcaatggggttaaaggcaaaatatgtttatggaagtgaactggagcttgata
aggattatacactgaatgtctatagattggctttaaaaactaccttaaaaagagcaagaaggagtatggaacttatagcaagagagga
tgaaaatccaaaggttgctgaagttatctatccaataatgcaggttaattctattcattatactggcgttgatgttgcagttggagggatgga

gcagagaaaaataaacatgttagcaagggagcttttaccaaaaaaggttgtttgtattcacaaccctgtcttaacgggtttggatggag
aaggaaagatgagttcttcaaaagggaattttatagctgttgatgactctccagaagagattagggctaagataaagaaagcatactg
cccagctggagttgttgaaggaaatccaataatggagatagctaaatacttccttgaatatcctttaaccataaaaaggccagaaaaat
ttggtggagatttgacagttaatagctatgaggagttagagagtttatttaaaaataaggaattgcatccaatggatttaaaaaatgctgta
gctgaagaacttataaagattttagagccaattagaaagagattatgactgcagtttcaaacgctaaattgcctgatgcgctacgcttatc
aggcctacatgatctctgcaatatattgagtttgcgtgcttttgtaggccggataaggcgttcacgccgcatccggcaagaaacagcaa
acaatccaaaacgccgcgttcagcggcgtttttctgcttttcttcgcgaattaattccgcttcgcaacatgtgagcaccggtttattgactac
cggaagcagtgtgaccgtgtgcttctcaaatgcctgaggccagtttgctcaggctctccccgtggaggtaataattgacgatatgatcag
tgcacggctaactaagcggcctgctgactttctcgccgatcaaaaggcattttgctattaagggattgacgagggcgtatctgcgcagta
agatgcgccccgcattccggcggtagttcagcagggcagaacggcggactctaaatccgcatggcaggggttcaaatcccctccgc
cggaccaaattcgaaaagcctgctcaacgagcaggctttttgcatgctcgagcagctcagggtcgaatttgctttcgaatttctgccatt
catccgcttattatcacttattcaggcgtagcaaccaggcgtttaagggcaccaataactgccttaaaaaaattacgccccgccctgcc
actcatcgcagtactgttgtaattcattaagcattctgccgacatggaagccatcacaaacggcatgatgaacctgaatcgccagcgg
catcagcaccttgtcgccttgcgtataatatttgcccatggtgaaaacgggggcgaagaagttgtccatattggccacgtttaaatcaaa
actggtgaaactcacccagggattggctgagacgaaaaacatattctcaataaaccctttagggaaataggccaggttttcaccgtaa
cacgccacatcttgcgaatatatgtgtagaaactgccggaaatcgtcgtggtattcactccagagcgatgaaaacgtttcagtttgctcat
ggaaaacggtgtaacaagggtgaacactatcccatatcaccagctcaccgtctttcattgccatacggaattccggatgagcattcatc
aggcgggcaagaatgtgaataaaggccggataaaacttgtgcttatttttctttacggtctttaaaaaggccgtaatatccagctgaacg
gtctggttataggtacattgagcaactgactgaaatgcctcaaaatgttctttacgatgccattgggatatatcaacggtggtatatccagt
gatttttttctccattttagcttccttagctcctgaaaatctcgataactcaaaaaatacgcccggtagtgatcttatttcattatggtgaaagtt
ggaacctcttacgtgccgatcaacgtctcattttcgccaaaagttggcccagggcttcccggtatcaacagggacaccaggatttattt at
tctgcgaagtgatcttccgtcacaggtatttattcggcgcaaagtgcgtcgggtgatgctgccaacttactgatttagtgtatgatggtgttttt
gaggtgctccagtggcttctgtttctatcagctgtccctcctgttcagctactgacggggtggtgcgtaacggcaaaagcaccgccggac
atcagcgctagcggagtgtatactggcttactatgttggcactgatgagggtgtcagtgaagtgcttcatgtggcaggagaaaaaagg
ctgcaccggtgcgtcagcagaatatgtgatacaggatatattccgcttcctcgctcactgactcgctacgctcggtcgttcgactgcggc
gagcggaaatggcttacgaacggggcggagatttcctggaagatgccaggaagatacttaacagggaagtgagagggccgcggc
aaagccgtttttccataggctccgcccccctgacaagcatcacgaaatctgacgctcaaatcagtggtggcgaaacccgacaggact
ataaagataccaggcgtttccccctggcggctccctcgtgcgctctcctgttcctgcctttcggtttaccggtgtcattccgctgttatggccg
cgtttgtctcattccacgcctgacactcagttccgggtaggcagttcgctccaagctggactgtatgcacgaaccccccgttcagtccga
ccgctgcgccttatccggtaactatcgtcttgagtccaacccggaaagacatgcaaaagcaccactggcagcagccactggtaattg
atttagaggagttagtcttgaagtcatgcgccggttaaggctaaactgaaaggacaagttttggtgactgcgctcctccaagccagttac
ctcggttcaaagagttggtagctcagagaaccttcgaaaaaccgccctgcaaggcggttttttcgttttcagagcaagagattacgcgc
agaccaaaacgatctcaagaagatcatcttattaatcagataaaatatttctagatttcagtgcaatttatctcttcaaatgtagcacctga
agtcagccccatacgatataagttgtaattctcatgtttgacagcttatcatcgataagcttggtacccaattatgacaacttgacggctac
atcattcactttttcttcacaaccggcacggaactcgctcgggctggccccggtgcattttttaaatacccgcgagaaatagagttgatcgt
caaaaccaacattgcgaccgacggtggcgataggcatccgggtggtgctcaaaagcagcttcgcctggctgatacgttggtcctcgc
gccagcttaagacgctaatccctaactgctggcggaaaagatgtgacagacgcgacggcgacaagcaaacatgctgtgcgacgct
ggcgatatcaaaattgctgtctgccaggtgatcgctgatgtactgacaagcctcgcgtacccgattatccatcggtggatggagcgact
cgttaatcgcttccatgcgccgcagtaacaattgctcaagcagatttatcgccagcagctccgaatagcgcccttccccttgcccggcgt
taatgatttgcccaaacaggtcgctgaaatgcggctggtgcgcttcatccgggcgaaagaaccccgtattggcaaatattgacggcca
gttaagccattcatgccagtaggcgcgcggacgaaagtaaacccactggtgataccattcgcgagcctccggatgacgaccgtagt
gatgaatctctcctggcgggaacagcaaaatatcactcggtcggcaaacaaattctcgtccctgatttttcaccaccccctgaccgcga
atggtgagattgagaatataacctttcattcccagcggtcggtcgataaaaaaatcgagataaccgttggcctcaatcggcgttaaacc
cgccaccagatgggcattaaacgagtatcccggcagcaggggatcattttgcgcttcagccatacttttcatactcccgccattcagag
